# Sirenian genomes illuminate the evolution of fully aquatic species within the mammalian superorder Afrotheria

**DOI:** 10.1101/2023.09.01.555811

**Authors:** Ran Tian, Yaolei Zhang, Hui Kang, Fan Zhang, Zhihong Jin, Jiahao Wang, Peijun Zhang, Xuming Zhou, Janet M. Lanyon, Helen L. Sneath, Lucy Woolford, Guangyi Fan, Songhai Li, Inge Seim

**Author notes:** These authors contributed equally: Ran Tian, Yaolei Zhang, Hui Kang. Correspondence (G.F.), (S.L.), and (I.S.).

## Abstract

Sirenians of the diverse superorder Afrotheria were the first mammals to retransition from land to water and are the sole herbivorous marine mammals, but little is known about the underlying genomic changes. Here, we generated a chromosome-level dugong (*Dugong dugon*) genome. A comparison of our assembly and a recent West Indian manatee genome with other afrotherian genomes revealed possible molecular solutions to aquatic life by sirenians, including a shift in daily activity patterns (circadian clock) and tolerance to a high-iodine plant diet mediated through changes in the iodide transporter NIS (*SLC5A5*) and its co-transporters. Functional *in vitro* assays confirmed that sirenian amino acid substitutions alter the properties of the circadian clock protein PER2 and NIS. Sirenians show evidence of convergent regression of integumentary system (skin and its appendages) genes with cetaceans, the only other fully aquatic mammal group. Our analysis also uncovered gene losses that may be maladaptive in a modern environment, including a candidate gene (*KCNK18*) for sirenian cold stress syndrome likely lost during their evolutionary shift in daily activity patterns. Finally, genomes from nine Australian locations and the functionally extinct Okinawan population confirm and date a genetic break on the Australian east coast and highlight the need for whole-genome resequencing data from small or isolated dugong populations worldwide for conservation and genetic management.

## Main

The terrestrial ancestors of the marine mammal groups Sirenia, Cetacea, and Pinnipedia independently transitioned from land to water ^1^. The first to leave land, sirenians, emerged around 60 million years ago within the afrotherian herbivorous clade Paenungulata, 10 and 30 million years before the emergence of cetaceans and pinnipeds ^2^. Afrotherian mammals were isolated from other mammals until ∼60 Mya when non-afrotherian mammals (ungulates from ∼25 Mya) began to enter the African continent from Eurasia and displaced many local species ^3,4^. This geographic isolation allowed the independent evolution of terrestrial mammals to an aquatic habitat in Africa (sirenians), paralleling the evolution of fully aquatic cetaceans from ungulates.

Dozens of sirenian species have existed in the past, but unlike cetaceans (about 90 extant species) and pinnipeds (about 30 extant species), sirenians are today far less diverse (**Figure 1a**). There are four extant sirenian species: the dugong (*Dugong dugon*) of the family Dugongidae (included the Steller’s sea cow, *Hydrodamalis gigas*, that became extinct about 250 years ago) and manatees (family Trichechidae: the West Indian manatee, *Trichechus manatus* (includes the subspecies Florida manatee, *T. m. latirostris*, and Antillean manatee, *T. m. manatus*); the Amazonian manatee, *T. inunguis*; and the African manatee, *T. senegalensis*) ^1,5^. The dugong and manatees have been found in distinct tropical and subtropical habitats since the middle Miocene (∼12.2 Mya) ^2,5^. The dugong originally dispersed into the Pacific from near Florida, and today inhabits the coastlines of the Indo-Pacific oceans, while the three species of manatee occupy the Atlantic Ocean and associated rivers (**Figure 1c**).

**Figure 1.**
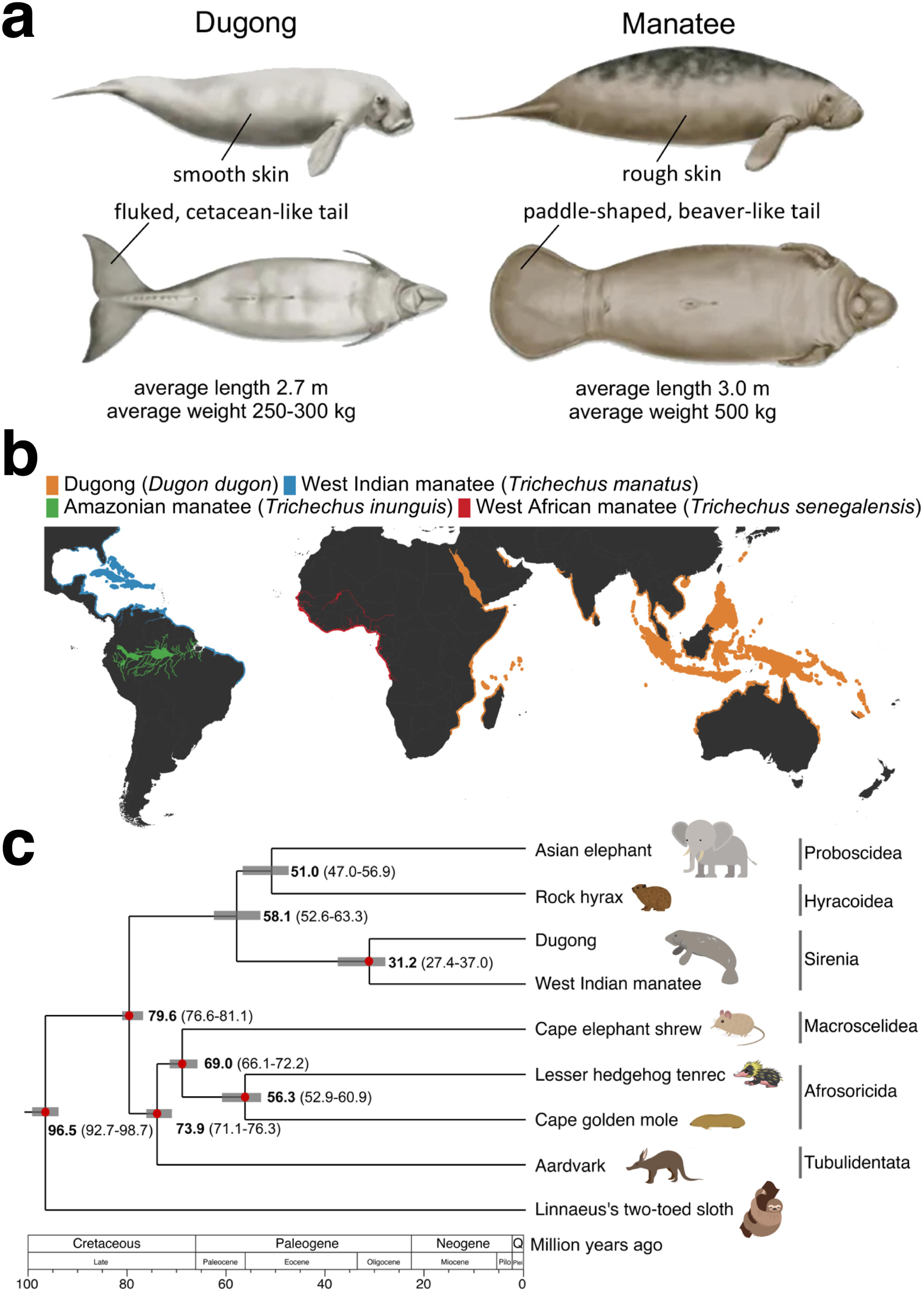
Overview of extant sirenian taxa. **a,** Features of extant sirenians. In addition to common sirenian features, the dugong (*Dugong dugon*) and manatees (genus *Trichechus*; here the West Indian manatee, *T. manatus*) show various morphological differences. By courtesy of Encyclopædia Britannica, Inc. **b,** Species distribution of sirenians. Species ranges ranges derived from spatial data on marine mammals from the International Union for the Conservation of Nature Red List of Threatened Species, December 2022 [version 6.3]. https://www.iucnredlist.org; Downloaded on 12 January 2023. Note that the dugong was very recently (2022) declared extinct in Chinese and Japanese waters. **c,** Maximum likelihood (ML) phylogenetic tree from the coding sequences of 7,695 orthologs from eight afrotherian species, rooted with a xenarthran outgroup (the Linnaeus’s two-toed sloth). 1,000 bootstraps were used, and all nodes have 100% support. The number in each node represents the divergence time among species a red circle indicates the fossil record used for calibration in the node. The gray bars on the nodes indicate the 95% credibility intervals of the estimated posterior distributions of the divergence times. Afrotherian orders are shown to the right of each representative species.

Many genome-scale studies have considered the evolution of marine mammals. However, compared to cetaceans and pinnipeds ^6,7^, the lack of high-quality genomes has limited insights into sirenians, the only fully aquatic mammals within Afrotheria. To better understand the evolution to aquatic life by sirenians, we generated and juxtaposed a chromosome-level genome and gene set of the dugong with other afrotherian mammals (including a manatee). Our genome also enabled insights into the diversity and demography of dugongs, including the genomic impact of severe population declines on a recently functionally extinct Okinawan population.

## Results

### Molecular adaptations of sirenians

We used synthetic long-read (stLFR) and chromosome conformation capture (Hi-C) sequencing to generate a 3.06 Gb 25-chromosome genome assembly (Ddugon_BGI) of a female dugong (**Supplementary Note 1**, **Figure S1**, and **Tables S1** and **S2**). The assembly and gene set BUSCO ^8^ completeness scores (94.4% and 91.7%, respectively) were comparable to seven other afrotherian species (**Tables S3** and **S4**).

Although similar in appearance, the dugong and manatees are not that closely related. They share a common ancestor (crown Sirenia) 31.2 Mya (95% CI: 27.4-37.0 Mya) (**Figure 1c**), providing a suitable backdrop to illuminate features present in the first fully aquatic sirenians. Our afrotherian dataset, which includes the phylogenetically closest extant species to sirenians (elephants and hyraxes) (**Supplementary Note 2**, **Figure 1c**, and **Figure S2**), was interrogated to identify protein-coding gene changes possibly associated with the evolution of all sirenians rather than focusing on their differences (e.g., see **Figure 1a**). Our analysis included gene family dynamics (**Tables S5** and **S6**, and **Figure S3**), positively selected genes (PSGs) (**Table S7**), rapidly evolving genes (REGs) (**Table S8**), gene loss (pseudogenes) (**Tables S9** and **Tables S10**), and amino acid substitutions (**Table S11**) shared by sirenians. Amino acid substitutions of interest were validated against 120 mammalian species in OrthoMaM ^9^ and by BLAST searches of NCBI and Ensembl databases, while pseudogenes were validated by interrogating genome and transcriptome sequencing reads. This effort identified gene changes that may contribute to aquatic herbivory, sirenian circadian activity patterns, and changes that may underly typical marine mammal features, such as modified cardiovascular, integumental (i.e., skin and associated structures), and sensory (vision, smell, and taste) systems (**Figure 2**). We also report historical gene loss events that may have become maladaptive in modern environments.

**Figure 2.**
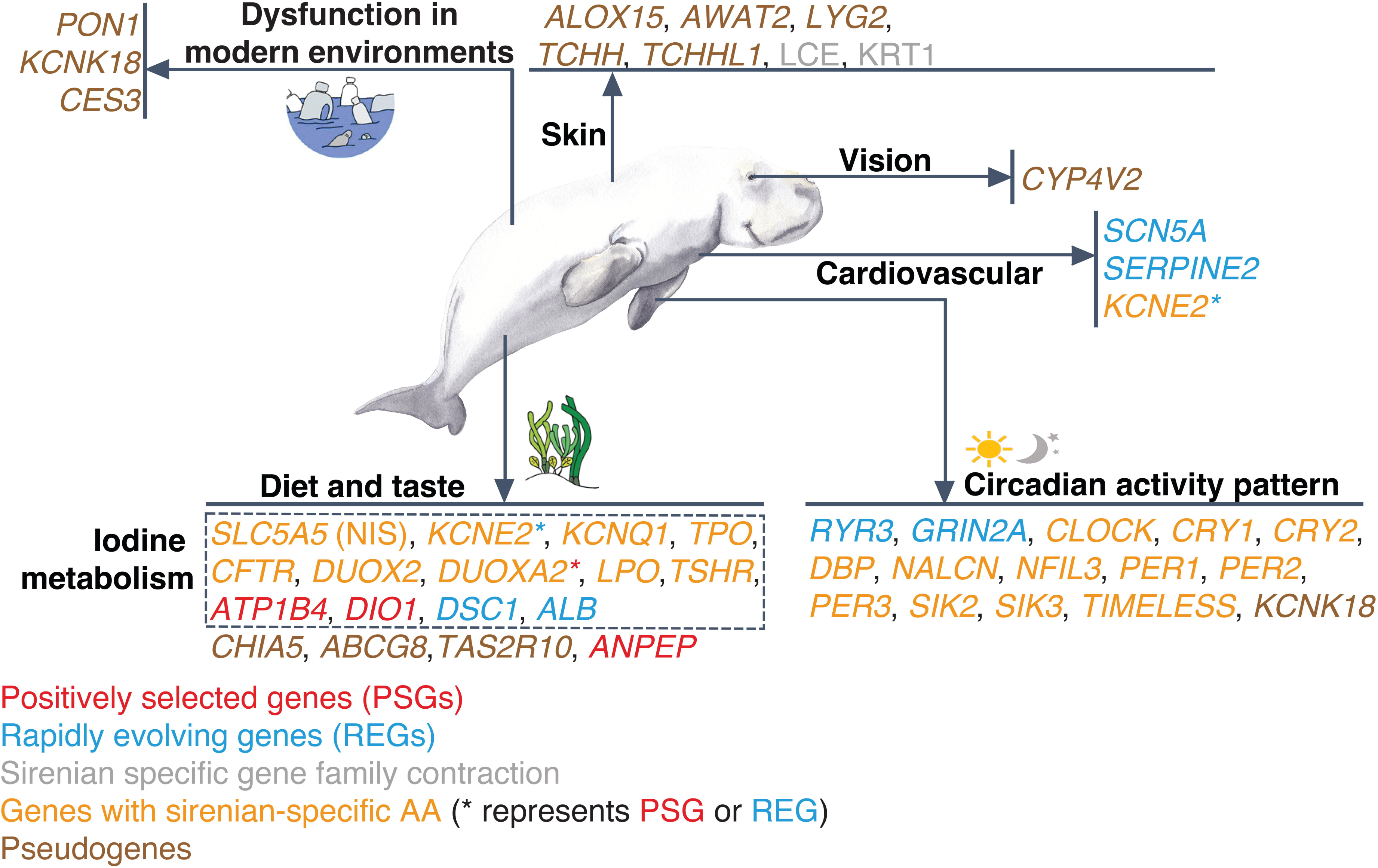
Genes with unique evolutionary signals in sirenians. Putative adaptive or maladaptive gene changes are shown.

#### Nutrient uptake of fully aquatic herbivores

Sirenians are the only aquatic herbivorous mammals, hence they have a common and misleading name, sea cows. Although sirenians can occasionally consume invertebrates, they rely primarily on aquatic plants, including marine seagrasses whose terrestrial ancestors independently returned to the sea during the Cretaceous period, from ∼66 to ∼100 Mya ^5,10^. Whilst dugongs are obligate, specialist seagrass grazers, all three species of manatee are more catholic feeders, taking a range of aquatic and semi-aquatic plant species, and in some cases, may consume emergent grassland species ^5^. Adult sirenians consume 10-15% of their body weight daily ^5,11^. Similar to the distantly related elephants and hyraxes and some more distantly related mammals (e.g., rhinoceroses), sirenians are hindgut fermenters ^5^. Manatees and the dugong have specialized mouthparts ^12,13^, long and capacious digestive tracts, and prolonged food retention times (up to a week in the dugong) ^14^. These modifications, coupled with unique symbiotic microbes ^15^, presumably maximize yield from nutrient-poor seagrass.

We observed gene losses consistent with a diet comprising few animal products (**Tables S9** and **S10**). The lipid transporter *ABCG8*, a gene essential for transport of dietary cholesterol, is inactivated in the dugong and West Indian manatee. A recent study comparing 60 mammals, including non-herbivorous afrotherians, reported *ABCG8* loss in the West Indian manatee and elephants ^16^. We also confirm a loss of chitinase 5 (*CHIA5*) ^17^, a gene essential for the digestion of insect and crustacean exoskeletons, in the West Indian manatee and dugong. *ANPEP* (aminopeptidase N) is under positive selection in sirenians (**Figure 2** and **Table S7**). It encodes an enzyme that finalizes the digestion of peptides after their hydrolysis in the intestines, brain, and other cells and tissues. *ANPEP* was previously reported to be under positive selection in sirenians (West Indian manatee), cetaceans (bottlenose dolphin, *Tursiops truncatus*, and killer whale, *Orcinus orca*), and pinnipeds (walrus, *Odobenus rosmarus*) and hypothesized to play a role in reducing oxidative stress in marine mammals by adjusting cysteine and glycine availability in the glutathione metabolism pathway ^18^. This gene is also under positive selection in the ancestral branch of the dietary-diverse bat family Phyllostomidae ^19^ and has lower enzyme activity in herbivorous bird ^20^ and rodent ^21^ species. We speculate that these observations hint at modifications of this digestive enzyme in marine mammals consistent with their dietary substrates, in addition to a role in oxidative stress resistance.

Nearshore marine plants and aquatic plants from rivers and swamps are a rich source of iodine, a nutrient required to synthesize thyroid hormones (**Figure 3a**) essential for systemic energy metabolism, thermoregulation, and the integrity of many tissues ^22–24^. Both inadequate and excess iodine uptake can result in thyroid dysfunction. An elevated iodine intake is tolerated in humans, but long-term chronic intake can have adverse effects (e.g., a life-threatening thyroid storm from extreme levels of thyroid hormones) ^22,23^. Excess iodine-intake also increases the production of harmful oxygen species (ROS) by target tissues, including the thyroid gland ^25,26^. A study of captive West Indian manatees found that switching their diet from terrestrial plants to seagrass over 19 days increased blood thyroid hormone levels ^27^ – as expected given the difference in dietary iodine content. Wild manatees showed higher levels of thyroid hormones than any captive diet group ^27^, which we hypothesize resulted from a lifetime on a high-iodine seagrass diet and associated energy metabolism balance. Despite their high-iodine diet, the blood thyroid levels of wild West Indian manatees are unremarkable compared to the tropical, carnivorous Amazon River dolphin (*Inia geoffrensis*) ^27,28^. The above observations support the idea that genomic changes accompanied sirenian evolution from a terrestrial to an iodine-rich aquatic plant diet. In agreement, we observed that nearly all proteins in the thyroid hormone pathway harbor sirenian-specific amino acid substitutions (**Figure 3b** and **Table S11**). These include *TSHR* (thyroid-stimulating hormone receptor), *DUXO2* (dual oxidase 2), *DUOXA2* (dual oxidase maturation factor 2), *TPO* (thyroid peroxidase), *SLC5A5* (solute carrier family 5 member 5), *KCNQ1* (potassium voltage-gated channel subfamily Q member 1), and *KCNE2* (potassium voltage-gated channel subfamily E regulatory subunit 2). We also identified positive selection of *ATP1B4* (ATPase Na^+^/K^+^ transporting family member β4), *DIO1* (iodothyronine deiodinase 1), and *DUOX2A*, and rapid evolution of *KCNE2* and the thyroid-hormone binding albumin (*ALB*).

**Figure 3.**
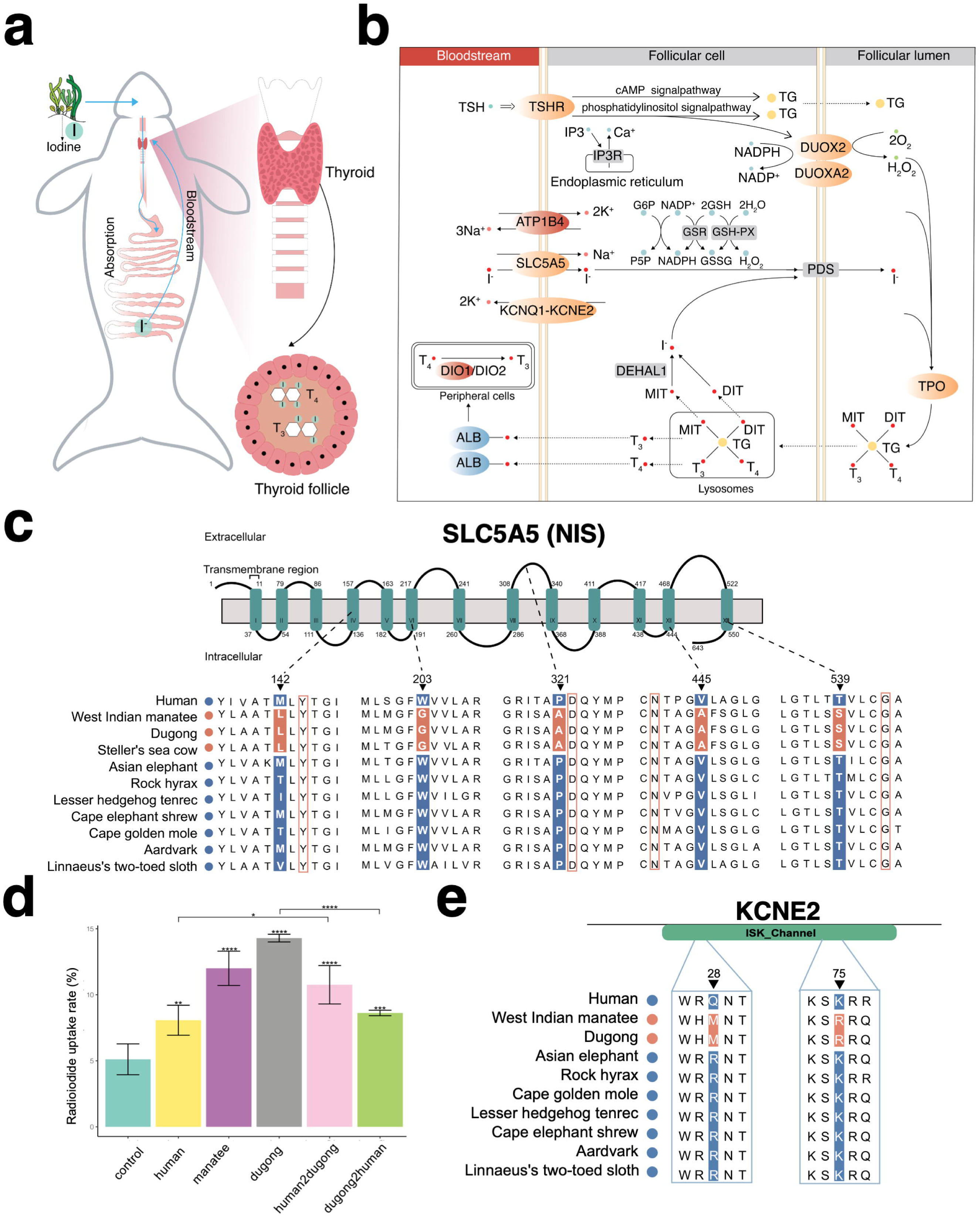
Sirenian-specific changes in iodine uptake and organification genes. **a,** Schematic diagram showing iodine uptake by a dugong, from seagrass ingestion to fermentation in the hindgut and reduction of iodine to iodide (I^-^), to iodide transport in the bloodstream to the thyroid. The thyroid is an endocrine gland consisting of follicles, structural units of epithelial cells enclosing a lumen (the colloid) harboring thyroglobulin (TG). TG is a precursor for thyroid hormones (indicated by T_3_ and T_4_) biogenesis and is also an iodide storage and carrier protein. **b,** Iodide is transported from the bloodstream to thyroid follicular cells by NIS (encoded by *SLC5A5*) interacting with KCNQ1 and KCNE2. Genes with sirenian-specific amino acid substitutions in orange and genes rapidly evolving and positively selected in sirenians in blue and red, respectively. **c,** Alignment showing an amino acid substitution in transmembrane domains of sirenian SLC5A5 (NIS). Sirenian-specific residues shown in red shading. Red boxes indicate residues shown to be important for NIS function^34–37^. **d,** Radioiodide uptake rates of human and sirenian SLC5A5 (NIS) constructs in HEK293T cells. The *y*-axis shows the percentage uptake of Na^125^I absorbed. The cDNA constructs human2dugong and dugong2human code for NIS proteins where the five sirenian-specific amino acids shown in (c) are replaced by their corresponding counterparts. Control is empty *pcDNA3.1* vector. Data are derived from three independent experiments and presented as mean ± S.D. \*\*\*\**P* < 0.0001, \*\*\**P* < 0.001, \*\**P* < 0.01, \**P* < 0.05, ANOVA with Tukey’s test. **e,** Alignment showing amino acid substitution in the ISK_Channel domain of sirenian KCNE2 (also known as LQT6). Sirenian-specific residues shown in red shading.

The transmembrane protein SLC5A5 (NIS) is the only known iodide transport protein ^29,30^. After reduction of iodine to iodide (I^-^), iodide is transported from the bloodstream into thyroid follicular cells by NIS acting in concert with the potassium transporters KCNQ1 and KCNE2 ^31,32^ (**Figure 3b**), two proteins which also have multiple sirenian-specific amino acid substitutions. Mutations in human NIS result in congenital I^-^ deficiency disorders (IDDs) ^29,33^. We identified five sirenian-specific NIS mutations. Four are in transmembrane domains (TMDs), and one is in an extracellular loop (**Figure 3c**). While none of the sirenian-specific residues have been associated with IDD to date, they are close to residues conserved in mammals shown by site-directed mutagenesis to be important for NIS function. Sirenian Leu-142 flanks Tyr-144 of TMD 4 ^34^; sirenian Ala-445 of TMD 12 flanks Asn-441, an extracellular region residue thought to mediate NIS structure via α-helix N-capping of TMD 12 ^35^; and sirenian Ala-321 is next to Asp-322 in the extracellular loop of TMD 8 and 9 ^36^. Sirenian Ser-539 flanks Gly-543 in TMD 13, a residue required for NIS cell surface targeting ^37^. We carried out I^−^ uptake assays of human and sirenian NIS in HEK293T cells, revealing higher uptake of I^-^ by dugong and West Indian manatee NIS (**Figure 3d**). Reciprocal site-directed mutagenesis of dugong and human NIS at the five residues further strengthens the evidence for more efficient iodide transport by the sirenian protein (**Figure 3d**). Our data indicate that evolutionary changes in thyroid hormone signaling, including the master regulator of iodine intake and thyroid hormone availability, played a crucial role in the secondary aquatic adaptation of sirenians.

#### The cardiovascular system

Sirenians forage in relatively shallow waters and perform short (typically a few minutes) but frequent dives ^5^. Although, perhaps not as striking as that of cetaceans, the sirenian cardiovascular system can also sustain conditions that would prove dangerous or deadly to their terrestrial relatives. For example, the Amazonian manatee does not dramatically reduce its heart rate during foraging but shows a relatively moderate 40-50 beats per minute both during dives and at the sea surface ^38^.

We identified rapid evolution of *SERPINE2* (also known as PN-1 or protease nexin-1), a gene with an emerging role in the cardiovascular system, from cardiac fibrosis to protection against arteriosclerosis and aneurysms ^39^ (**Figure 2**). *SCN5A* (cardiac Nav1.5 channel α subunit) and *KCNE2* (potassium voltage-gated channel subfamily E regulatory subunit 2) associated with heart rhythm disturbances are rapidly evolving in sirenians. Human mutations in *KCNE2* (also known as LQT6) result in long QT syndrome (LQTS) ^40^. The QT interval is an electrocardiographic measurement that measures the time taken by the cardiac ventricles to depolarize and repolarize at every heartbeat, and its prolongation or shortening is associated with morbidity and mortality ^40^. Two sirenian-specific amino acid changes in *KCNE2* were observed (**Figure 3e** and **Table S11**). The West Indian manatee has a prolonged QT interval compared to Asian and African elephants (*Elephas maximus* and *Loxodonta* spp., respectively) ^41,42^. However, like cetaceans and pinnipeds, the sirenian cardiovascular system has many unique anatomical features compared to their closest terrestrial relatives and a direct comparison remains challenging ^41,42^. Thus, although sirenian-specific *KCNE2* amino acid substitutions may be damaging in a human or elephant, such effects on the cardiovascular system may be less severe (or even adaptive) in a species with a cardiac anatomy modified for aquatic life. Instead, we propose that the unique amino acid substitution of sirenian KCNE2 is primarily associated with their unique diet by serving as a critical constituent of iodide transport (see section above).

#### The integumentary system

The skin of mammals consists of the epidermis (outer layer), the dermis (middle layer), and the hypodermis (deep layer) tissue. The sirenian epidermis, the outermost skin layer, is restructured compared to its terrestrial sister taxa. While skin appendages (hair follicles, sebaceous glands, and sweat glands) are associated with the epidermis, they project deep into the dermal layer and are absent (sweat glands) or sparsely distributed (e.g., vibrissae, specialized tactile hairs scattered over the body) in sirenians ^43^. Consistent with previous reports ^43–46^, we observed a thin epidermis and a thick dermis in the dugong (**Figure 4a**) and the West Indian manatee (**Figure S4**). We identified and validated using dugong epidermis RNA-seq reads the loss of multiple skin-associated genes (**Figure 4b** and **Table S9**). Many of these genes are also lost in cetaceans (**Supplementary Note 3**). In addition, our screen revealed sirenian loss of late cornified envelope (LCE) gene family proteins expressed by the top layer of the epidermis (**Figure 4c**). LCE gene numbers are also reduced in the afrotherian aardvark (*Orycteropus afer*; common ancestor ∼80 Mya), suggesting a role in the evolution of its sparsely haired skin ^47^.

**Figure 4.**
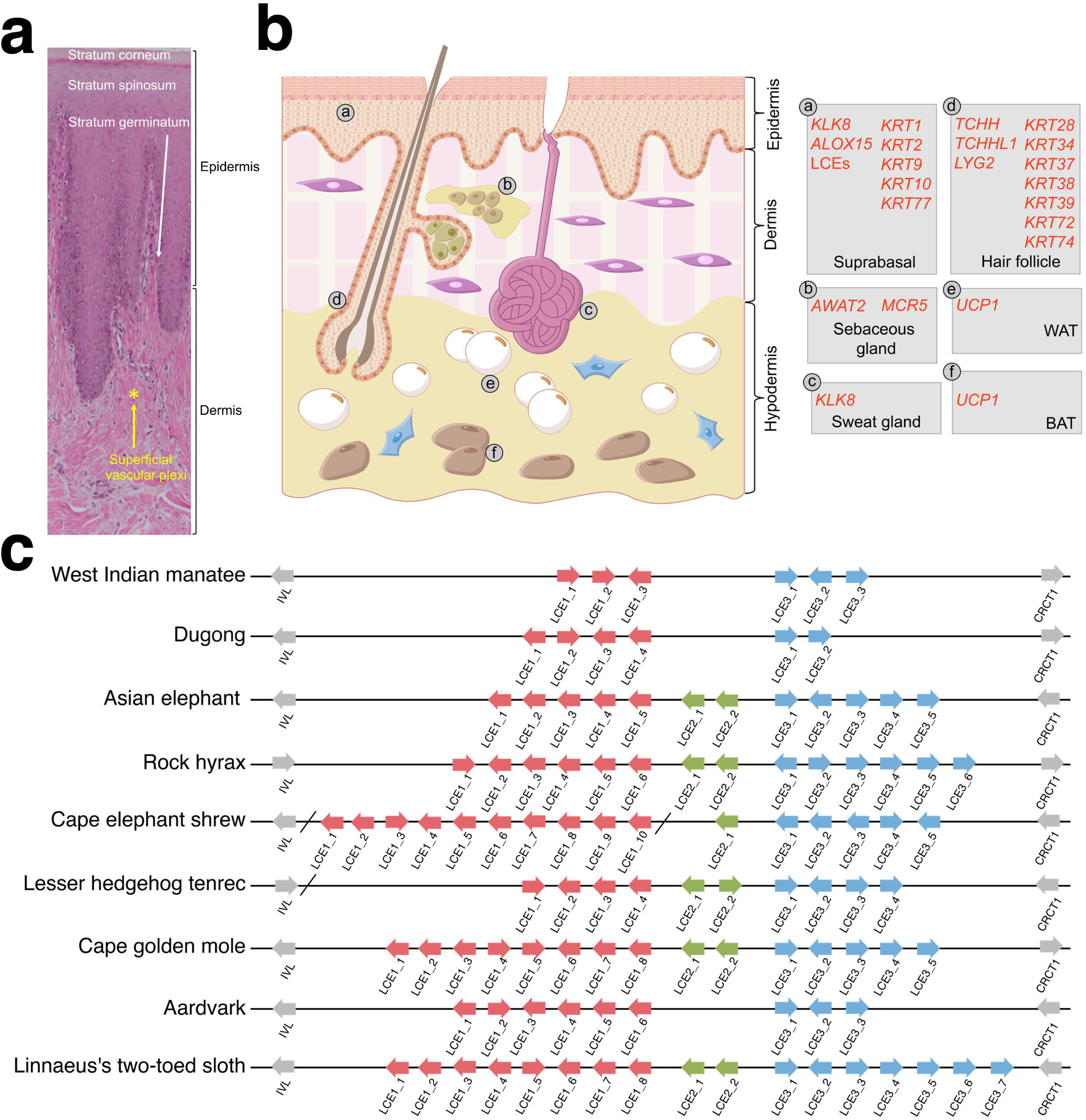
Anatomy and gene changes of the sirenian integumentary system. **a,** Histology of dorsal dugong skin **b,** Schematic drawing of human skin and associated genes with changes in (as a model of mammals). Left, overview of major skin anatomical structures: epidermis, dermis, hypodermis, and ectodermal appendages (hair follicles, sebaceous glands, and sweat glands). Right, genes and their site of expression. Inactivated genes are in red (see *main text*). WAT denotes white adipocyte; BAT, brown adipocyte. **c,** Schematic representation of late cornified envelope (LCE) gene clusters in afrotherian genomes. Arrows indicate genes and their direction of transcription; different clusters are indicated by colored boxes. The slashes on each line indicate the ends of scaffolds in the assembled genomes.

#### Daily activity patterns

Most terrestrial mammals rely on circadian rhythmicity, a molecular clock mediated by the sun’s daily cycle, to mediate activity patterns ^48,49^. In contrast, many marine mammals inhabiting coastal reef habitats and shallow waters rely heavily on the lunar (moon) cycle and its effect on water depth, food availability, and temperature ^48^. The pineal gland is absent or non-functional in sirenians, cetaceans, and some terrestrial mammals. These species do not encode genes associated with the synthesis and reception of melatonin, a hormone essential for light stimulation of the circadian clock ^50–54^. Sirenians do not show a ‘classical’ diel (24-hour) activity pattern, have unihemispheric slow wave sleep (i.e., one brain half is awake), and appear to respond behaviorally to tidal currents (tides may restrict foraging) and seasonal changes in water temperature ^5,55,56^.

We identified sirenian-specific amino acid substitutions in most core circadian clock genes (**Figure 5a**, **Figure 3** and **Table S11**), including numerous substitutions in all Period circadian regulator genes (five in *PER1*, 20 in *PER2*, and 13 in *PER3*). PER2 is expressed by circadian pacemaker cells of the hypothalamic suprachiasmatic nucleus (SCN) and binds to cryptochromes (CRY) to regulate light-associated circadian rhythmicity and timing (including sleep patterns) ^57^. A coimmunoprecipitation assay showed that dugong PER2 bound CRY1 better than wildtype human PER2 or human PER2 with a sirenian substitution (C1220P) in the CRY-binding domain (**Figure 5b,c**). The latter result suggests that at least one of the 19 other sirenian-specific PER2 residues is required for the enhanced CRY-binding. Our analysis also revealed loss of *KCNK18* (also known as TRESK, TWIK-related spinal cord K^+^ channel) (**Figure 5d** and **Tables S9** and **S10**), a circadian clock-regulated ion channel. *KCNK18* is highly expressed in the SCN, and *Kcnk18* ^−/−^ mice cannot use light to differentiate between day and night ^58^. Taken together, the lack of a pineal gland and a genetic background where many circadian genes have unique amino acid changes or are lost strengthens the idea that the sirenian circadian clock has been recalibrated.

**Figure 5.**
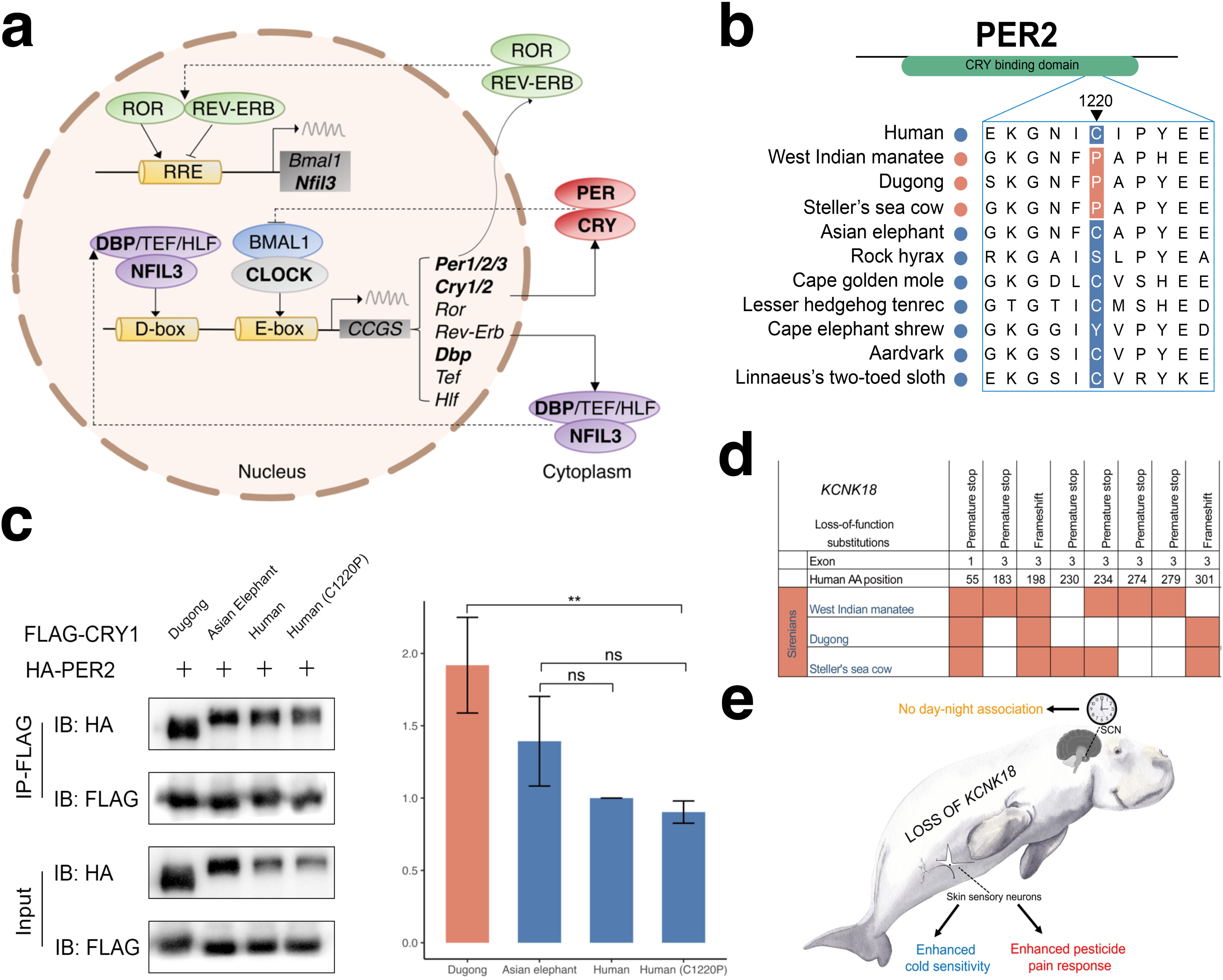
Multiple gene changes may underly the circadian activity of sirenians. **a,** Diagram of core circadian clock network. Genes with evolutionary changes in sirenians are bolded. D-box and E-box denote promoter elements found upstream of clock-controlled genes regulated by the transcription factors NFIL3 and CLOCK. **b,** Alignment showing an amino acid substitution in the CRY binding domain of sirenian PER2. A sirenian-specific proline at residue 1220 is shown in red shading. **c,** Left, A coimmunoprecipitation assay reveals the stronger binding ability of dugong PER2 with CRY1, as compared with human. Human C1220P denotes human PER2 mutated to match the dugong residue 1220. Right, quantification of densitometry analysis of blot using ImageJ. Shown as arbitrary density units (AU). \*\**P* < 0.01, \**P* < 0.05, ANOVA with Tukey’s test. **d,** Overview of inactivating substitutions in sirenian *KCNK18*. Colored cells indicate substitutions. **e,** Illustration of anatomical sites where *KCNK18* plays important roles and potential functional outcomes of its loss in sirenians.

#### Gene loss and maladaptation in a human-altered environment

A gene can become lost (inactivated) by chance or because it no longer serves a relevant function. Such losses can be functionally compensated by other genes ^59^ or may be adaptive ^60^. For example, the loss of dozens of genes re-organized the skin of cetaceans ^61^ and sirenians (see above) over millions of years, allowing semi-aquatic ancestors to become fully aquatic.

Climate change and human activities can influence marine life at all levels. Threats to sirenians, particularly through loss of aquatic plant habitats, include pollution of waterways, fishing operations, and coastal dredging and reclamation, all of which may be exacerbated by changing climate patterns ^62–64^. While the herbivorous sirenians are at a much lower position in the food web and show relatively low levels of persistent organic pollutants compared to cetaceans at similar geographic locations ^65^, they are likely also sensitive to certain manmade chemicals. It follows that such conditions can far outpace natural selection and that historical gene loss that may have been adaptive or tolerated in an ancestral environment can become maladaptive in a modern environment, especially in species with a large generation time ^66^. Sirenians are long-lived (∼70 years), with a generation time of ∼20 years for the West Indian manatee and ∼27 years for the dugong ^5,67^. We identified loss of three genes that may be disadvantageous today: *PON1*, *KCNK18*, and *CES3*.

Loss of *PON1* (paraoxonase 1) in most aquatic mammals – sirenians (including the extinct Steller’s sea cow *Hydrodamalis gigas*), cetaceans, pinnipeds, beavers, and the sea otter *Enhydra lutris* – is thought to reduce oxidative stress and associated inflammation induced by diving ^68,69^. *PON1* was also identified in our pseudogene analysis, confirming a single shared premature stop codon event in the dugong and West Indian manatee as well as several independent inactivation mutations (**Tables S9** and **S10**, and **Figure S5**). The presumably adaptive loss of *PON1* likely became maladaptive in aquatic mammals after the industrialization of agriculture as the enzyme protects against organophosphate pesticides (the ozone forms of chlorpyrifos and diazinon) ^68^.

*KCNK18* loss was probably not damaging (see above) and may similarly expose sirenians to natural and anthropogenic threats. Shared and unique *KCNK18* inactivating mutations were observed in the dugong, Steller’s sea cow, and West Indian manatee (**Tables S9** and **S10**, and **Figure 5d**). Besides the hypothalamic SCN, skin sensory neurons express the gene, and *Kcnk18* knockout mice show elevated pain and avoidance behaviors when exposed to pyrethroid insecticides ^70^ or temperatures below 20 °C ^71^. Given its geographic range (**Figure 1c**), the Florida manatee (i.e., a subspecies of the West Indian manatee) manifests cold stress syndrome (CSS), a condition resulting from prolonged exposure to water temperatures below 20 °C, and characterized by multiple physiological changes and comorbidities of unknown genetic cause ^5,72,73^. We speculate that loss of *KCNK18* decreases manatee temperature stress tolerance (**Figure 5e**) and that CSS is similar to semelparity in marsupials ^74,75^ in that a progressive and systemic deterioration of body condition and physiological function is mediated by an endocrine factor, perhaps from elevated levels of the stress hormone cortisol. Extant sirenians have relatively thin blubber compared to cetaceans and have lost *UCP1* (**Supplementary Note 3**), which could render them further susceptible to cold temperatures. Dugongs should arguably also be inherently sensitive to cold temperatures but have a more insulating integument, higher metabolic rate, and live in warmer waters throughout the year than manatees ^45,76^. The extinct Steller’s sea cow related to the dugong further evolved a huge body size and thicker blubber to survive in the cold, sub-Arctic environment ^77^.

We identified loss of carboxylesterase 3 (*CES3*; also known as *ES31*) in sirenians (**Table S9** and **S10**), cetaceans (loss of first nine coding exons), and phocids (10-bp deletion in Phocidae, the largest pinniped family ^78^) (**Figure S6**). Loss of the *CES3 –* by the West Indian manatee and killer whale ^51^, and by four cetaceans and two hippos ^79^ – has previously been reported but not discussed. A premature stop codon is shared by all sirenians, while the dugong and Steller’s sea cow share an additional stop codon (**Figure S6**). *CES3* and its homolog *CES1* have been associated with macrophage cholesterol ester metabolism and are both also expressed by the liver and intestines, where they break down ester bonds of endogenous compounds and xenobiotics ^80–82^. While *CES3* expression is much lower than *CES1* and its enzyme has several magnitudes lower catalytic efficiency than CES1 for many compounds (and, thus, *CES3* loss in marine mammals is likely compensated), carboxylesterase 3 may show exclusive specificity against manufactured compounds such as pesticides.

### Diversity and demography of vulnerable and recently extinct dugongs

Despite their numbers, the ∼165,000 dugongs in Australian waters are listed as *Vulnerable* (at high risk of extinction in the medium-term future) by the International Union for Conservation of Nature (IUCN) ^83,84^. The number of dugongs in other locations is orders of magnitude lower. The dugong recently became functionally extinct in Chinese ^85,86^ and Japanese waters ^87^ and is at risk elsewhere in Asia, Oceania, and eastern Africa ^84,88^.

We considered the genomic diversity and demography of dugongs from ten locations (**Figure 6a**). Short-read whole-genome resequencing data were generated from seven locations (99 individuals) spanning 2,000 km of the Australian east coast (from Torres Strait to Moreton Bay, Queensland) (**Table S12**). Resequencing data (one individual carcass each) was also obtained from two other Australian locations, Coogee Beach (New South Wales) ^77^ and Exmouth Gulf (Western Australia), and waters off Okinawa (Japan). Principal component analysis (PCA) indicated that the individuals from Exmouth Gulf on the Australian west coast and Okinawa are genetically distinct to dugongs from the Australian east coast (Queensland and Coogee Beach) (**Figure 6b**). The Coogee Beach individual clustered with Moreton Bay and Hervey Bay individuals. Because this dugong stranded ∼750 km from the accepted eastern Australian range during the summer (November), we speculate it represents one of the few instances ^89^ of seasonal long-distance ranging of a single individual from a population in close geographic proximity to Moreton Bay.

**Figure 6.**
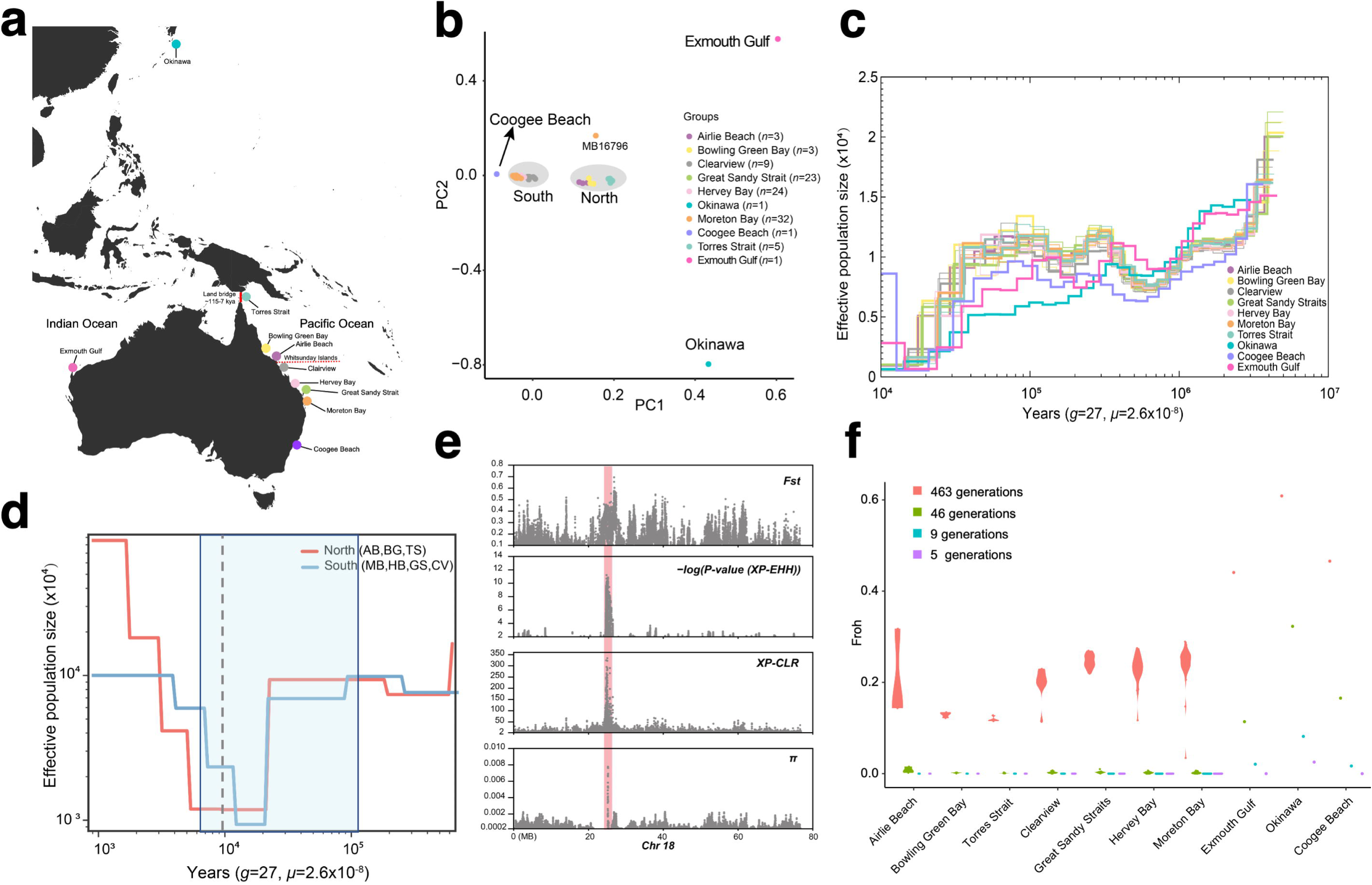
Demographic history of dugongs from 11 locations. **a,** Map indicating approximate sites of origin of resequenced dugongs. The red continuous line indicates a historical Torres Strait land border (115–7 kya); the dotted red line, a potential ecological barrier at or near the Whitsunday Islands on the Great Barrier Reef. Geographical locations are approximate. **b,** Principal component (PC) analysis of the dugong individuals. The gray highlighting shows clustering consistent with geographic locations south and north of the Whitsunday Islands also reported by ^95^. **c,** Demographic history of dugong individuals inferred using PSMC. Okinawa, Exmouth Gulf, and Coogee Beach are single individuals, while three individuals from the remaining Queensland locations are plotted. **d,** Effective population size of populations inferred using SMC++. The most recent Torres Strait land border period is shaded in blue; the grey continuous line indicates the estimated emergence time of a potential ecological barrier at or near the Whitsunday Islands. **e,** A ∼2 Mb selective sweep region on chromosome 18 identified by comparing population groups north and south of the Whitsunday Islands. Population differentiation (*Fst*), normalized XP-EHH, XP-CLR, and relative nucleotide diversity (π) are plotted. Note the positive XP-EHH value, indicating selection acting on the northern dugong population. **f,** Inbreeding in dugong individuals up to 463 generations ago (12,501 years, ROHs>100 kb); 46 generations ago (1,242 years; ROHs>1 Mb); nine generations ago (243 years ago; ROHs>5 Mb); and five generations ago (135 years; ROHs >10 Mb). The *y*-axis shows F_ROH_, the sum length of all ROHs of a particular length cutoff divided by the autosomal genome length; the x-axis indicates various dugong populations.

Pairwise sequential Markovian coalescent (PSMC) ^90^ analysis of dugong autosomes was used to track changes in *N_e_* during the Pleistocene (about 2.6 Mya to 20 kya) (**Figure 6c** and **Figure S7**). All dugongs showed a *N_e_* decline in the mid-Pleistocene until ∼400-500 kya also observed in a Steller’s sea cow population off the Arctic Commander Islands ^77,91^. Individuals from the seven eastern Queensland locations had near-identical demographic histories. The Coogee Beach individual’s PSMC curve mirrored the Queensland individuals, agreeing with the above PCA. The Exmouth Gulf individual from the Indian Ocean showed a distinct *N_e_*to the other Australian populations from the Pacific Ocean range but, nevertheless, a curve that likely reflects the similar environmental conditions across the Australian continent (**Figure 6c**). In contrast, the *N_e_*of the individual from Japanese waters declined continuously over the last 400,000 years.

Indian Ocean and Pacific Ocean dugong populations off Australia became separated by a land bridge across the Torres Strait (**Figure 6a**) from ∼115 kya until ∼7 kya when sea levels rose to their present levels ^5,92^ and allowed resumed panmixia with populations from the Indian Ocean and further southward migration. Genetic evidence of this event on dugong demography has been reported by several studies ^67,93,94^. Further interrogation of the 99-individual Queensland dataset, revealed pronounced genetic structure into a northern and a southern group since ∼10.2 kya, consistent with a recently reported potential ecological barrier at or near the Whitsunday Islands on the Great Barrier Reef ^95^ (**Figure 6a**, **Supplementary Note 4** and **Figure S8**). We next assessed proxies of genomic diversity, relative nucleotide diversity (π) (**Figure S8e**) and genome-wide heterozygosity (**Figure S8f**). The northern group showed higher diversity, suggesting that individuals from these locations are more ‘genetically healthy’, consistent with effective population size differences in the recent past reconstructed using SMC++ (**Figure 6d**). Four complementary methods (XP-EHH ^96^, XP-CLR ^97^, π, and *F_st_*) were used to detect regions with putative selective sweeps between the northern and southern groups. We identified a ∼2 megabase region (24.48-26.50 Mb) on chromosome 18 under positive selection in the northern dugong group (**Figure 6e**). This region contains six annotated protein-coding genes (**Table S13**). They comprise three immunoglobulin genes, the nuclear pore transporter *NUP42* (also known as *NUPL2*), ClpXP protease (*CLPX*), and an apparent *GLPX* paralog (‘*GL*PX-like’). NUP42 is expressed by stress granules, membraneless cell compartments formed in response to various environmental stressors ^98^, in organisms ranging from mammals ^99^ to yeast ^100^. The mitochondrial protein CLPX is required to synthesize oxygen-binding hemoglobin and buffers oxidative stress ^101,102^. While it is difficult to conclude the driving force behind the selective sweeps of *NUP42*, *CLPX*, and *CLPX-like* loci, sirenians are vulnerable to climate change and regional environmental conditions that dramatically alter their nearshore habitats (particularly seagrass growth) ^63^. Northern Queensland has seen distinct climate change events after the formation of the ostensible ecological barrier at the Whitsunday Islands ∼10.2 kya. The sea level on the north-east Queensland coast has continuously decreased since about 6,000 years ago ^103,104^. In addition, organisms that inhabit nearshore habitats must also acclimate to and survive oxidative stress stemming from changes in salinity by high seasonal summer rainfall and coastal run-offs ^105^. Northern Queensland has such environmental conditions.

Extinction events can often be predicted from the genetic history of a species, with loss of genomic diversity reflecting a dwindling population. The 99 dugongs from the Queensland coast had a mean genome-wide heterozygosity of 9.83 **×** 10^-4^ (standard deviation (SD) = 6.75 **×** 10^-5^ heterozygotes per site) (**Figure S9a**). The heterozygosity of the Exmouth Gulf (7.73 **×** 10^-4^) individual was slightly lower, and markedly lower in the individual from the now functionally extinct Okinawa population (5.65 **×** 10^-4^). The heterozygosity values of the Australian dugongs were similar to a West Indian manatee from Florida (1.08 **×** 10^-3^) ^18^ and the critically endangered Yangtze finless porpoise *Neophocaena asiaeorientalis* (∼1 **×** 10^-3^) ^106,107^ but an order of magnitude higher than the extinct baiji *Lipotes vexillifer* (1.23 **×** 10^-4^) ^108^ and much higher than the critically endangered vaquita *Phocoena sinu* (9.04 **×** 10^-5^) ^109^.

The length and number of runs of homozygosity (ROHs) in a genome reflect the level of inbreeding and can provide conservation management guidance ^110,111^. Longer ROHs suggest recent inbreeding. The total length and number of ROHs were generally small and similar across the 99 dugongs sampled on the Queensland coast (**Supplementary Note 4**), with no evidence of inbreeding in recent times (since 1,242 years ago) (**Figure 6f** and **Table S14**). The individual from Exmouth Gulf (F_ROH>1Mbp_ = 0.021, nine ROHs spanning 60.1 Mb) showed evidence of inbreeding as recent as nine generations ago (243 years), consistent with a small or isolated population at this time. An illustration of how environmental events may affect population size in this locality, is tropical cyclone Vance (1999) which appears to have resulted in large-scale emigration from Exmouth Gulf, reducing the dugong population from 1,000 to less than 200 individuals in five years ^112^. It follows that other migrations on a population scale in the last two centuries may have improved the genetic diversity of the Exmouth Gulf ancestors. The historical demography (PSMC) of the Okinawa individual aligned with its ROH estimates. One-third of the genome is in ROH segments above one megabase (389 ROHs spanning 934.4 Mb), and there is evidence of inbreeding as recently as 135 years ago (F_ROH>10Mbp_ = 0.082, four ROHs spanning 73.6 Mb). This pattern is consistent with an ancient population bottleneck and subsequent extensive inbreeding since ∼135 years ago, mirroring a *Critically Endangered* Sumatran rhinoceros (*Dicerorhinus sumatrensis*) population on the Malay Peninsula ^113^. Dugongs in Japanese waters were hunted for centuries, and the Okinawan dugong showed a further dramatic reduction in population size from the late 1900s until its likely functional extinction in 2019 ^87^.

We conclude that while dugongs on the Australian east and west coasts have a genomic diversity comparable to other marine mammals of conservation concern, most populations (as indicated by our resequencing of 99 individuals from the Queensland coast) are in numbers that likely limit inbreeding. Viewed through a genetic lens, these populations can be considered relatively robust and healthy. In contrast, the dugong from the functionally extinct Okinawan population showed a continuous decline in effective population size, lower genomic diversity, and an ROH pattern consistent with extensive inbreeding for millennia and, likely, climatic ages.

## Discussion

By comparing afrotherian genomes and validating against 120 mammal species, we identified unique amino acid substitutions in protein-coding genes that may underlie sirenian-specific features such as tolerance to a high-iodine aquatic plant diet and an activity pattern heavily reliant on lunar tidal currents and water temperature fluctuations. Because the ancestors of extant manatees and the dugong diverged at crown Sirenia ∼30 Mya ^2^, these gene changes were likely critical for the transition to an aquatic habitat. Many of the same genes of the integumentary system, particularly expressed in the outer layers of the skin ^51,79,114–119^, were lost by both sirenians and cetaceans. This further supports the idea that convergent gene loss occurs in species with similar ecological pressures ^120^ (here, the transition to a fully aquatic lifestyle by distantly related species over ∼50-60 My of evolution). We also observed gene losses that may be maladaptive in a modern environment, including the previously reported paraoxonase 1 gene (*PON1*) ^68^. Sirenian *KCNK18* loss is possibly related to their shift in activity patterns and may render sirenians susceptible to impacts on water temperatures by climate change and human activities. Its role in cold stress syndrome (CSS) should be investigated. Our population genomics analysis offers insights into dugong diversity and demography. We confirm ^95^ and (for the first time) date a north-south genetic break on the Australian east coast and reveal a ∼2 Mb genetic sweep region that may be associated with differences in salinity or other environmental factors between the north and south coast. The findings from a functionally extinct population (Okinawa, Japan) highlights an urgency to obtain whole-genome resequencing data from dugong populations worldwide to support conservation efforts. Recent human activities in the last 100 years, including overfishing and coastal development that reduced seagrass abundance, likely fast-tracked extinction of dugong populations in Chinese and Japanese waters ^64,85–87^, and is a considerable risk factor elsewhere. Our study reveals new insights into sirenian biology and the transition of terrestrial mammals to an aquatic lifestyle and provides a basis for future genomics explorations.

## Online Methods

### Sample collection and research ethics

Fetal liver tissue (sample D201106) for reference genome sequencing was obtained from a recently deceased near-term female dugong fetus recovered from a cow that was hunted illegally in the Burrum Heads region of Hervey Bay Queensland in November 2020. This fetal liver sample and a skin sample from a dugong (D110419) sampled after an indigenous subsistence hunt in Torres Strait in 2011 were used for RNA-sequencing.

Skin samples for whole-genome resequencing were collected from 99 dugongs from seven geographic locations on the east coast of Australia: Airlie Beach (AB, *n*=3), Bowling Green Bay (BG, *n*=3), Clairview (CV, *n*=9), Great Sandy Strait (GS, *n*=23), Hervey Bay (HB, *n*=24), Moreton Bay (MB, *n*=32), and Torres Strait (D, *n*=5; prefix ‘TS’ elsewhere in the manuscript) (**Table S12** and **Figure 6a**). Briefly, skin was collected from the dorsum of wild free-swimming dugongs at each location, except Torres Strait, using a handheld scraper device ^121^. In the Torres Strait, skin was excised from fresh dugong carcasses post-hunt by local Traditional Owners at Mabiuag Island. Skin samples were stored in salt-saturated 20% DSMO and stored frozen at -20 °C until further genetic processing for resequencing (see ^95^). Skin samples for histology were sub-sampled from the Torres Strait specimens and stored in 10% neutral buffered formalin until sectioned and stained.

The dugong skin samples were collected under the following permits issued to J.M.L.: the University of Queensland Animal Ethics Permits SBS/360/14, SBS/181/18, Scientific Purposes Permits WISP07255110 and WISP14654414, Moreton Bay Marine Parks Permit #QS2000 to #QS2010CV L228, Great Sandy Marine Parks Permit QS2010-GS043, and Great Barrier Reef Marine Park Permits #G07=23274:1 and G14/36987.1.

Sampling and species distribution maps were generated using the R package ‘OpenStreetMap’ ^122^

### Genome sequencing

High-molecular weight DNA was extracted from a fetal liver sample (D201106). DNA quantity, purity and integrity were assessed by Qubit fluorometry (Invitrogen), Nanodrop spectrophotometry (Thermo Fisher Scientific), and pulse-field gel electrophoresis. Single tube long fragment read (stLFR) libraries ^123^ were sequenced on a MGISEQ-2000 sequencer. A total of ∼358Gb (∼100×) stLFR clean reads were generated after removing low-quality reads, PCR duplicates, and adaptors using SOAPnuke (v1.5) ^124^. Hi-C libraries (Lieberman-Aiden et al., 2009) were prepared from the same fetal liver sample. Hi-C data (200 Gb 150 bp paired-end reads) were generated on the BGISEQ-500 platform.

### RNA sequencing

RNA from fetal liver (sample D201106) and skin (sample D110419) were sequenced on the BGISEQ-500 platform to generate 86.7 and 96.3 Gb of raw RNA-seq data, respectively.

### Genome assembly

We carried out *k*-mer analysis with ∼50× clean reads to estimate the genome size (*G*) of dugong using the formula *G* = num of *k*-mer/peak-depth.

The stLFRdenovo pipeline (https://github.com/BGI-biotools/stLFRdenovo), which is based on Supernova v2.11 ^125^ and customized for stLFR data, was used to generate a *de novo* genome assembly. GapCloser v1.12 ^126^ and clean stLFR reads (with the barcode removed with https://github.com/BGI-Qingdao/stLFR_barcode_split) were used to fill gaps. Redundans v0.14a ^127^ was used to remove heterozygous contigs. Clean paired-end Hi-C reads validated by HiC-Pro v3.2.0_devel ^128^ were used to construct chromosome clusters with the 3D de novo assembly (3D DNA) pipeline v170123 ^129^. The assembly was further improved by interactive correction using Juicebox (v1.11.08) ^130^.

Assembly quality was assessed using BUSCO (Benchmarking Universal Single-Copy Orthologs) BUSCO v5.4.3 ^8^, employing the gene predictor AUGUSTUS (v3.2.1) ^131^ and a 9,226-gene BUSCO mammalian lineage data set (mammalia_odb10).

### Genome annotation

The sex of the sequenced individual (i.e., to determine if we could assemble the Y chromosome) was determined by visual inspection of the specimen prior to dissection, as well as BLAST interrogation of African elephant ^132^ and published dugong ^133^ sex chromosome genes against our initial stLFR assembly and by comparing the mapping rate of clean stLFR sequencing reads against chromosome X and autosomes of the chromosome-level genome assembly (see the whole-genome resequencing section below for method). All approaches suggested that the sequenced individual was female.

We identified repetitive elements by integrating homology and de novo prediction data. Protein-coding genes were annotated using homology-based prediction, de novo prediction, and RNA-seq-assisted prediction methods.

Homology-based transposable elements (TE) annotations were obtained by interrogating a genome assembly with known repeats in the Repbase database v16.02 ^134^ using RepeatMasker v4.0.5 (DNA-level) ^135^ and RepeatProteinMask (protein-level; implemented in RepeatMasker). De novo TE predictions were obtained using RepeatModeler v1.1.0.4 ^136^ and LTRharvest v1.5.8 ^137^ to generate database for a RepeatMasker run. Tandem Repeat Finder (v4.07) ^138^ was used to find tandem repeats (TRs) in the genome. A non-redundant repeat annotation set was obtained by combining the above data.

Protein-coding genes were annotated using homology-based prediction, de novo prediction, and RNA-seq-assisted [generated from fetal liver and skin] prediction methods.

For homology-based prediction, proteins sequences from five mammals [African bush elephant (*Loxodonta africana*) assembly Loxafr3.0; Cape elephant shrew (*Elephantulus edwardii*) assembly EleEdw1.0; aardvark (*Orycteropus afer*) assembly OryAfe1.0; West Indian manatee (*Trichechus manatus latirostris*) assembly TriManLat1.0, and human (*Homo sapiens*) assembly GRCh38.p12)] were downloaded from NCBI. These protein sequences were aligned to the repeat-masked genome using BLAT v0.36 ^139^. GeneWise v2.4.1 ^140^ was employed to generate gene structures based on the alignments of proteins to a genome assembly. De novo gene prediction was performed using AUGUSTUS v3.2.3 ^141^, GENSCAN v1.0 ^142^, and GlimmerHMM v3.0.1 ^143^ with a human training set. For RNA-seq-assisted gene prediction, 150 bp PE reads from fetal liver and skin, generated on an BGI-SEQ 500 instrument, were filtered using Flexbar v3.4.0 ^144,145^ with default settings (removes reads with any uncalled bases). Any residual ribosomal RNA reads [the majority ostensibly removed by poly(A) selection prior to sequencing library generation] were removed using SortMeRNA v2.1b ^146^ against the SILVA v119 ribosomal database ^147^. Transcriptome data (clean reads) were mapped to the assembled genome using HISAT2 v2.1.0 ^148^ and SAMtools v1.9 ^149^, and coding regions were predicted using TransDecoder v5.5.0 ^150,151^.

A final, non-redundant reference gene set was generated by merging the three annotated gene sets using EvidenceModeler v1.1.1 (EVM) ^152^. The gene models were translated into amino acid sequences and used in local BLASTp ^153^ searches against the public databases Kyoto Encyclopedia of Genes and Genomes (KEGG; v89.1) ^154^, NCBI non-redundant protein sequences (NR; v20170924) ^155^, Swiss-Prot (release-2018_07) ^156^, TrEMBL (Translation of EMBL [nucleotide sequences that are not in Swiss-Prot]; release-2018_07) ^157^, and InterPro (v69.0) ^158^. The gene set was also examined using BUSCO v5.4.3 and its mammalia_odb10 gene set (’transcriptome mode’).

### Phylogeny and divergence time estimation

Genome and protein sequences of eight afrotherians and Linnaeus’s two-toed sloth *Choloepus didactylus* (outgroup) (see **Table S2**) were downloaded from the NCBI or DNA Zoo databases.

We identified single-copy gene family orthologs using OrthoFinder (v2.5.4) ^159,160^. The coding sequences (CDS) for each species were aligned using PRANK v70427 ^161,162^ and filtered by Gblocks v0.91b ^163^ to identify conserved blocks (removing gaps, ambiguous sites, and excluding alignments less than 300 bp in size). Finally, 7,695 single-copy genes were concatenated into supergenes for phylogenetic analyses.

To identify conserved non-exonic elements (CNEEs) we generated whole-genome alignments (WGAs) using LASTZ v1.04.22 ^164^ with the parameters ‘H=2000 Y=3400 L=3000 K=2400’ and our dugong reference genome (Ddugon_BGI) as the reference. We then merged aligned sequences using MULTIZ (version 11.2) ^165^. To estimate the non-conserved model, we employed phyloFit (v1.4) in the PHAST package ^166^ with 4d sites in the Afrotheria alignments and the topology using the default parameters. We ran phastCons with the non-conserved model to estimate the conserved regions with the parameters ‘target-coverage = 0.3, expected-length = 45, rho = 0.31’. Exon regions were excluded from the highly conserved elements to generate 627,279 CNEEs with a total length of 103.9M longer than or equal to 50 bp.

Mitochondrial genomes and protein-coding sequences for the following species were obtained from NCBI GenBank: dugong (NC_003314.1), West Indian manatee (MN105083.1), Asian elephant (Elephas maximus; NC_005129_CDS), rock hyrax (*Procavia_capensis*; NC_004919.1), lesser hedgehog tenrec (*Echinops telfairi*; NC_002631.2), golden mole (*Chrysochloris asiatica*; NC_004920.1), aardvark (NC_002078.1), Cape elephant shrew (NC_041486.1), and Linnaeus’s two-toed sloth (*Choloepus didactylus*; NC_006924.1). We used MARS ^167^ to rotate the mitochondrial sequences to the same origin as the dugong.

Separate maximum-likelihood (ML) phylogenetic trees of eight afrotherians and Linnaeus’s two-toed sloth (outgroup) were generated with RaxML v8.2.9 ^168^ (1,000 bootstrap iterations) using coding sequences from 7,695 genes, 1,127,156 fourfold degenerate sites in the 7,695 genes, 5,508 single-copy Benchmarking Universal Single-Copy Ortholog (BUSCO) genes ^8,169^, 627,279 conserved non-exonic elements (CNEEs), mitochondrial genomes, and 13 mitochondrial coding sequences. The resulting tree with the highest GTRGAMMA likelihood score was selected as the best tree.

We also used ASTRAL-III v5.6.2 ^170^ to generate a coalescent species tree from non-overlapping 20 kb windows (to minimize linkage between subsequent windows from the WGAs generated above (see ^171^). Briefly, after excluding windows from alignments less than 2,000-bp in size and with more than 25% gaps, 152,478 windows were used to generate window-based gene trees (WGTs) using RaxML (with 1,000 bootstraps). WGTs with mean bootstrap support ≥80% were input into ASTRAL-III (with default parameters) to estimate an unrooted species tree. Apart from the mitochondrial trees (its whole genome and protein coding genes were not identical), the nuclear genome-derived ML trees showed the same topology as the ASTRAL WGT tree.

Divergence times between species was estimated using MCMCTree [a Bayesian molecular clock model implemented in PAML v4.8 ^172^] with the JC69 nucleotide substitution model, and the concatenated whole-CDS supergenes as inputs. We used 100,000 iterations after a burn-in of 10,000 iterations. MCMCTree calibration points (million years ago; Mya) were obtained from TimeTree ^173^ [Cape golden mole-lesser hedgehog tenrec (58.4-81.7 Mya), Cape golden mole-Cape elephant shrew (63.0-87.5 Mya), Cape golden mole-aardvark (54.9-89.3 Mya), Cape golden mole-West Indian manatee (76.0-80.8 Mya), Linnaeus’s two-toed sloth-West Indian manatee (84.0-97.9 Mya)] and a recent manuscript ^2^ that employed a total-evidence approach (incorporating morphological, molecular, temporal, and geographic data from living and fossil species) to estimate a divergence of Trichechidae (i.e., West Indian manatee ancestor) from Dugongidae (i.e., dugong) ancestors 31.7-36.7 Mya.

To further examine the relationship between paenungulates, we also considered retroelements. Retroelements (i.e., LINEs, SINEs, and LTRs) are considered near homoplasy-free markers given their insertion mode ^174–177^. We employed a recently developed pipeline ^176^ that requires the 2,000 bp flanking a retroelement insertion site to assign informative phylogenetic markers from pairwise whole-genome alignments (here, with Ddugong_BGI as the reference genome). The Kuritzin-Kischka-Schmitz-Churakov (KKSC) test ^178^ was used to assess presence/absence matrixes.

### Gene family analysis

Gene family expansion and contraction analysis was performed using CAFE v4.2 ^179^ with the our consensus phylogenetic tree generated (also see **Figure 1c**) as the input. Expanded and contracted gene families on each branch of the tree were detected by comparing the cluster size of each branch with the maximum-likelihood cluster size of the ancestral node leading to that branch. A smaller ancestral node indicates gene family expansion, whilst a larger ancestral node indicates family contraction. Gene families with a *P*-value < 0.01 were defined as significantly expanded or contracted in a branch of interest.

### Sirenian gene selection

Selection signatures of the 7,695 single-copy gene family orthologs in our nine-species data set were identified using their coding sequences and PAML ‘codeml’ v4.8 ^172^.

We tested for positively selected genes (PSGs) on sirenian branches by comparing branch-site models, allowing a codon site class with (d*N*/d*S*; also known as ‘omega’, ω) >1 along foreground branches, with branch-site null models. We identified sites under positive selection using Bayes’ Empirical Bayes (BEB) in PAML ^180^ and a Benjamini–Hochberg *P*-value cut-off set at 0.05.

Rapidly evolving genes [REGs, i.e., genes with an elevated d*N*/d*S*] in sirenians were identified using the PAML branch model. The two-ratio model (model=2) allows one ratio for background branches and another for foreground (sirenians) branches, while the one-ratio model (model=0) enforces the same ratio for all branches. Genes with a *P*-value (computed using the χ^2^ statistic) less than 0.05 and a higher ω value in the foreground lineage were considered REGs.

### Gene loss in the sirenian lineage

To screen for gene loss in sirenians, defined as genes harboring premature stop codons and/or frameshifts in a species, we employed a previously published approach ^181^. Briefly, the longest human protein sequence for each gene was mapped to genomes of the dugong, West Indian manatee, rock hyrax, and Asian elephant using BLAT v36 ^139^ and genBlastA v1.0.1 ^182^. Next, the mapped mapped genomic regions and 1,000 bp upstream and downstream were examined for disruptions (nonsense mutations and frameshifts) to the gene coding sequences using GeneWise v2.4.1 ^140^. We removed loci hits belonging to large gene families (including olfactory receptors, zinc finger proteins, and vomeronasal receptors) and predicted proteins or intronless cDNA/expressed sequences ^183^. False positives with disruptive mutations introduced by GeneWise, sequencing errors, or annotation errors were removed following the steps in ^181^. Candidate pseudogenes with multiple disruptions were manually inspected to remove short or low-quality alignments. We also interrogated raw sequencing reads from dugong and West Indian manatee to confirm each disruption.

Gene loss events were dated using the same procedure employed by ^184^ (based on ^183,185^). Briefly, a gene is assumed to have evolved under selective pressure before it was inactivated and neutrally after being inactivated (i.e., it should accumulate synonymous and non-synonymous mutations at an equal rate). The *K*_a_/*K*_s_ value (*K*) estimated for this entire branch is considered the average of the *K*_a_/*K*_s_ value for the part of the branch where the gene was under selection (*K*_s_) and the *K*_a_/*K*_s_ value for the part of the branch where the gene evolved neutrally (*K*_n_=1), weighted by the proportion of time for which the gene evolved under selection (*T*_s_/*T*) and neutrally (*T*_n_/*T*): *K* = *K*_s_×*T*_s_/*T*+*K*_n_×*T*_n_/*T*, where *T* represents the time since the split from the last common ancestor. Using the lower and upper bound of the confidence interval for the species divergence time *T* obtained from TimeTree ^173^ and *K*_a_/*K*_s_ value for afrotherians with a functional gene, one can calculate a lower and upper bound for T_n_ as *T*_n_=*T*×(*K*-*K*_s_)/(1-*K*_s_) to estimate how long a gene has evolved neutrally. A limitation of this method is that sirenian gene loss cannot be dated if a gene of interest is missing or heavily fragmented in any non-sirenian species within our afrotherian dataset.

### Sirenian lineage-specific amino acid changes

Amino acid alignments of the 7,695 single-copy gene family orthologs in our nine-species data set (eight afrotherians and Linnaeus’s two-toed sloth) and FasParser v2 ^186,187^ (https://github.com/Sun-Yanbo/FasParser) was used to identify amino acid residues specific to the sirenian lineage. We employed KOBAS v3.0 ^188^ (http://kobas.cbi.pku.edu.cn) gene enrichment and literature searches to identify genes that may underlie aquatic specializations of sirenians. Putative sirenian-specific residues of interest were further validated using 120 mammalian sequences downloaded from OrthoMaM v10c ^9^ (available at FigShare (https://figshare.com/s/16ca82c4e5f0a9f97161), as well as by interrogating NCBI and Ensembl databases using BLAST. Potential functional effects of substitutions were predicted by PolyPhen-2 ^189,190^ and SIFT ^191,192^. PolyPhen-2 predicts the possible impact of amino acid substitutions on the stability and function of human proteins using structural and comparative evolutionary considerations. SIFT predicts the potential impact of amino acid substitutions or indels on protein function.

### NIS radioiodide uptake assay

DNA sequences containing the protein-coding region of dugong, manatee, and human *SLC5A5* (NIS), as well as dugong (L142M, G203W, A321P, A445V, and S539T) and human (M142L, W203G, P321A, V445A, and T539S) sequences where five amino acid substitutions were changed to their reciprocal residues, were synthesized by GenScript Biotech Corp. (Nanjing, China). Each DNA was individually subcloned into the *pcDNA3.1* vector (Invitrogen) to generate ‘NIS plasmids’. All constructs were sequenced to verify their nucleotide sequences. HEK293T cells were cultured in 6-well plates until reaching 50-60% confluency. Cells were transfected with 2 μg NIS plasmid. Empty *pcDNA3.1* vector was used as a control. One day after transfection, HEK293T cells were seeded into 24-well plates and treated with radiopharmaceuticals after 24 hours. Briefly, cells were incubated with Na^125^I (Shanghai Xinke Pharmaceutical, Shanghai, China) at 37 [with 5% CO_2_ for one hour. The Na^125^I added to each well of the culture plate was counted using a radioactivity meter (FJ-391A4) to obtain the total radioactive count T (μCi) per well. After discarding the radioactive supernatant, cells were washed twice with PBS solution. Radioactive Na^125^I absorbed by the HEK293T cells was denoted the cell radioactive count C (μCi). The radioiodide uptake rate is shown as C/T%.

### Coimmunoprecipitation and immunoblotting of PER2 and CRY1

DNA sequences containing the protein-coding region of dugong, Asian elephant, and human *PER2* and *CRY1* were synthesized by GenScript Biotech. In addition, a human PER2 sequence with a sirenian-specific proline at residue 1220 (C1220) was synthesized. The CRY1 sequences contained a C-terminal 3×FLAG tag, the *PER2* sequences a C-terminal human influenza hemagglutinin (HA) tag. Each of the seven DNA sequences was individually subcloned into the *pcDNA3.1* vector (Invitrogen) and sequenced to confirm their identity. HEK293T cells were cultured in 10-cm Petri dishes until reaching 70-80% confluency and then transfected with 4 μg *CRY1-3×FLAG-pcDNA3.1* and 6 μg *PER2-3×HA-pcDNA3.1*. Two days after transfection, cells were lysed in Western IP Cell Lysis buffer (Beyotime, Nantong, China) supplemented with 1 mM PMSF (Biosharp) and subjected to coimmunoprecipitation. In total, 10% of the cell extracts were retained for input. Cell lysates were incubated with Anti-FLAG M2 Magnetic Beads (Sigma) at 4 °C overnight. After washing three times, the precipitates were resuspended in SDS–PAGE sample buffer, boiled for 5 min, and run on a 6% SDS–PAGE gel. Immunoblotting was done with mouse monoclonal anti-HA or anti-Flag antibodies (Proteintech).

### Whole-genome resequencing

Sequencing data from 99 dugongs sampled on the Australian east coast (see ‘Sample collection and research ethics’) were sequenced on a DNBSEQ-G400 RS instrument by BGI-Australia to generate 100-bp paired end reads. We also obtained public sequencing data from Okinawa (Japan; DRR251525; sampled 17 November 2019), Coogee Beach (New South Wales, Australia; ERR5621402; sampled 25 November 2009) ^77^, and Exmouth Gulf (Western Australia, Australia; SRR17870680; sampled 3 June 2018). We also obtained 101-bp paired-end WGS reads generated on the Illumina HiSeq 2000 platform (NCBI SRA SRR331137, SRR331139, and SRR331142) from a female West Indian manatee (Lorelei, born in captivity in Florida, USA) ^18^. The manatee reads were employed as outgroups in population genomics analyses. Raw data were filtered using SOAPnuke v2.1.5 ^124^ to remove adapters and low-quality reads. For comparative analyses, all samples were down-sampled to ∼10× coverage using SAMtools v1.9 ^149^

Because we had a chromosome-level genome of the dugong, we assigned sex to samples by using the Rx ratio method, where sequencing reads are mapped to a genome with an assigned X chromosome and the number of reads mapping to autosomes are compared to the X-chromosome (normalized by chromosome length) ^193,194^. The Rx ratio approximates 1.0 for females and 0.5 for males. Briefly, we first identified the X chromosome of the West Indian manatee (assembly TriManLat1.0_HiC; chromosome-level genome based on an assembly reported by ^18^) and dugong (assembly Dugong_BGI) by BLAST searches using the coding sequences of genes evenly distributed across the X chromosome of the African savanna elephant (*Loxodonta africana*) ^132^ and dugong Y chromosome genes ^133^. This effort revealed that the ∼167 Mb HiC_scaffold_7 and the ∼149 Mb chr7 in West Indian manatee and dugong corresponds to their respective X chromosome. Next, we aligned reads (the reads generated prior to the X chromosome-removal step above) to the chromosome-level genomes using bowtie2 v2.3.4.3 ^195,196^ (parameter: --no-unal to only retained mapping reads), followed by conversion to a BAM file and filtering using SAMtools v1.7 ^149,197^ (removal of PCR duplicate and retaining reads with a quality score, Q, above 30). Index statistics for BAM files were generated using idxstats in SAMtools and parsed by modifying an R script available at ^198^. Average sequencing depth was estimated from indexed and sorted BAM files after reads using mosdepth v0.3.3 ^199^ (parameters: -n --fast-mode --by 500). See **Table S12** for sample statistics.

### Identification and characterization of SNPs

Including the X chromosome can interfere with demographic history estimates ^90,200,201^ and other population genomics analyses ^202,203^ (X chromosome SNP effects). Therefore, reads that could be mapped to the X chromosome were removed using bowtie2 (v2-2.2.5) ^196^ with default parameters, resulting in ∼34.14 Tb of clean data with an average sequencing depth of around 11.41-fold. The filtered clean reads were aligned to our Dugong_BGI reference genome using BWA (v0.7.12-r1039) ^204^ with default parameters. SAMtools (v1.2) ^205^ was employed to convert SAM files to BAM format and to sort alignments, followed by the Picard package (v1.54; https://broadinstitute.github.io/picard) for duplicate removal. Next, GATK (v4.1.2.0) ^206^ was utilized to realign reads around InDels and detect SNPs. Briefly, we obtained the genomic variant call format (GVCF) in ERC mode based on read mapping with the parameters ‘-T HaplotypeCaller, -stand-call-conf 30.0 -ERC GVCF’. Joint variant calling was then conducted with the GATK *CombineGVCFs* module. Lastly, the GATK’s *VariantFiltration* module was used for hard filtering with the parameters ‘--filter-name LowQualFilter --filterExpression QD < 2.0 || MQ < 40.0 || FS > 60.0 || ReadPosRankSum < -8.0 || MQRankSum < -12.5 || SOR > 3.0’, as recommended by GATK. This process generated 61,741,769 SNPs.

### Analysis of population structure

To quantify the genetic structure of dugong populations, we first carried out SNPs filtration using vcftools (v0.1.16) ^207^ with the parameters ‘-max-missing 0.95’. Plink (v1.90b6.6) ^208^ was used to perform SNP quality control with the parameters ‘-geno 0.1 --maf 0.01’. Linkage disequilibrium was also used as a criterion to filter SNPs for downstream analysis using Plink with the parameters ‘-indep-pairwise 50 5 0.2’. Next, ADMIXTURE (v1.3.0) ^209^, with the parameters ‘--cv -j20 -B5’ for multiple repeats, was used to perform ancestry inference. To construct the population evolutionary tree, an identity by state (IBS) distance matrix was constructed using Plink with the parameters ‘--distance 1-ibs’. MEGA7 ^210^ was used to construct the Neighbor-Joining Tree (NJ tree) based on the IBS matrix, and the evolutionary tree was visualized using the iTOL online tool ^211^. In addition, PCA (principal component analysis) was performed using Plink with the parameters ‘--make-rel --pca 3’ and visualized by the R ggplot2 package ^212^.

### Estimation of genome heterozygosity and runs of homozygosity

To estimate the heterozygosity of each dugong individual, we used the Plink function ‘—het’ to detect heterozygous SNPs from the final SNP dataset. Additionally, we estimated runs of homozygosity (ROH) using the R package *detectRUNS* ^213^ with the parameters ‘windowSize = 50, minSNP = 30, maxGap= 1000000, minLengthBps = 100000, minDensity = 1/100000’ based on the same filtered SNPs set used in the ‘population structure analysis’ section.

ROH’s per dugong generation was calculated as follows. FASTEPRR (v2) ^214^ was used to estimate the recombination rate of the dugong population based on the 99 Queensland individuals. It was estimated to be ∼1.08cM/Mb. The coalescent times of ROH for each subpopulation were calculated as *g* = 100/(2rL) (see ^215^), where *g* is the expected time (in generations; 27 years ^67^) back to the parental common ancestor, *r* is the recombination rate and *L* is the length of the ROH in megabases. Thus, ROHs <100kb in length are estimated to result from inbreeding more than 12,501 years ago; ROHs>100 kb, up to 12,501 years ago (463 generations); ROHs>1 Mb, up to 1,242 years ago (46 generations); ROHs>5 Mb, up to 243 years ago (9 generations); and ROHs>10 Mb, up to 135 years ago (5 generations).

### Identification of selective sweep regions

Four different methods, including *Fst* (population differentiation), π (relative nucleotide diversity), *XP-EHH* (the cross-population extended haplotype homozygosity statistic) ^96^, and *XP-CLR* (the cross-population composite likelihood ratio test) ^97^ were employed to assess selective sweeps between the northern and southern groups. Briefly, vcftools (v0.1.16)^207^ with the parameters ‘-max-missing 0.95 --maf 0.01’ was used to perform autosome SNPs quality control. Next, the pairwise *Fst* was calculated between the seven groups and between the whole northern and southern groups using vcftools with the parameter ‘--fst-window-size 10000 --fst-window-step 2000’. π was calculated for each of the seven groups and the whole northern and southern groups using vcftools with the parameters ‘--window-pi 10000 --window-pi-step 2000’. XP-CLR scores were calculated using xpclr v1.1.2 ^216^ with default parameters. Because XP-EHH requires the genetic distance between adjacent SNPs, we considered a chromosome segment of 1 Mb to be 1 cM. The filtered SNPs were phased using ‘vcf_phase.py’, with the parameter “--phase-algorithm beagle” from the PPP (v0.1.12) pipeline ^217^, followed by vcftools with the parameter ‘—IMPUTE’. Next, the *xpehhbin* module from Hapbin (v1.3.0) ^218^ was used to calculate XP-EHH values (see ^219^). The XP-EHH scores were normalized and corresponding *P* values were calculated. If a *P* value is less than 0.01, we considered the region as a candidate sweep region. An XP-EHH score is directional: a positive score suggests that selection occurred in the northern group; a negative score, the southern group. Genomic regions that overlapped between all four methods were considered candidate selective sweep regions.

### Demographic history inference

The generation time (*g*) and mutation (μ) rate are necessary to infer the demographic history of populations. The estimated generation time of the dugong (∼27 years) has been reported previously ^67^. We estimated the mutation rate for dugong using r8s ^220^ and the single copy orthologous genes described in the ‘Phylogeny and divergence time estimation’ section. The final estimated mutation rate was 2.6×10^-8^ per site per generation.

The pairwise sequentially Markovian coalescent method (PSMC) v0.6.5-r67 ^221^ was employed to infer historical effective population size (*Ne*) fluctuations in dugongs. PSMC can quantitively reveal changes in *N_e_* from approximately 1 million to 20 thousand years ago ^90^. We first constructed diploid consensus sequences for each sample using SAMtools (v1.9) ^205^ mpileup and BCFtools (v1.4) ^222^ with the parameters ‘-C50’ and ‘-d 4 -D 24’. The consensus sequences were transformed to PSMC input format using fq2psmcfa with the parameter ‘-q20’. Finally, PSMC was used to infer the population history with the parameters ‘-N25 -t15 -r5 -p 4 + 25∗2 + 4+6’ and 100 rounds of bootstrapping. SMC++ (v1.15.5) ^223^ was used to infer more recent population history for Queensland dugong individuals grouped into two distinct populations (see ‘Analysis of population structure’ above). The SMC++ modules *vcf2smc*, *estimate*, and *plot* were used.

## Data availability

The data for *D*. *dugon* including stLFR data, Hi-C data, RNA-seq data, resequencing data, genome assembly, and gene annotation in this study has been deposited at the China National GeneBank Nucleotide Sequence Archive (CNSA: https://db.cngb.org/cnsa) under accession number CNP0004405. Multiple sequence alignments (MSAs) of thyroid hormone pathway and circadian clock genes with sirenian-specific amino acid substitutions are available on FigShare (https://figshare.com/s/16ca82c4e5f0a9f97161).

## Acknowledgements

We thank those who assisted with Australian dugong tissue collection, including The University of Queensland Dugong Team (especially Alex McGowan, Erin Neal, and Rob Slade) members of the Mabuiag Island community of Torres Strait (especially Terrence Whap), and Steve Hoseck (Southern Marine Parks, Queensland Parks and Wildlife) for facilitating access to the dugong fetus used for stLFR and RNA-sequencing. We also thank the laboratory of Prof. Kai Yang at Soochow University (China) for their radioisotope expertise and access to equipment, and Dr. Erina Young at Murdoch University (Australia) for information on the dugong individual from Exmouth Gulf, Western Australia.

Support for this research was provided by the National Natural Science Foundation of China (grant 32270441 to R.T. and grant 42225604 to S.L.) National Key Programme of Research and Development, Ministry of Science and Technology (grant 2022YFF1301601 to R.T.); Sea World Foundation (Australia) and Winifred Violet Scott Foundation grants (to J.M.L.); the ‘One Belt and One Road’ Science and Technology Co-operation Special Program of the International Partnership Program of the Chinese Academy of Sciences (grant 183446KYSB20200016 to S.L.); the Specially-appointed Professor Program of Jiangsu Province (to I.S.), the Jiangsu Foreign Expert Bureau (to I.S.), and the Jiangsu Provincial Department of Technology (grant JSSCTD202142 to I.S.).

## Author contributions

R.T., J.M.L., G.F., S.L., and I.S. conceived the study; Y.Z., P.Z., J.M.L., H.L.S. collected or curated samples/specimens; R.T., Z.J., and L.W. performed laboratory work; R.T., Y.Z., H.K., F.Z., J.W., and I.S. performed computational biology work; R.T., S.L., G.F., and I.S. managed the project; R.T. and I.S. wrote the original draft; all authors commented on and proof-read the manuscript, with significant contributions from J.M.L., G.F., and S.L.

