## Supplementary information for "Sirenian genomes illuminate the evolution of fully aquatic species within the mammalian superorder Afrotheria"

### SUPPLEMENTARY NOTES

**Supplementary Note 1. Generation of a chromosome-level dugong reference genome**

We produced a 3.06 Gb chromosome-level reference assembly (Ddugon_BGI) with the expected [1] 25 pairs of chromosomes ($2n$=50) (**Figure S1**) from a female dugong by combining stLFR and Hi-C sequencing data (**Table S1**), leaving 13.35 Mb (0.44%) unplaced scaffolds. We annotated 18,663 protein-coding genes (**Table S1**) in our assembly.

There are currently (June 2023) four publicly available dugong assemblies. These are a short-read sequencing assembly generated by the Japanese National Institute for Environmental Studies (NCBI Assembly GCA_015147995.1), a synthetic long-read assembly generated using 10x Genomics Chromium technology (hereafter: Ddugon_MaxPlanck) [2], a chromosome-level assembly generated by the DNA Zoo consortium after scaffolding an in-house short-read sequencing assembly (hereafter, Ddugon_DNAzoo), and a very recent (late May 2023) assembly from the Vertebrate Genome Project (VGP) [3] (NCBI Assembly GCA_030035595.1).

Two West Indian manatee (*Trichechus manatus*) genomes are available (**Table S2**). An assembly generated by Foote and colleagues in 2015 [4] recently scaffolded into a chromosome-level assembly by the DNA Zoo consortium (scaffold and contig N50 143.72 Mb and 37.75 kb) was employed in this manuscript. A draft genome [5] (H_Gigas_1.0 in **Table S2**) and a pseudo-genome [2] (generated by mapping short reads to assembly Ddugon_MaxPlanck) of the extinct cold-adapted Steller's sea cow (*Hydrodamalis gigas*) are available but are derived from centuries-old DNA samples and thus highly fragmented, and were not employed in the phylogenomic and genome-scale comparative analyses in this manuscript.

**Supplementary Note 2. The enigmatic phylogeny of paenungulates**

It is accepted that the closest living relatives to sirenians are the other afrotherian herbivores, the terrestrial proboscideans (elephantids) and hyracoids (hyraxes). They are grouped within the superorder Paenungulata ('almost ungulates' in Latin; now a misnomer as afrotherians are distantly related to ungulates). Their internal topology remains debated. Some studies group proboscideans and sirenians into Tethytheria to suggest a shared origin near the Tethys Sea between Africa and Eurasia before the continents connected, others group hyracoids and proboscideans, or hyracoids and sirenians [6-9]. Morphology (large size and near-hairlessness) may suggest that proboscideans and sirenians should be exclusively grouped. However, these features may mask the true relationships: although extant hyracoids are small (body weight <5kg), their ancestors had a body mass similar to their contemporary proboscideans and sirenians [10]. Furthermore, while it is not known if the shared ancestor of furred modern hyraxes and near-hairless elephants and sirenians had abundant body hair or not, wooly mammoths illustrate that gain of fur from a near-hairless ancestor can be achieved by paenungulates [11,12].

Our phylogenetic analyses based on nuclear DNA (coding regions, 4-fold degenerate sites, and the third codon positions 7,695 protein-coding genes; 5,508 single-copy BUSCO genes; and 627,279 conserved non-exonic elements) supported a sister group relationship between Sirenia with Hyracoidea + Proboscidea (**Figure 1c** and **Figure S2**). This phylogeny agrees with one generated from ultraconserved sites evolving neutrally (or near-neutrally) in coding and non-coding genome regions of 242 placental mammals, including the West Indian manatee, rock hyrax, and African savanna elephant (*Loxodonta africana*) [13]. However, despite strong support, our tree is not necessarily correct. Because paenungulates radiated within a relatively short time, it has been reasoned [e.g., see [14]] that ancient paenungulates were subjected to incomplete lineage sorting (ILS) and introgressive hybridization (also known as introgression). These processes can blur species boundaries and introduce phylogenetic noise. To address this problem, intragenic indels and retroelements (LINEs, SINEs, and LTRs) have been championed as near homoplasy-free phylogenetic markers since there is a low probability that they arose in two species independently and their gain/loss is usually permanent (retroelements, in particular, given their mode of insertion) [15-18]. A recent study [19] considered whole-genome sequences and ≥50-bp intragenic indels and provided support for proboscideans + sirenians (406 indels) but also proboscideans + hyracoids (238 indels; i.e., consistent with our topology in **Figure 1c**) to indicate ILS and possibly also introgression between paenungulates, showing the ongoing difficulty in resolving their phylogeny. Using our chromosome-level dugong genome as the reference species, we could not reliably call retroelement markers in afrotherian genomes using a recently developed approach that requires 2 kb of sequence flanking both sides of a retrotransposon [17]. The genomes of afrotherian species are currently at various stages of completion and lack chromosome-level assemblies generated in concert with long-read or synthetic long-read data. Therefore, most assemblies examined here were highly fragmented and contain numerous short contigs (**Table S2**) that may not resolve many retroelements. However, near-complete genomes of all afrotherians will likely become available within the next few years, mainly via the Vertebrate Genome Project (VGP) [3]. Regardless of the true relationship between paenungulates, we provide a phylogenetic framework suitable for understanding the evolution of the only fully aquatic species within Afrotheria.

**Supplementary Note 3. Molecular evolution of the sirenian integumentary system**

The integumentary system, the skin and its appendages, is the largest organ of the mammalian body and a barrier to the external environment that maintains water balance and contributes to thermoregulation [20]. Recent studies highlight skin adaptations since the sirenian and elephantid ancestors diverged (~58 Mya; **Figure 1c**). Elephantid species acquired novel gene changes associated with skin modifications to life on the arid African continent [21] and in the Arctic [11]. Sirenians adapted their skin to life in water, with the extinct Steller's sea cow further evolving a larger body size, thicker blubber, and thick, rough skin [2] to adjust to the sub-Arctic environment. The integumentary system of manatees and the dugong is quite similar. Exceptions include their epidermis and blubber [22,23]. The manatee epidermis is structurally more similar to cetaceans (black in color), while the dugong epidermis is brown-gray. The dugong has a single-blubber layer (except for the centrum), while the manatee has two layers throughout the body [22,23] (**Figure S4**).

Previously reported sirenian epidermal gene loss events include type I keratins [24,25], arachidonic acid 15-lipoxygenase-1 (*ALOX15*) [26], and kallikrein 8 (*KLK8*; also expressed by sweat glands) [27]. We observed *TCHH1* and *TCHHL1* loss, genes highly expressed by the hair follicle and the hair shaft [28] that produce their structural component, trichohyalin. Our analysis and recent studies also identified sirenian loss of hair follicle-associated keratin genes, goose-type lysozyme 2 (*LYG2*), and acyl-CoA wax alcohol acyltransferase 2 (*AWAT2*) [24,25,29,30]. Shared morphological features of sirenians and cetaceans, the only fully aquatic mammals, include an absence of pelage (i.e., coat) hair follicles [31]. Sirenians are not completely hairless, however, but harbor sparsely distributed body hairs that likely serve a tactile function [31,32]. Not surprisingly, many of the same epidermal and dermal genes have been independently lost in cetaceans and sirenians: type I and II keratins (KRTs) [24,25], *ALOX5* [26], *LYG2* [30], *AWAT2* [29], *MCR5* [33,34], and *KLK8* [27]. Four genes identified in our analysis (*TCHH1L*, *AWAT15*, *KRT2*, and *KRT77*) are among eight independently lost in cetaceans and hippopotamuses (hippos), a semi-aquatic sister group to cetaceans [35]. Two recent studies demonstrated a loss of melanocortin receptor 5 (*MC5R*), a gene with sebaceous gland functions, in the West Indian manatee, cetaceans, and other species with no apparent sebaceous glands [33-35].

Most fat in marine mammals is found in their hypodermis (blubber), a skin layer rich in white adipose tissue (WAT) with an inner stratum of brown adipose tissue (BAT) [31,36,37]. The mammalian hypodermis also contains adipose-derived stem cells (ADSC) that give rise to adipocytes of the BAT and WAT. Brown adipocytes of BAT and beige adipocytes of WAT express uncoupling protein 1 (*UCP1*), a gene essential for non-shivering thermogenesis (NST) [38]. During cold exposure, thyroid hormones regulate thermogenesis directly via *UCP1* [39]. The blood levels of thyroid hormones in the West Indian manatee [40] and a tropical cetacean are similar, while the levels of cold-water cetaceans are approximately twice as high [41]. *UCP1* is reportedly inactivated in many terrestrial mammals, cetaceans, sirenians (including the cold-adapted Steller's sea cow), and some pinnipeds [42-45]. Numerous factors may explain why *UCP1* loss is tolerated in certain mammals. Prominent among these is that species with a large body size are inherently better at conserving heat [44,46]. *UCP1* may only be lost in northern and southern elephant seals (genus *Mirounga*), species with very large body sizes [47,48]. All pinnipeds also have fur, a much better insulator than blubber [46] that may somewhat reduce their reliance on UCP1-mediated thermogenesis. Gene loss may not always abolish function, however [49]. A recent study reported that, despite harboring several ostensibly inactivating mutations, *UCP1* is expressed by cetacean BAT and shows a reduced function that may serve to maintain a balance between maintaining a thick blubber and thermogenesis [36].

Within Afrotheria, inactivating *UCP1* mutations is not unique to sirenians. It is also found in hyraxes (their ancestor had a much larger body size [10]) and elephants, while the smaller species of their insectivorous sister clade (Afroinsectiphilia) have retained the gene and show evidence of *UCP1* neo-functionalization [43,44]. Thus, large-bodied paenungulates, which emerged on the warm African continent, likely later evolved cold-hardiness adaptations that do not rely on *UCP1*-mediated thermogenesis, as illustrated by the larger body size and thick, hyperkeratotic epidermis of the Steller's sea cow [2] and fur of woolly mammoths. While no blubber expression and functional data on *UCP1* is currently available for an extant sirenian, their thin blubber compared to cetaceans and pinnipeds [31], low metabolic rate (at least in manatees), and restriction to tropical and subtropical habitats suggests that the dugong and manatees are naturally susceptible to cold temperatures.

**Supplementary Note 4. Population structure and history of dugongs on the Queensland coast**

A very recent microsatellite study provided evidence of restricted gene flow and distinct Australian populations north and south of the Whitsunday Islands [50] (**Figure S8a**), although the timing of this event remained unknown. Given that our seven-population data set contained numerous individuals across the Queensland range, we further examined this apparent genetic break. Principal component analysis (PCA) (**Figure S8b**), a neighbor-joining (NJ) tree (**Figure S8c**), and ADMIXTURE analysis (**Figure S8d**) supported a grouping by latitude, north ($n$=11) and south ($n$=88) of the Whitsunday Islands. The mean population differentiation (*Fst*) between the northern and southern group was ~0.086 (**Figure S8e**), further agreed with the north-south grouping. Heterozygosity (see **Figure S9a**) was significantly higher in the northern Torres Strait population compared to all other populations (Dunn test with Holm correction, $P < 0.05$). We speculate that this northernmost dugong populations resumed panmixia with populations from the Indian Ocean once sea levels rose and covered the Torres Strait land bridge ~7 kya, removing an ecological barrier present for 100,000 years.

Members from the seven Queensland dugong locations carried a small number of >1Mbp runs of homozygosity (median five ROHs) (**Figure S9b,c** and **Table S14**). An exception was individuals from Airlie Beach Whitsunday Islands, the location of the apparent genetic break border. Only three individuals were obtained from this location; sampling additional individuals is necessary before speculating on inbreeding in this population. Nevertheless, the inbreeding coefficient ($F_{ROH}$ >1Mbp) was very low across all Queensland dugongs (median $F_{ROH>1Mbp} = 0.0023$), including the outlier individual AB16008 ($F_{ROH>1Mbp} = 0.015$), and did not differ significantly between populations (**Figure S9d**) (Kruskal-Wallis $P = 0.25$).

Because of the insufficient resolution of PSMC in estimating demography more recently than ~20 kya [8], we used SMC++ [51]. One of the 32 individuals sampled from Moreton Bay (MB16796; an older female) clustered with the northern group in a PCA and a neighbor-joining tree (**Figure S8b,c**), which may reflect low-level individualistic movement rather than population migration (see [50,52,53]) between northern and southern populations, was excluded from the demographic history analysis. We observed similar $N_e$ trajectories until ~10 kya (**Figure S6d**), in agreement with the PSMC result (**Figure 6c**). The divergence time between the southern and northern group was estimated as ~10.7 kya (95% CI: 9.1-12.2 kya) using *dadi* [54]. A larger effective population size was observed for the northern group from ~3 to 1 kya ($N_e$ of 80,000 individuals compared to 10,000 in the southern group). The smaller recent effective population size of dugong populations south of this barrier may benefit from assignment as distinct conservation management units. While the cause and nature of the ecological barrier between the two population groups at the Whitsunday Islands have yet to be determined [50], there is a high degree of geographic variability in water currents around the reef-dense Whitsunday Islands [55].

### SUPPLEMENTARY FIGURES

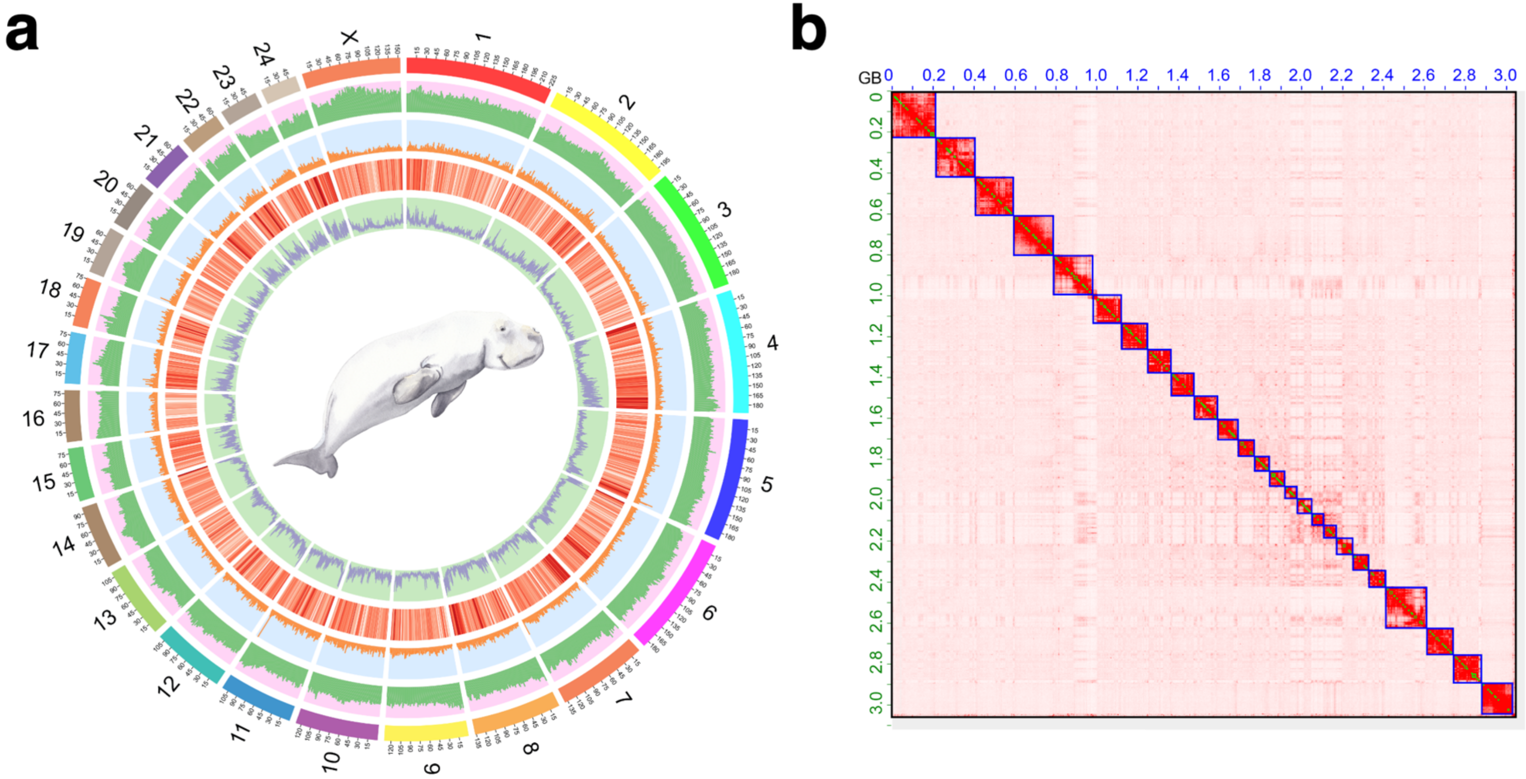

**Figure S1 | Overview of dugon reference assembly Ddugon_BGI**

**a,** Circos plot of *Dugong dugon* assembly Ddugon_BGI. The outermost segment represents chromosome sequences, with the numbers on the external surface indicating genome size in Mb. The line plots, from outside to inside, respectively represent the distribution of retrotransposon ratio (from 0.02 to 0.82), DNA transposon ratio (from 0 to 0.08), gene density (from 0 to 54) at 1Mb windows and GC content (from 0.33 to 0.64) at 100 kb windows. **b,** Genome-wide Hi-C contact matrix of Ddugon_BGI. The heat map shows long-range contacts and scaffolding of the genome assembly. Blue squares denote chromosomes.

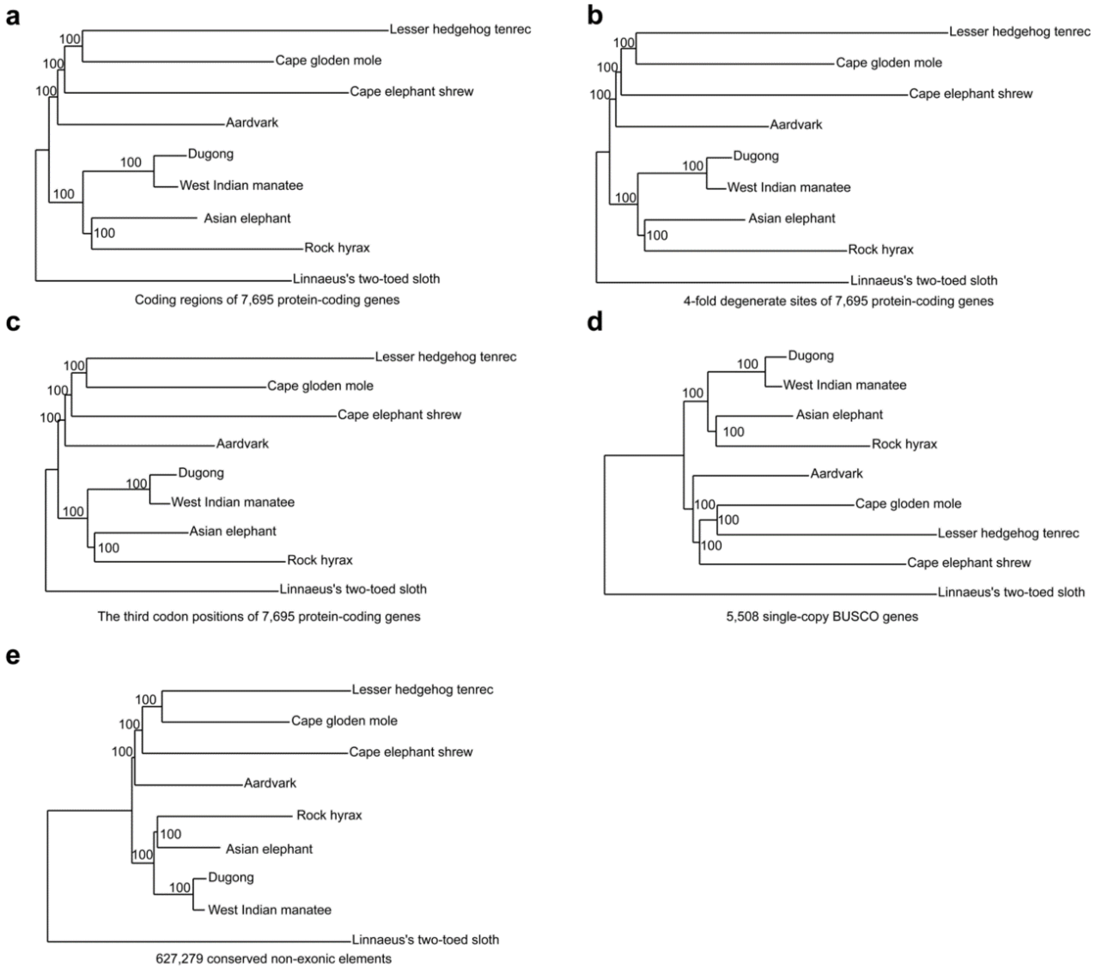

**Figure S2 | ML phylogenetic trees of afrotherians inferred using nuclear DNA**

**a,** coding regions of 7,695 protein-coding genes **b,** 4-fold degenerate sites of 7,695 protein-coding genes. **c,** the third codon positions of 7,695 protein-coding genes. **d,** 5,508 single-copy BUSCO genes. **e,** 627,279 conserved non-exonic elements (CNEEs). Trees were concatenated into a single super sequence and a ML phylogenetic tree construed using RAxML. 1,000 bootstrap replicates were performed to compute the node support (100 represents 100% support).

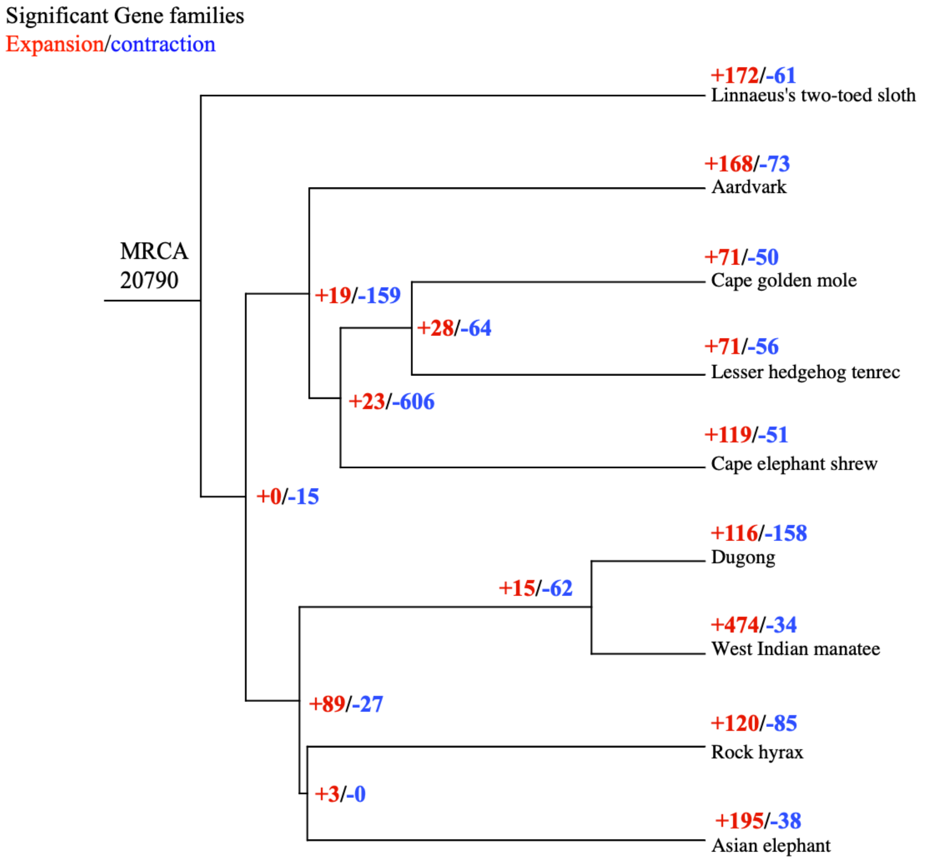

**Figure S3 | Overview of gene families expanded and contracted in afrotherians**

A maximum likelihood tree based on 7,695 orthologous proteins was generated and used to infer expansion and contraction of 20,790 gene families. The number of expanded and contracted gene families is in blue and red, respectively. MRCA, most recent common ancestor. Linnaeus's two-toed sloth was included as an outgroup.

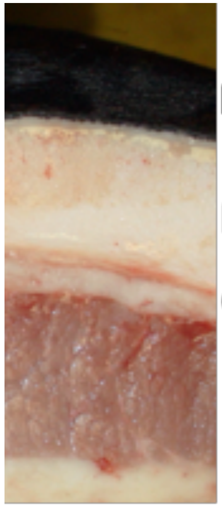

**Figure S4 | Histology of West Indian manatee flank skin**

Note the thin epidermis (black) and thick dermis. Photo by Michael Lusk (URL: https://www.flickr.com/photos/killkudzu/7619052292) under CC BY-SA 2.0.

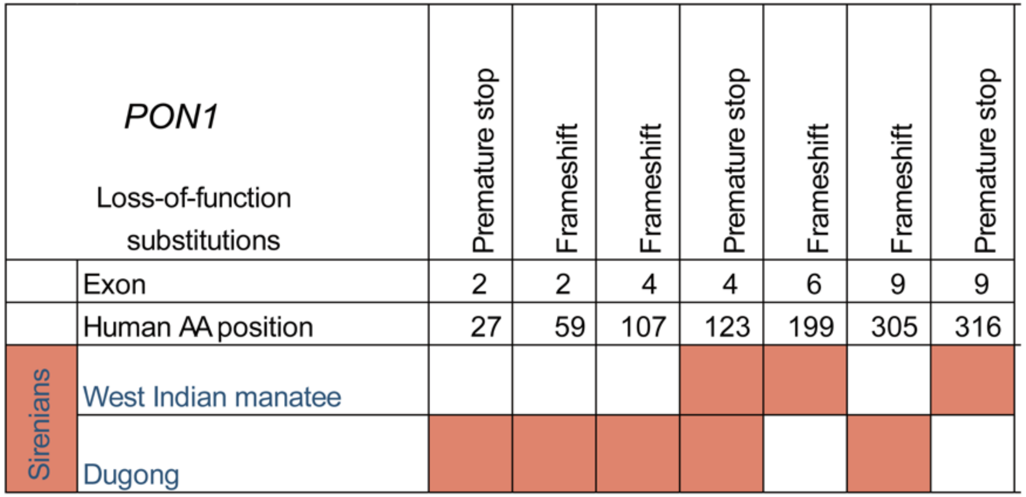

**Figure S5 | Overview of gene inactivation substitutions in sirenian paraoxonase (*PON1*)**

Colored cells indicate inactivating events. Loss of function substitutions include exon deletions, premature stop codons, and frameshifts.

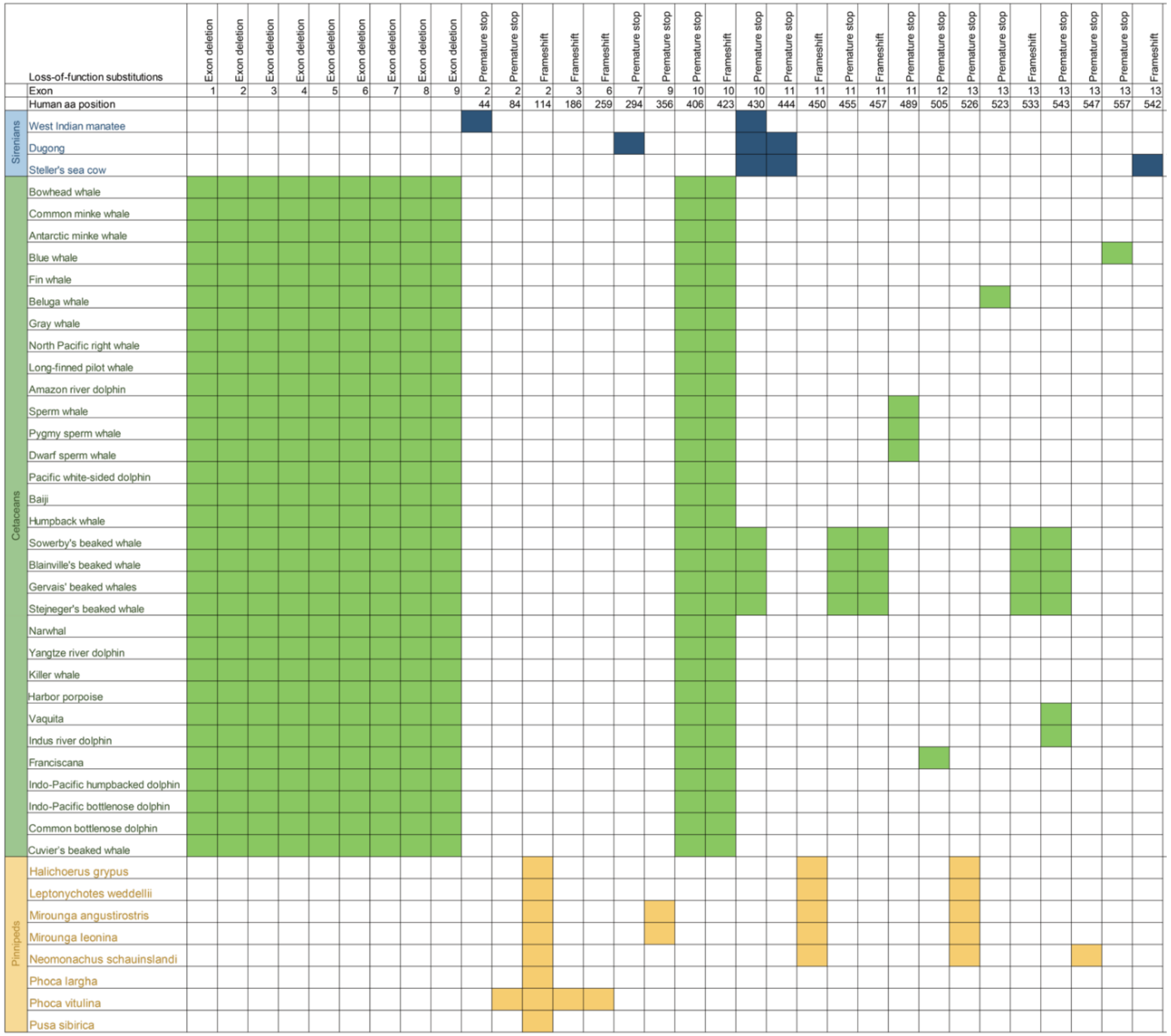

**Figure S6 | Overview of carboxylesterase 3 (*CES3*) gene inactivation substitutions in sirenians, cetaceans, and pinniped family Phocidae**

Colored cells indicate inactivating events. Loss of function substitutions include exon deletions, premature stop codons, and frameshifts.

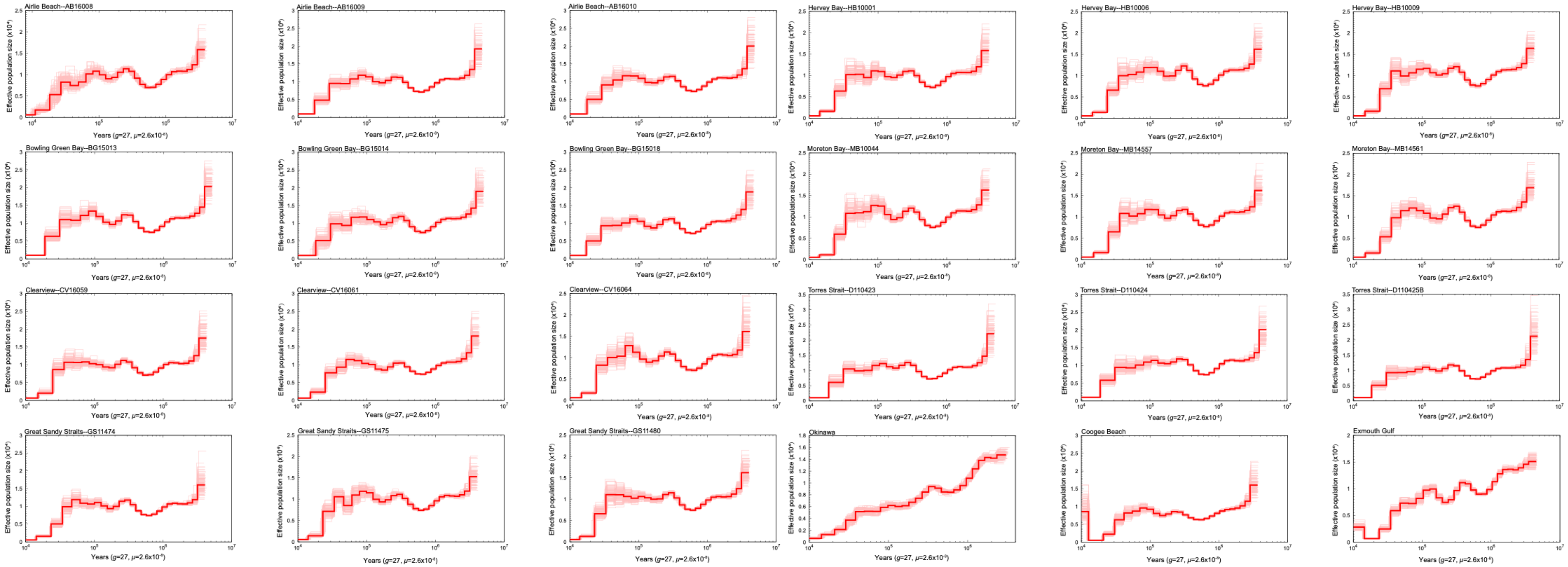

**Figure S7 | Population size history of dugongs**

Historical population sizes ($N_e$) were estimated using the pairwise sequentially Markovian coalescent (PSMC) method and diploid genome sequences (autosomes-only). The *y*-axis shows $N_e$, with 100 bootstrap estimates indicated by lighter lines. Plots were scaled using a mutation rate ($\mu$) of $2.60 \times 10^{-8}$ substitutions per nucleotide per generation and species-specific generation times ($g$) of 27 years. Three random individuals were selected from the Queensland locations Torres Strait, Bowling Green Bay, Airlie Beach, Clairview, Hervey Bay, Great Sandy Strait, and Moreton Bay.

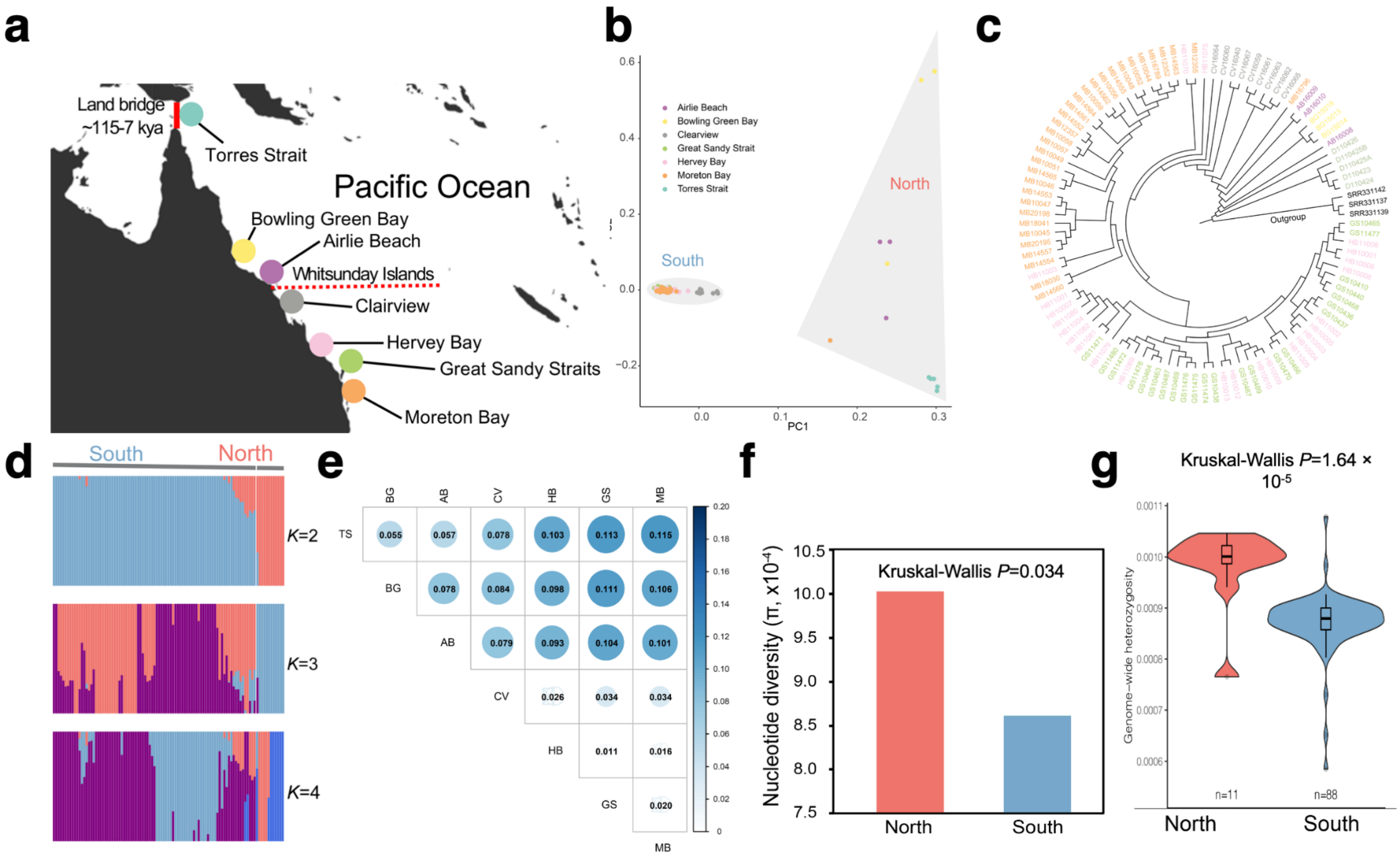
a
Land bridge
~115-7 kya
Torres Strait
Pacific Ocean
Bowling Green Bay
Airlie Beach
Whitsunday Islands
Clairview
Hervey Bay
Great Sandy Straits
Moreton Bay
b
Airlie Beach
Bowling Green Bay
Clearview
Great Sandy Strait
Hervey Bay
Moreton Bay
Torres Strait
South
North
PC1
c
Outgroup
d
South
North
K=2
K=3
K=4
e
TS
BG
AB
CV
HB
GS
MB
f
Kruskal-Wallis P=0.034
Nucleotide diversity (π, x10⁻⁴)
North
South
g
Kruskal-Wallis P=1.64 × 10⁻⁵
Genome-wide heterozygosity
n=11
n=88
North
South

**Figure S8 | Evidence of population structure by dugongs on the Queensland coast**

**a,** Map indicating approximate sites of origin for dugong samples from the Queensland coast, eastern Australia. A Torres Strait land border (115–7 kya) is indicated by a continous line; the dotted red line, a contemporary ecological barrier. The map was generated using the R package 'OpenStreetMap' [56]. **b,** Principal component (PC) analysis, a neighbor-joining (NJ) tree, and **d,** structure (ADMIXTURE) plots demonstrate substantial structure among seven dugong populations Australian east coast, supporting a break between population north and south of the Whitsunday Islands. Torres Strait individuals in the NJ tree (c) have a 'D' prefix. The West Indian manatee was used as an outgroup (black font) in the NJ tree. **e,** Pairwise population differentiation index (*Fst*) differences between northern (TS, BG, and AB) and southern (CV, HB, GS, and MB) population groups. Calculated in non-overlapping 10-kb sliding windows with 2-kb step sizes. **f,** relative nucleotide diversity ($\pi$) of northern (TS, BG, and AB) and southern (CV, HB, GS, and MB) population groups. Statistics performed on the mean $\pi$ values of the seven locations. **g,** Genome-wide heterozygosity of northern (TS, BG, and AB) and southern (CV, HB, GS, and MB) population groups. Statistics performed on heterozygosity values of 99 individuals.

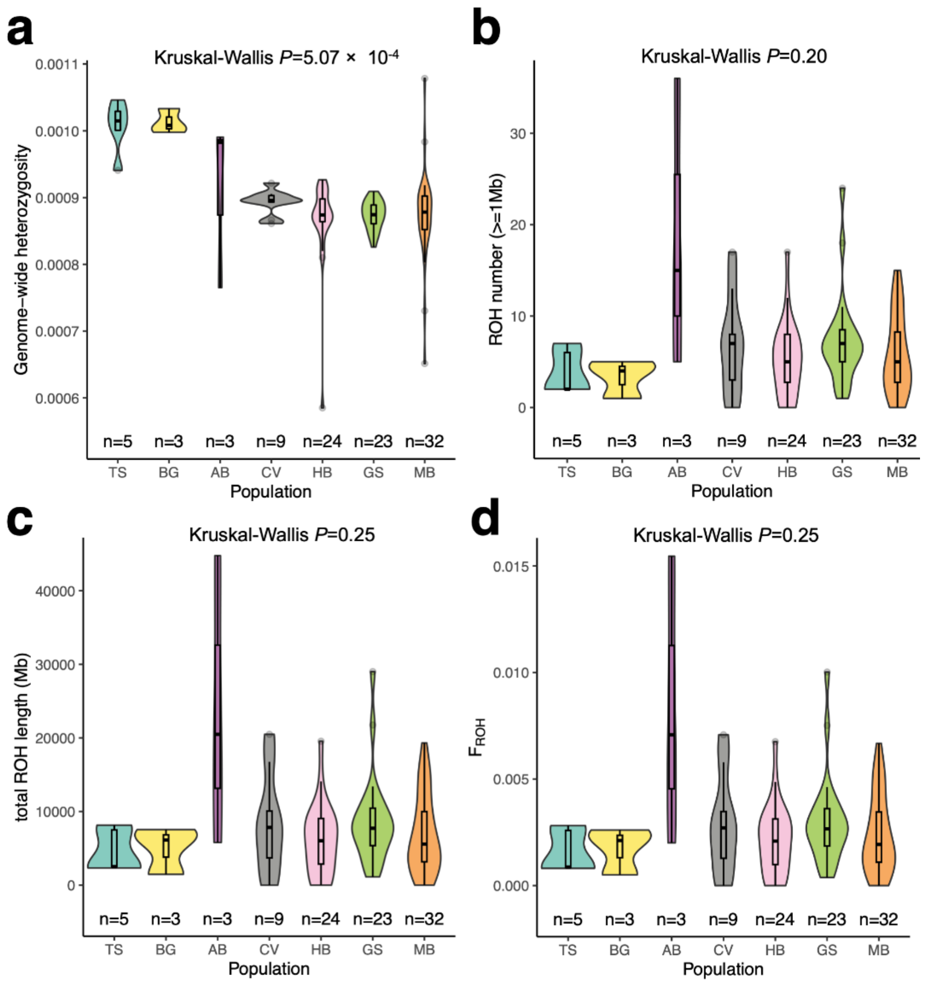

**Figure S9 | Population diversity of dugong populations on the Queensland coast**

**a,** Genome-wide heterozygosity. **b,** Number of runs of homozygosity (ROH) larger than 1 Mb. TS denotes Torres Strait, QLD; BG, Bowling Green Bay, QLD; AB, Airlie Beach, QLD; CV, Clairview, QLD; HB, Hervey Bay, QLD; GS, Great Sandy Strait, QLD; MB, Moreton Bay, QLD. **c,** Total lengths of ROHs. **d,** The proportion of the autosomal genome in ROH ($F_{ROH}$ >1Mbp).

### SUPPLEMENTARY TABLES

#### Table S1 | Summary of Ddugon_BGI sequencing and annotation

Hi-C anchored rate refers the proportion of scaffolded bases assembled onto 25 pseudochromosomes (chr 24 autosomes + chr X).

| | | |
|---|---|---|
| **Genome assembly** | **Estimated genome size** | **3.43 Gb** |
| | Assembly size (scaffold) | 3.06 Gb |
| | Assembly size (contig) | 2.99 Gb |
| | Hi-C anchored rate | 99.56% |
| | Contig number | 26,850 |
| | Contig N50 | 226.99 kb |
| | Longest contig | 2.29 Mb |
| | Scaffold number | 2,926 |
| | Scaffold N50 | 136.91 Mb |
| | Longest scaffold | 228.81 Mb |
| | GC content | 40.58% |
| | Gaps (N) | 2.27% |
| **Transposable elements** | **Annotation** | **Percent** |
| | DNA | 1.23% |
| | LINE | 44.92% |
| | SINE | 3.41 % |
| | LTR | 10.44 % |
| | Unknown | 0.0030 % |
| | Total | 49.200% |
| **Protein-coding genes** | Predicted genes | 19,897 |
| | Average transcript length | 63,305.31 bp |
| | Average coding sequence length | 1,649.23 bp |
| | Average exon length | 168.60 bp |
| | Average intron length | 7,020.79 bp |
| | Functionally annotated genes | 18,663 |

**Table S2 | Comparison of assembly quality between afrotherians genome assemblies**

Ddugon_BGI denotes the assembly generated in our study. Assemblies with the suffix _HiC were generated by DNA Zoo, the remainder are assemblies are available via NCBI. Assemblies GCA_905400935.1, GCA_015147995.1 and Dugong_dugon_HiC are denoted 'Ddugon_MaxPlanck', 'Ddugon_NIES' and 'Ddugon_DNAzoo)' elsewhere in this manuscript. The DNA Zoo assemblies can be found at dnazoo.org/assemblies/Dugong_dugon (Ddugon_HiC), dnazoo.org/assemblies/Trichechus_manatus (TriManLat1.0_HiC), dnazoo.org/assemblies/Elephas_maximus (Elephas_maximus_HiC), dnazoo.org/assemblies/Procavia_capensis (Pcap_2.0_HiC), dnazoo.org/assemblies/Orycteropus_afer (OryAfe1.0_HiC). Please note that the Steller's sea cow (*Hydrodamalis gigas*) was assembled using short-insert reads from ancient DNA samples and is partial assembly of a genome approximately the same size as the dugong and manatee (~3 Gb).

| Latin name | Common name | Assembly | Chromosome-level? | Scaffold genome size (Gb) | Contig genome size (Gb) | G+C content (%) | No. scaffolds | No. contigs | Scaffold N50 (Mb) | Contig N50 (kb) |
|---|---|---|---|---|---|---|---|---|---|---|
| *Dugong dugon* | Dugong | Ddugon_BGI | Yes | 3.06 | 2.99 | 40.58 | 2,926 | 26,850 | 136.91 | 226.99 |
| *Dugong dugon* | Dugong | Ddugon_HiC | Yes | 3.10 | 3.08 | 40.63 | 335,494 | 391,685 | 118.74 | 63.95 |
| *Dugong dugon* | Dugong | GCA_030035595.1 | Yes | 3.16 | 3.16 | 40.65 | 198 | 294 | 140.67 | 57,632.67 |
| *Dugong dugon* | Dugong | GCA_905400935.1 | No | 3.20 | 3.10 | 40.46 | 16,045 | 52,168 | 3.11 | 155.49 |
| *Dugong dugon* | Dugong | GCA_015147995.1 | No | 2.62 | 2.61 | 40.73 | 116,852 | 285,112 | 0.031 | 17.96 |
| *Hydrodamalis gigas* | Steller's sea cow | H_Gigas_1.0 | No | 1.24 | 1.24 | 44.13 | 998,083 | 1,091,214 | 0.0014 | 13.45 |
| *Trichechus manatus* | West Indian manatee | TriManLat1.0_HiC | Yes | 3.10 | 2.77 | 40.73 | 5,652 | 166,508 | 143.72 | 37.75 |
| *Trichechus manatus* | West Indian manatee | ASM3001377v1 | No | 3.09 | 3.09 | 40.66 | 15,608 | 15,641 | 0.49 | 486.87 |
| *Elephas maximus* | Asian elephant | Elephas_maximus_HiC | Yes | 3.21 | 3.19 | 40.83 | 543,483 | 602,016 | 95.96 | 57.61 |
| *Procavia capensis* | Rock hyrax | Pcap_2.0_HiC | Yes | 3.61 | 3.20 | 41.00 | 59,965 | 327,292 | 133.72 | 35.45 |
| *Orycteropus afer* | Aardvark | OryAfe1.0_HiC | Yes | 4.42 | 3.42 | 40.07 | 22,223 | 434,556 | 644.00 | 17.65 |
| *Echinops telfairi* | Lesser hedgehog tenrec) | ASM31398v2 | No | 2.95 | 2.61 | 43.01 | 7,490 | 277,896 | 54.42 | 20.43 |
| *Elephantulus edwardii* | Cape elephant shrew | EleEdw1.0 | No | 3.84 | 3.12 | 40.30 | 8,768 | 288,162 | 15.01 | 24.22 |
| *Chrysochloris asiatica* | Cape golden mole | ChrAsi1.0 | No | 4.21 | 3.36 | 40.01 | 20,500 | 391,344 | 13.47 | 19.63 |

### Table S3 | BUSCO evaluation of afrotherian genome assemblies

For consistency, all genome assemblies were examined using the same version and library of BUSCO (5.0.0_cv1 with the 9,226-gene mammalia_odb10 dataset). For additional assembly information, please see Table S2. Assemblies with the suffix _HiC were generated by DNA Zoo, the remainder are assemblies available via NCBI.

| Afrotheria family | Latin name | Common name | Assembly | Complete BUSCOs (C) | Complete and single-copy BUSCOs (S) | Complete and duplicated BUSCOs (D) | Fragmented BUSCOs (F) | Missing BUSCOs (M) |
|---|---|---|---|---|---|---|---|---|
| Dugongidae | *Dugong dugon* | Dugong | Ddugon_BGI | 8,712 (94.4%) | 8,645 | 67 | 156 | 358 |
| Dugongidae | *Dugong dugon* | Dugong | Ddugon_HiC | 8,637 (93.6%) | 8,564 | 73 | 195 | 394 |
| Dugongidae | *Dugong dugon* | Dugong | GCA_030035595.1 | 8,875 (96.2%) | 8,790 | 85 | 72 | 279 |
| Dugongidae | *Dugong dugon* | Dugong | GCA_905400935.1 | 7,996 (86.6%) | 7,900 | 96 | 460 | 770 |
| Dugongidae | *Dugong dugon* | Dugong | GCA_015147995.1 | 5,843 (63.3%) | 5,780 | 63 | 1,221 | 2,162 |
| Dugongidae | *Hydrodamalis gigas* | Steller's sea cow | H_Gigas_1.0 | 1,288 (14.0%) | 1,269 | 19 | 1,221 | 6,717 |
| Trichechidae | *Trichechus manatus* | West Indian manatee | TriManLat1.0_HiC | 8,682 (94.1%) | 8,607 | 75 | 174 | 370 |
| Trichechidae | *Trichechus manatus* | West Indian manatee | ASM3001377v1 | 8,490 (92.1%) | 8,410 | 80 | 285 | 451 |
| Elephantidae | *Elephas maximus* | Asian elephant | Elephas_maximus_HiC | 8,483 (91.9%) | 8,407 | 76 | 297 | 446 |
| Procaviidae | *Procavia capensis* | Rock hyrax | Pcap_2.0_HiC | 8,632 (93.6%) | 8,567 | 65 | 167 | 427 |
| Orycteropodidae | *Orycteropus afer* | Aardvark | OryAfe1.0_HiC | 8,654 (93.8%) | 8,582 | 72 | 185 | 387 |
| Tenrecidae | *Echinops telfairi* | Lesser hedgehog tenrec | ASM31398v2 | 8,308 (90.0%) | 8,151 | 157 | 301 | 617 |
| Macroscelididae | *Elephantulus edwardii* | Cape elephant shrew | EleEdw1.0 | 8,727 (94.6%) | 8,035 | 692 | 139 | 360 |
| Chrysochloridae | *Chrysochloris asiatica* | Cape golden mole | ChrAsi1.0 | 8,853 (95.9%) | 8,591 | 262 | 132 | 241 |

### Table S4. BUSCO evaluation of afrotherian gene sets

For consistency, all genome assemblies were examined using the same version and library of BUSCO (5.0.0_cv1 with the 9,226-gene mammalia_odb10 dataset). Ddugon_BGI denotes the assembly generated in our study. Assemblies with the suffix _HiC were generated by DNA Zoo, the remainder are assemblies available via NCBI. Assembly Dugong_dugon_HiC is denoted 'Ddugon_DNAzoo' elsewhere in this manuscript.

| Afrotheria family | Latin name | Common name | Assembly | Complete BUSCOs (C) | Complete and single-copy BUSCOs (S) | Complete and duplicated BUSCOs (D) | Fragmented BUSCOs (F) | Missing BUSCOs (M) |
|---|---|---|---|---|---|---|---|---|
| Dugongidae | *Dugong dugon* | Dugong | Ddugon_BGI | 8,458 (91.7%) | 8,410 | 48 | 300 | 468 |
| Dugongidae | *Dugong dugon* | Dugong | Ddugon_HiC | 7,407 (80.3%) | 7,367 | 40 | 766 | 1,053 |
| Trichechidae | *Trichechus manatus* | West Indian manatee | TriManLat1.0_HiC | 7,358 (79.8%) | 7,316 | 42 | 784 | 1,084 |
| Elephantidae | *Elephas maximus* | Asian elephant | Elephas_maximus_HiC | 7,274 (78.9%) | 7,274 | 35 | 857 | 1,095 |
| Procaviidae | *Procavia capensis* | Rock hyrax | Pcap_2.0_HiC | 6,562 (71.2%) | 6,520 | 42 | 867 | 1,797 |
| Orycteropodidae | *Orycteropus afer* | Aardvark | OryAfe1.0_HiC | 6,892 (74.7%) | 6,860 | 32 | 962 | 1,372 |
| Tenrecidae | *Echinops telfairi* | Lesser hedgehog tenrec | ASM31398v2 | 8,608 (93.3%) | 5,901 | 2,707 | 276 | 342 |
| Macroscelididae | *Elephantulus edwardii* | Cape elephant shrew | EleEdw1.0 | 8,904 (96.5%) | 8,904 | 1,283 | 106 | 216 |
| Chrysochloridae | *Chrysochloris asiatica* | Cape golden mole | ChrAsi1.0 | 9,056 (98.2%) | 7,750 | 1,306 | 88 | 82 |

### Table S5 | Gene families expanded and contracted in sirenian genomes

Cdid, *Choloepus didactylus*; Oafe, *Orycteropus afer*; Eedw, *Elephantulus edwardii*; Pcap, *Procavia capensis*; Casi, *Chrysochloris asiatica*; Emax, *Elephas maximus*; Etel, *Echinops telfairi*; Ddug, *Dugong dugon*; Tman, *Trichechus manatus*. Nodes correspond to the NEXUS tree ((((Emax<0>,Pcap<2>)<1>,(Tman<4>,Ddug<6>)<5>)<3>,((Eedw<8>,(Etel<10>,Casi<12>)<11>)<9>,Oafe<14>)<13>)<7>,Cdid<16>)<15>.

| **Family ID** | **<11>** | **Cdid<16>** | **<3>** | **Oafe<14>** | **Eedw<8>** | **<1>** | **Pcap<2>** | **<7>** | **<13>** | **Casi<12>** | **Emax<0>** | **Etel<10>** | **<9>** | **<5>** | **Ddug<6>** | **Tman<4>** | **Gene family Description** |
|---|---|---|---|---|---|---|---|---|---|---|---|---|---|---|---|---|---|
| OG0000004 | 18 | 14 | 15 | 28 | 9 | 15 | 5 | 16 | 17 | 47 | 46 | 11 | 17 | 2 | 2 | 2 | olfactory receptor |
| OG0000006 | 16 | 28 | 17 | 1 | 15 | 17 | 13 | 17 | 16 | 19 | 35 | 17 | 16 | 13 | 5 | 16 | olfactory receptor 4F4-like |
| OG0000009 | 8 | 16 | 16 | 25 | 7 | 16 | 27 | 15 | 14 | 6 | 26 | 4 | 11 | 8 | 7 | 8 | vomeronasal type-1 receptor 2-like |
| OG0000018 | 12 | 25 | 9 | 23 | 9 | 9 | 3 | 12 | 12 | 15 | 9 | 9 | 12 | 5 | 3 | 5 | olfactory receptor |
| OG0000023 | 7 | 15 | 10 | 13 | 7 | 10 | 10 | 10 | 9 | 5 | 27 | 6 | 8 | 6 | 1 | 6 | olfactory receptor |
| OG0000024 | 10 | 24 | 9 | 7 | 6 | 9 | 7 | 9 | 9 | 10 | 9 | 17 | 9 | 6 | 7 | 1 | olfactory receptor |
| OG0000028 | 8 | 11 | 8 | 6 | 4 | 8 | 5 | 8 | 8 | 8 | 32 | 9 | 8 | 5 | 1 | 6 | olfactory receptor |
| OG0000029 | 10 | 17 | 8 | 14 | 2 | 8 | 0 | 9 | 9 | 23 | 15 | 8 | 9 | 2 | 1 | 2 | olfactory receptor |
| OG0000030 | 10 | 14 | 10 | 10 | 11 | 10 | 11 | 10 | 10 | 5 | 14 | 12 | 10 | 4 | 4 | 0 | olfactory receptor |
| OG0000034 | 10 | 12 | 6 | 21 | 9 | 6 | 3 | 9 | 10 | 18 | 7 | 8 | 10 | 1 | 0 | 1 | olfactory receptor |
| OG0000038 | 2 | 6 | 7 | 5 | 5 | 8 | 10 | 6 | 5 | 1 | 40 | 1 | 5 | 4 | 2 | 4 | olfactory receptor |
| OG0000042 | 2 | 12 | 10 | 3 | 2 | 10 | 22 | 9 | 4 | 1 | 20 | 2 | 3 | 5 | 4 | 5 | UDP-glucuronosyltransferase 2B31 |
| OG0000043 | 10 | 9 | 6 | 4 | 11 | 6 | 4 | 7 | 8 | 21 | 6 | 9 | 9 | 3 | 2 | 3 | keratin, type I |
| OG0000045 | 9 | 16 | 7 | 5 | 10 | 7 | 6 | 9 | 9 | 9 | 7 | 11 | 9 | 3 | 3 | 2 | olfactory receptor |
| OG0000046 | 3 | 8 | 9 | 7 | 10 | 9 | 18 | 8 | 7 | 1 | 14 | 1 | 7 | 6 | 4 | 6 | 60S ribosomal protein L9-like |
| OG0000051 | 7 | 15 | 7 | 6 | 8 | 7 | 2 | 7 | 7 | 9 | 13 | 5 | 7 | 4 | 4 | 3 | olfactory receptor |
| OG0000058 | 3 | 5 | 8 | 7 | 6 | 8 | 16 | 7 | 6 | 1 | 16 | 2 | 6 | 5 | 5 | 5 | HLA class II histocompatibility antigen |
| OG0000059 | 6 | 6 | 6 | 14 | 4 | 6 | 20 | 6 | 6 | 6 | 5 | 6 | 6 | 1 | 1 | 1 | saoe class I histocompatibility antigen |
| OG0000067 | 6 | 17 | 7 | 6 | 4 | 7 | 7 | 7 | 6 | 3 | 14 | 7 | 6 | 4 | 4 | 0 | olfactory receptor |
| OG0000071 | 7 | 12 | 7 | 6 | 5 | 7 | 12 | 7 | 7 | 4 | 9 | 9 | 7 | 1 | 1 | 1 | olfactory receptor |
| OG0000077 | 7 | 10 | 7 | 8 | 4 | 7 | 9 | 7 | 7 | 7 | 4 | 10 | 7 | 3 | 2 | 3 | glyceraldehyde-3-phosphate dehydrogenase-like |
| OG0000082 | 6 | 19 | 6 | 6 | 3 | 6 | 0 | 6 | 6 | 6 | 14 | 7 | 6 | 1 | 0 | 1 | olfactory receptor |

| | | | | | | | | | | | | | | | | |
|---|---|---|---|---|---|---|---|---|---|---|---|---|---|---|---|---|
| OG0000084 | 4 | 4 | 4 | 12 | 3 | 4 | 0 | 4 | 4 | 7 | 25 | 0 | 4 | 2 | 2 | 2 | melanoma-associated antigen B4-like |
| OG0000090 | 9 | 0 | 4 | 13 | 0 | 4 | 1 | 4 | 6 | 18 | 10 | 9 | 6 | 2 | 0 | 2 | olfactory receptor |
| OG0000091 | 6 | 14 | 6 | 2 | 6 | 6 | 7 | 6 | 6 | 5 | 3 | 9 | 6 | 3 | 3 | 3 | 60S ribosomal protein L17-like |
| OG0000094 | 4 | 7 | 5 | 16 | 7 | 5 | 5 | 5 | 5 | 3 | 6 | 3 | 5 | 2 | 2 | 2 | olfactory receptor |
| OG0000097 | 6 | 12 | 6 | 2 | 6 | 6 | 6 | 6 | 6 | 8 | 4 | 6 | 6 | 3 | 3 | 3 | olfactory receptor |
| OG0000120 | 6 | 5 | 5 | 5 | 9 | 5 | 2 | 5 | 6 | 6 | 8 | 9 | 6 | 1 | 1 | 1 | taste receptor type 2 member 3-like |
| OG0000123 | 4 | 0 | 3 | 0 | 3 | 3 | 2 | 3 | 3 | 7 | 29 | 5 | 3 | 0 | 0 | 0 | olfactory receptor |
| OG0000137 | 4 | 3 | 4 | 16 | 1 | 4 | 2 | 4 | 4 | 12 | 5 | 2 | 4 | 2 | 1 | 2 | olfactory receptor |
| OG0000144 | 5 | 4 | 5 | 2 | 7 | 5 | 3 | 5 | 5 | 4 | 11 | 10 | 5 | 1 | 1 | 1 | olfactory receptor |
| OG0000150 | 3 | 8 | 5 | 1 | 3 | 5 | 5 | 5 | 3 | 2 | 17 | 4 | 3 | 1 | 1 | 1 | olfactory receptor |
| OG0000157 | 4 | 9 | 4 | 11 | 2 | 4 | 1 | 4 | 4 | 5 | 7 | 1 | 4 | 2 | 2 | 2 | olfactory receptor |
| OG0000158 | 3 | 3 | 3 | 1 | 5 | 3 | 2 | 3 | 3 | 5 | 20 | 2 | 3 | 1 | 1 | 1 | olfactory receptor |
| OG0000166 | 4 | 9 | 4 | 1 | 3 | 4 | 1 | 4 | 4 | 4 | 9 | 10 | 4 | 1 | 1 | 1 | olfactory receptor |
| OG0000176 | 4 | 5 | 4 | 1 | 1 | 4 | 6 | 4 | 4 | 8 | 11 | 4 | 4 | 2 | 0 | 2 | olfactory receptor |
| OG0000197 | 4 | 9 | 4 | 7 | 4 | 4 | 1 | 4 | 4 | 4 | 6 | 3 | 4 | 2 | 0 | 2 | olfactory receptor |
| OG0000199 | 3 | 15 | 3 | 3 | 3 | 3 | 2 | 3 | 3 | 4 | 6 | 3 | 3 | 0 | 0 | 0 | olfactory receptor |
| OG0000207 | 4 | 3 | 4 | 3 | 5 | 4 | 7 | 4 | 4 | 3 | 5 | 6 | 4 | 2 | 1 | 2 | antigen-presenting glycoprotein CD1d |
| OG0000210 | 4 | 9 | 4 | 3 | 7 | 4 | 3 | 4 | 4 | 3 | 4 | 3 | 4 | 2 | 1 | 2 | olfactory receptor |
| OG0000211 | 5 | 6 | 4 | 2 | 4 | 4 | 2 | 4 | 4 | 6 | 5 | 6 | 4 | 2 | 2 | 2 | olfactory receptor |
| OG0000215 | 4 | 3 | 3 | 3 | 6 | 3 | 4 | 3 | 4 | 3 | 1 | 12 | 4 | 1 | 1 | 1 | 60S ribosomal protein L29-like |
| OG0000224 | 4 | 9 | 4 | 2 | 5 | 4 | 3 | 4 | 4 | 1 | 5 | 5 | 4 | 2 | 2 | 2 | 60S ribosomal protein L29-like |
| OG0000243 | 4 | 5 | 4 | 5 | 5 | 4 | 2 | 4 | 4 | 1 | 4 | 7 | 4 | 2 | 2 | 2 | 60S ribosomal protein L29-like |
| OG0000266 | 2 | 8 | 4 | 1 | 6 | 4 | 4 | 4 | 3 | 1 | 7 | 1 | 3 | 2 | 2 | 2 | putative LRRC37B-like protein 1-like |
| OG0000274 | 4 | 9 | 4 | 4 | 1 | 4 | 3 | 4 | 4 | 2 | 5 | 5 | 4 | 2 | 0 | 2 | glycine cleavage system H protein, mitochondrial-like |
| OG0000308 | 3 | 3 | 3 | 4 | 9 | 3 | 5 | 3 | 3 | 1 | 3 | 3 | 3 | 1 | 0 | 1 | cytochrome P450 2J2-like |
| OG0000345 | 1 | 2 | 4 | 1 | 4 | 4 | 7 | 3 | 2 | 1 | 10 | 0 | 2 | 2 | 1 | 2 | olfactory receptor |
| OG0000347 | 7 | 1 | 2 | 0 | 4 | 2 | 0 | 2 | 2 | 12 | 3 | 8 | 4 | 0 | 0 | 0 | olfactory receptor |

| | | | | | | | | | | | | | | | | | |
|---|---|---|---|---|---|---|---|---|---|---|---|---|---|---|---|---|---|
| OG0000348 | 2 | 8 | 3 | 2 | 3 | 3 | 2 | 3 | 3 | 1 | 8 | 2 | 3 | 1 | 1 | 1 | olfactory receptor |
| OG0000437 | 2 | 5 | 3 | 2 | 1 | 3 | 3 | 3 | 2 | 1 | 9 | 3 | 2 | 1 | 1 | 1 | olfactory receptor |
| OG0000506 | 2 | 3 | 3 | 3 | 1 | 3 | 6 | 3 | 3 | 2 | 6 | 1 | 2 | 1 | 1 | 1 | olfactory receptor |
| OG0000523 | 3 | 2 | 3 | 4 | 3 | 3 | 2 | 3 | 3 | 5 | 6 | 2 | 3 | 0 | 0 | 0 | olfactory receptor |
| OG0000554 | 3 | 7 | 2 | 2 | 5 | 2 | 2 | 3 | 3 | 4 | 2 | 1 | 3 | 0 | 0 | 0 | late cornified envelope protein 3E （LCE gene family） |
| OG0000576 | 2 | 9 | 3 | 3 | 1 | 3 | 3 | 3 | 3 | 2 | 3 | 1 | 2 | 1 | 1 | 0 | olfactory receptor |
| OG0000610 | 2 | 4 | 3 | 0 | 3 | 3 | 3 | 3 | 2 | 2 | 8 | 2 | 2 | 0 | 0 | 0 | olfactory receptor |
| OG0000883 | 2 | 3 | 2 | 0 | 4 | 2 | 2 | 2 | 2 | 2 | 7 | 1 | 2 | 0 | 0 | 0 | olfactory receptor |
| OG0000904 | 0 | 1 | 3 | 1 | 0 | 3 | 9 | 2 | 1 | 0 | 7 | 0 | 0 | 1 | 0 | 1 | LOC119515707 |
| OG0001087 | 1 | 4 | 3 | 1 | 2 | 3 | 4 | 3 | 2 | 0 | 5 | 1 | 2 | 1 | 0 | 1 | olfactory receptor |
| OG0001240 | 1 | 5 | 2 | 1 | 2 | 2 | 2 | 2 | 2 | 1 | 5 | 1 | 2 | 0 | 0 | 0 | olfactory receptor |
| OG0001547 | 1 | 4 | 2 | 2 | 4 | 2 | 3 | 2 | 2 | 0 | 1 | 1 | 2 | 0 | 0 | 0 | C2orf78-like |
| OG0005176 | 0 | 2 | 2 | 2 | 0 | 2 | 3 | 2 | 2 | 0 | 3 | 0 | 0 | 0 | 0 | 0 | vomeronasal type-1 receptor 90-like |
| OG0000001 | 23 | 2 | 28 | 7 | 3 | 28 | 26 | 20 | 19 | 21 | 39 | 50 | 19 | 52 | 48 | 75 | zinc finger protein 420-like |
| OG0000032 | 3 | 5 | 8 | 16 | 4 | 8 | 10 | 7 | 7 | 1 | 6 | 3 | 5 | 15 | 14 | 21 | ferritin, mitochondrial |
| OG0000049 | 4 | 25 | 6 | 3 | 5 | 6 | 3 | 6 | 5 | 1 | 5 | 4 | 5 | 10 | 10 | 11 | elongation factor 1-alpha 2 |
| OG0000060 | 6 | 15 | 6 | 1 | 8 | 6 | 5 | 6 | 6 | 8 | 2 | 3 | 6 | 9 | 12 | 9 | histone H3.2-like |
| OG0000063 | 4 | 1 | 6 | 15 | 1 | 6 | 4 | 5 | 5 | 4 | 8 | 2 | 4 | 11 | 10 | 17 | elongation factor 1-alpha 1, partial |
| OG0000110 | 1 | 0 | 3 | 0 | 1 | 3 | 1 | 1 | 1 | 3 | 13 | 0 | 1 | 7 | 5 | 25 | putative serine protease K12H4.7-like |
| OG0000290 | 1 | 0 | 3 | 4 | 0 | 3 | 1 | 2 | 2 | 0 | 4 | 1 | 1 | 8 | 8 | 13 | |
| OG0000296 | 2 | 1 | 3 | 2 | 1 | 3 | 4 | 2 | 2 | 3 | 2 | 1 | 2 | 7 | 7 | 9 | leukemia inhibitory factor-like |
| OG0000457 | 2 | 1 | 2 | 4 | 1 | 2 | 2 | 2 | 2 | 1 | 2 | 3 | 2 | 4 | 4 | 7 | UPF0711 protein C18orf21 homolog |
| OG0000520 | 2 | 1 | 2 | 2 | 2 | 2 | 2 | 2 | 2 | 1 | 3 | 4 | 2 | 4 | 5 | 4 | protein FAM47E |
| OG0000620 | 1 | 2 | 2 | 1 | 2 | 2 | 2 | 2 | 2 | 1 | 1 | 1 | 2 | 4 | 3 | 9 | histo-blood group ABO system transferase-like |
| OG0000765 | 1 | 0 | 2 | 4 | 1 | 2 | 2 | 2 | 2 | 1 | 3 | 1 | 1 | 4 | 4 | 4 | probable palmitoyltransferase ZDHHC11-like |
| OG0001367 | 1 | 1 | 1 | 1 | 2 | 1 | 1 | 1 | 1 | 1 | 1 | 1 | 1 | 3 | 3 | 5 | MKI67 FHA domain-interacting nucleolar phosphoprotein |

| | | | | | | | | | | | | | | | |
|---|---|---|---|---|---|---|---|---|---|---|---|---|---|---|---|
| OG0001770 | 0 | 0 | 1 | 4 | 0 | 1 | 1 | 1 | 1 | 0 | 2 | 0 | 0 | 3 | 4 | 3 |

**Table S6 | Gene function enrichment of expanded or contracted sirenian gene families**

GO denotes Gene ontology (biological process); KEGG, KEGG Kyoto Encyclopedia of Genes and Genomes Pathways.

| Terms | Database | ID | *P*-value | Adjusted *P*-value |
|---|---|---|---|---|
| Detection of chemical stimulus involved in sensory Perception of smell | Gene Ontology | GO:0050911 | $6.80\times10^{-15}$ | $7.66\times10^{-13}$ |
| Olfactory receptor activity | Gene Ontology | GO:0004984 | $6.80\times10^{-15}$ | $7.66\times10^{-13}$ |
| G protein-coupled receptor activity | Gene Ontology | GO:0004930 | $1.46\times10^{-13}$ | $8.21\times10^{-12}$ |
| Odorant binding | Gene Ontology | GO:0005549 | $9.38\times10^{-12}$ | $3.52\times10^{-10}$ |
| G protein-coupled receptor signaling pathway | Gene Ontology | GO:0007186 | $2.82\times10^{-11}$ | $9.07\times10^{-10}$ |
| Cytosolic large ribosomal subunit | Gene Ontology | GO:0022625 | $3.68\times10^{-9}$ | $1.04\times10^{-7}$ |
| SRP-dependent cotranslational protein targeting to membrane | Gene Ontology | GO:0006614 | $3.15\times10^{-8}$ | $7.87\times10^{-7}$ |
| Viral transcription | Gene Ontology | GO:0019083 | $8.38\times10^{-8}$ | $1.89\times10^{-6}$ |
| Nuclear-transcribed mRNA catabolic process, Nonsense-mediated decay | Gene Ontology | GO:0000184 | $1.18\times10^{-7}$ | $2.21\times10^{-6}$ |
| Translation elongation factor activity | Gene Ontology | GO:0003746 | $1.67\times10^{-6}$ | $2.21\times10^{-5}$ |
| Cornification | Gene Ontology | GO:0070268 | $4.76\times10^{-6}$ | $5.36\times10^{-5}$ |
| Antigen processing and presentation, endogenous Lipid antigen via MHC class Ib | Gene Ontology | GO:0048006 | $1.91\times10^{-5}$ | 0.00015897 |
| Endogenous lipid antigen binding | Gene Ontology | GO:0030883 | $1.91\times10^{-5}$ | 0.00015897 |
| Exogenous lipid antigen binding | Gene Ontology | GO:0030884 | $1.91\times10^{-5}$ | 0.00015897 |
| Keratinization | Gene Ontology | GO:0031424 | $2.73\times10^{-5}$ | 0.00020448 |
| Ferroxidase activity | Gene Ontology | GO:0004322 | $4.98\times10^{-5}$ | 0.00033982 |
| Sensory perception of smell | Gene Ontology | GO:0007608 | 0.00017698 | 0.00107621 |
| Positive regulation of T cell mediated cytotoxicity | Gene Ontology | GO:0001916 | 0.00029183 | 0.00172793 |
| Immune response | Gene Ontology | GO:0006955 | 0.00078533 | 0.00392667 |
| Antimicrobial humoral immune response mediated by antimicrobial peptide | Gene Ontology | GO:0061844 | 0.00176877 | 0.00829112 |
| Olfactory transduction | KEGG | hsa04740 | $1.19\times10^{-14}$ | $8.89\times10^{-13}$ |

| | | | | |
|---|---|---|---|---|
| Ribosome | KEGG | hsa03010 | $4.45\times10^{-7}$ | $6.68\times10^{-6}$ |
| Estrogen signaling pathway | KEGG | hsa04915 | $1.10\times10^{-5}$ | 0.00011229 |
| Legionellosis | KEGG | hsa05134 | $2.48\times10^{-5}$ | 0.00019722 |
| Ferroptosis | KEGG | hsa04216 | 0.00076561 | 0.00391504 |
| Hematopoietic cell lineage | KEGG | hsa04640 | 0.00416644 | 0.01802785 |
| Carbon metabolism | KEGG | hsa01200 | 0.0059574 | 0.02393598 |
| RNA transport | KEGG | hsa03013 | 0.01142395 | 0.03418939 |
| Tight junction | KEGG | hsa04530 | 0.01208376 | 0.03530968 |

**Table S7 | Positively selected genes at the ancestral branch of Sirenia**

Positively selected genes (PSG) were identified using PAML with branch-site models. *P*-value denotes a $\chi 2$-test statistic and adjusted *P*-value its multiple testing correction using the Benjamini-Hochberg method. A *P*-value or adjusted *P*-value of 0 indicates a number below $2.22\times10^{-16}$.

| Genes | Name | *P*-value | Adjusted *P*-value |
|---|---|---|---|
| *ACNAT2* | acyl-coenzyme A amino acid N-acyltransferase 2 | 0.00053156 | 0.05095063 |
| *ANK1* | ankyrin 1 | $2.40\times10^{-6}$ | 0.00088731 |
| *ANPEP* | alanyl aminopeptidase, membrane | $4.34\times10^{-5}$ | 0.01247991 |
| *ATP1B4* | ATPase Na+/K+ transporting family member beta 4 | $1.10\times10^{-6}$ | 0.00047447 |
| *ATP6V0A2* | ATPase H+ transporting V0 subunit a2 | $3.18\times10^{-9}$ | $2.06\times10^{-6}$ |
| *BAP18* | Chromatin complexes subunit bap18 | $2.40\times10^{-8}$ | $1.24\times10^{-5}$ |
| *CCDC39* | coiled-coil domain containing 39 | 0.00071635 | 0.05793448 |
| *CD34* | CD34 molecule | $9.54\times10^{-5}$ | 0.02139378 |
| *CENPB* | centromere protein B | 0.00014053 | 0.02139378 |
| *CPA1* | carboxypeptidase A1 | 0.00013905 | 0.02139378 |
| *DGLUCY* | D-glutamate cyclase | 0 | 0 |
| *DSG2* | desmoglein 2 | 0.00017564 | 0.02525299 |
| *DUOXA2* | dual oxidase maturation factor 2 | 0.00066794 | 0.0576207 |
| *EBP* | EBP cholestenol delta-isomerase | 0.0002267 | 0.02793832 |
| *ENPP6* | ectonucleotide pyrophosphatase/phosphodiesterase 6 | $9.17\times10^{-5}$ | 0.02139378 |
| *ERI1* | exoribonuclease 1 | 0.00020555 | 0.02659843 |
| *IDO1* | indoleamine 2,3-dioxygenase 1 | 0.00062997 | 0.05621939 |
| *MAG* | myelin associated glycoprotein | 0.00031966 | 0.03596836 |
| *METTL2A* | methyltransferase 2A, methylcytidine | 0.00052871 | 0.05095063 |
| *MMP27* | matrix metallopeptidase 27 | 0.0001252 | 0.02139378 |
| *MOG* | myelin oligodendrocyte glycoprotein | 0.000133 | 0.02139378 |
| *MPO* | myeloperoxidase | 0.00057974 | 0.05358454 |
| *NCBP2* | nuclear cap binding protein subunit 2 | 0.00050518 | 0.05095063 |
| *PPP4R3B* | protein phosphatase 4 regulatory subunit 3B | 0.00052968 | 0.05095063 |

| | | | |
|---|---|---|---|
| *PTPRJ* | protein tyrosine phosphatase receptor type J | $8.99\times10^{-15}$ | $7.76\times10^{-12}$ |
| *SETDB2* | SET domain bifurcated histone lysine methyltransferase 2 | 0.00019891 | 0.02659843 |
| *SPTB* | spectrin beta, erythrocytic | 0.00010855 | 0.02139378 |
| *STAB1* | stabilin 1 | $2.10\times10^{-5}$ | 0.0067935 |
| *VTN* | vitronectin | 0.00030595 | 0.03596836 |
| *ZFHX3* | zinc finger homeobox 3 | 0.00011686 | 0.02139378 |

**Table S8 | Rapidly evolving genes at the ancestral branch of Sirenia**

Rapidly evolving genes (REGs) were identified using PAML with branch-site models. *P*-value denotes a χ2-test statistic and adjusted *P*-value its multiple testing correction using the Benjamini-Hochberg method. A *P*-value or adjusted *P*-value of 0 indicates a number below $2.22{\times}10^{-16}$.

| Genes | Name | *P*-value | Adjusted *P*-value |
|---|---|---|---|
| *ABCB1* | ATP binding cassette subfamily B member 1 | $1.24{\times}10^{-5}$ | 0.00409465 |
| *ABCB11* | ATP binding cassette subfamily B member 11 | 0.00017477 | 0.02517397 |
| *ABCC2* | ATP binding cassette subfamily C member 2 | 0.00028074 | 0.03141092 |
| *ABCF3* | ATP binding cassette subfamily F member 3 | $3.03{\times}10^{-5}$ | 0.00738499 |
| *ABRA* | actin binding Rho activating protein | 0.00026921 | 0.03092321 |
| *ACACA* | acetyl-CoA carboxylase alpha | 0.00019939 | 0.02535466 |
| *ACADM* | acyl-CoA dehydrogenase medium chain | 0.00031883 | 0.03381749 |
| *ACSS3* | acyl-CoA synthetase short chain family member 3 | $2.09{\times}10^{-12}$ | $6.11{\times}10^{-9}$ |
| *ALB* | albumin | $6.61{\times}10^{-5}$ | 0.01258586 |
| *ANK1* | ankyrin 1 | 0 | 0 |
| *ANKRD6* | ankyrin repeat domain 6 | $6.67{\times}10^{-5}$ | 0.01258586 |
| *ANXA2* | annexin A2 | $4.35{\times}10^{-7}$ | 0.00031807 |
| *APH1A* | aph-1 homolog A, gamma-secretase subunit | 0.00059065 | 0.04934763 |
| *ATP11C* | ATPase phospholipid transporting 11C | $6.21{\times}10^{-9}$ | $8.07{\times}10^{-6}$ |
| *BAP18* | BCL2 related protein A1 | 0.00019619 | 0.02535466 |
| *BCL2A1* | C1q and TNF related 2 | 0.00019025 | 0.02535466 |
| *C1QTNF2* | C1q and TNF related 2 | 0.00019611 | 0.02535466 |
| *C3ORF49* | chromosome 3 open reading frame 49 | 0.00017234 | 0.02517397 |
| *CAP2* | cyclase associated actin cytoskeleton regulatory protein 2 | 0.00026346 | 0.03082265 |
| *CAPN5* | calpain 5 | $2.41{\times}10^{-5}$ | 0.00655688 |
| *CARMIL3* | capping protein regulator and myosin 1 linker 3 | $3.94{\times}10^{-13}$ | $1.54{\times}10^{-9}$ |
| *CCRL2* | C-C motif chemokine receptor like 2 | 0.00010053 | 0.01704409 |
| *CD34* | CD34 molecule | 0.00037189 | 0.03687089 |
| *ADGRE5* | Adhesion G protein-coupled receptor E5 | 0.00037017 | 0.03687089 |

| | | | |
|---|---|---|---|
| *CLCN1* | chloride voltage-gated channel 1 | 0.00046366 | 0.04269724 |
| *CLTCL1* | clathrin heavy chain like 1 | $2.02\times10^{-5}$ | 0.0057639 |
| *CMTM7* | CKLF like MARVEL transmembrane domain containing 7 | $2.55\times10^{-5}$ | 0.0067801 |
| *COL1A2* | collagen type I alpha 2 chain | 0.00020373 | 0.02562874 |
| *COL5A2* | collagen type V alpha 2 chain | 0.0004323 | 0.04118907 |
| *COL9A2* | collagen type IX alpha 2 chain | 0.00059014 | 0.04934763 |
| *CORO6* | coronin 6 | $2.74\times10^{-5}$ | 0.00712339 |
| *CPA4* | carboxypeptidase A4 | 0.00045017 | 0.04179811 |
| *CPS1* | carbamoyl-phosphate synthase 1 | $1.71\times10^{-7}$ | 0.00014289 |
| *CYP2F3* | cytochrome P450 family 2 subfamily F member 3 | $3.27\times10^{-5}$ | 0.00765115 |
| *DAP* | death associated protein | $9.40\times10^{-5}$ | 0.01641352 |
| *DHX36* | DEAH-box helicase 36 | $3.68\times10^{-5}$ | 0.00812308 |
| *DOCK11* | dedicator of cytokinesis 11 | 0.00060018 | 0.04944739 |
| *DOCK2* | dedicator of cytokinesis 2 | $2.04\times10^{-6}$ | 0.00099442 |
| *DSC1* | desmocollin 1 | 0.00029289 | 0.03208281 |
| *DSE* | dermatan sulfate epimerase | $6.39\times10^{-5}$ | 0.01245944 |
| *DSG2* | desmoglein 2 | $1.16\times10^{-}5$ | 0.00399142 |
| *DSP* | desmoplakin | $4.81\times10^{-}10$ | $8.04\times10^{-7}$ |
| *ELN* | elastin | $4.72\times10^{-5}$ | 0.01003987 |
| *ENPEP* | glutamyl aminopeptidase | 0.00010939 | 0.01729334 |
| *EPB42* | erythrocyte membrane protein band 4.2 | $7.10\times10^{-6}$ | 0.00259572 |
| *FAM219B* | family with sequence similarity 219 member B | 0.00012215 | 0.01880229 |
| *FAT4* | FAT atypical cadherin 4 | 0.00053451 | 0.04605698 |
| *GLG1* | golgi glycoprotein 1 | 0.00046716 | 0.04269724 |
| *GPR82* | G protein-coupled receptor 82 | $3.57\times10^{-5}$ | 0.00812308 |
| *GRIN2A* | glutamate ionotropic receptor NMDA type subunit 2A | $6.96\times10^{-5}$ | 0.01292461 |
| *HK3* | hexokinase 3 | 0.00026196 | 0.03082265 |
| *HSPA8* | heat shock protein family A (Hsp70) member 8 | $8.15\times10^{-6}$ | 0.0028893 |

| | | | |
|---|---|---|---|
| *ITGA1* | integrin subunit alpha 1 | 0.00035134 | 0.03574248 |
| *ITGA8* | integrin subunit alpha 8 | 0.00044085 | 0.04125994 |
| *JPH4* | junctophilin 4 | $8.36\times10^{-5}$ | 0.01504671 |
| *KCNE2* | potassium voltage-gated channel subfamily E regulatory subunit 2 | 0.00032086 | 0.03381749 |
| *KCNH1* | potassium voltage-gated channel subfamily H member 1 | 0.00029343 | 0.03208281 |
| *KCNJ10* | potassium inwardly rectifying channel subfamily J member 10 | 0.00025714 | 0.03069662 |
| *KEL* | Kell metallo-endopeptidase (Kell blood group) | $1.38\times10^{-5}$ | 0.00436341 |
| *KIAA1109* | kinesin family member 21A | 0.00011556 | 0.01802644 |
| *KIF21A* | keratinocyte proline rich protein | $2.94\times10^{-6}$ | 0.00127389 |
| *KPRP* | lipase A, lysosomal acid type | $7.52\times10^{-7}$ | 0.00046303 |
| *LIPA* | lipoprotein lipase | $1.26\times10^{-5}$ | 0.00409465 |
| *LPL* | LY6/PLAUR domain containing 8 | 0.00010935 | 0.01729334 |
| *LYPD8* | LY6/PLAUR domain containing 8 | $1.92\times10^{-5}$ | 0.00575951 |
| *MAG* | myelin associated glycoprotein | $1.29\times10^{-6}$ | 0.00068599 |
| *MMP19* | matrix metallopeptidase 19 | 0.00023962 | 0.02920095 |
| *MON2* | MON2 homolog, regulator of endosome-to-Golgi trafficking | $4.92\times10^{-5}$ | 0.01009808 |
| *MYBPC1* | myosin binding protein C1 | $2.45\times10^{-7}$ | 0.00019108 |
| *MYH1* | myosin heavy chain 1 | $3.56\times10^{-6}$ | 0.00148744 |
| *MYH2* | myosin heavy chain 2 | 0.00032798 | 0.0340249 |
| *MYO5C* | myosin VC | 0.00016642 | 0.02464461 |
| *MYO6* | myosin VI | $4.82\times10^{-5}$ | 0.0100695 |
| *MYOF* | myoferlin | $5.40\times10^{-5}$ | 0.01089217 |
| *NBEA* | neurobeachin | $5.56\times10^{-6}$ | 0.00216821 |
| *NECTIN4* | nectin cell adhesion molecule 4 | $2.01\times10^{-8}$ | $2.14\times10^{-5}$ |
| *NFATC4* | nuclear factor of activated T cells 4 | 0.00036693 | 0.03687089 |
| *NLRX1* | NLR family member X1 | 0.00043657 | 0.04118907 |
| *NT5C2* | 5'-nucleotidase, cytosolic II | 0.00018796 | 0.02535466 |
| *PAQR9* | progestin and adipoQ receptor family member 9 | $1.23\times10^{-6}$ | 0.00068523 |

| | | | |
|---|---|---|---|
| *PCDHGA2* | protocadherin gamma subfamily A, 2 | 0.00018451 | 0.02535466 |
| *PCOLCE2* | procollagen C-endopeptidase enhancer 2 | $8.87\times10^{-9}$ | $1.04\times10^{-5}$ |
| *PGK2* | phosphoglycerate kinase 2 | $2.35\times10^{-8}$ | $2.29\times10^{-5}$ |
| *PKP3* | plakophilin 3 | 0.00010937 | 0.01729334 |
| *PODXL* | podocalyxin like | 0.00029849 | 0.03233409 |
| *POLB* | DNA polymerase beta | 0.00048484 | 0.04366778 |
| *PPP4R3B* | protein phosphatase 4 regulatory subunit 3B | $5.50\times10^{-6}$ | 0.00216821 |
| *PTGIR* | prostaglandin I2 receptor | 0. 00030919 | 0.03318513 |
| *PTPRJ* | protein tyrosine phosphatase receptor type J | $5.04\times10^{-7}$ | 0.00034684 |
| *PXYLP1* | 2-phosphoxylose phosphatase 1 | 0.00010423 | 0.01729334 |
| *PYGM* | glycogen phosphorylase, muscle associated | 0.00059475 | 0.04934763 |
| *RAG1* | recombination activating 1 | $1.03\times10^{-6}$ | 0.0006025 |
| *RARA* | retinoic acid receptor alpha | $2.38\times10^{-9}$ | $3.48\times10^{-6}$ |
| *RNASE10* | ribonuclease A family member 10 (inactive) | 0.00019066 | 0.02535466 |
| *RTN1* | reticulon 1 | 0.00053189 | 0.04605698 |
| *RYR3* | ryanodine receptor 3 | 0.00017645 | 0.02517397 |
| *SCD* | stearoyl-CoA desaturase | 0.00028192 | 0.03141092 |
| *SCN5A* | sodium voltage-gated channel alpha subunit 5 | $2.40\times10^{-6}$ | 0.00107991 |
| *SEMA4A* | semaphorin 4A | $2.99\times10^{-5}$ | 0.00738499 |
| *SERPINE2* | serpin family E member 2 | $5.51\times10^{-5}$ | 0.01092568 |
| *SH3RF1* | SH3 domain containing ring finger 1 | 0.00053211 | 0.04605698 |
| *SHISA4* | shisa family member 4 | 0.0001559 | 0.0233836 |
| *SLC13A1* | solute carrier family 13 member 1 | 0.00026961 | 0.03092321 |
| *SLC22A7* | solute carrier family 22 member 7 | $2.93\times10^{-5}$ | 0.00738499 |
| *SLC26A2* | solute carrier family 26 member 2 | 0.00022539 | 0.02775606 |
| *SLC31A1* | solute carrier family 31 member 1 | 0.00033155 | 0.0340249 |
| *SLC4A1* | solute carrier family 4 member 1 (Diego blood group) | $1.98\times10^{-5}$ | 0.0057639 |

| | | | |
|---|---|---|---|
| *SLC4A9* | solute carrier family 4 member 9 | 0.00043517 | 0.04118907 |
| *SLC5A1* | solute carrier family 5 member 1 | 0.00019849 | 0.02535466 |
| *SLC6A3* | solute carrier family 6 member 3 | 0 | 0 |
| *SPNS1* | sphingolipid transporter 1 (putative) | 0.00021476 | 0.02672885 |
| *SPTB* | spectrin beta, erythrocytic | $4.25\times10^{-11}$ | $8.29\times10^{-8}$ |
| *SRR* | serine racemase | 0.00027804 | 0.03141092 |
| *SYPL1* | synaptophysin like 1 | 0.00053541 | 0.04605698 |
| *TAOK3* | TAO kinase 3 | 0.00025582 | 0.03069662 |
| *TFAP4* | transcription factor AP-4 | $1.42\times10^{-5}$ | 0.00437173 |
| *TGFBR3* | transforming growth factor beta receptor 3 | $2.38\times10^{-5}$ | 0.00655688 |
| *TGM1* | transglutaminase 1 | 0.00042648 | 0.04118907 |
| *TLR7* | toll like receptor 7 | 0.00049014 | 0.04377212 |
| *TNNC2* | troponin C2, fast skeletal type | $9.65\times10^{-5}$ | 0.01660226 |
| *TNS3* | tensin 3 | 0.00033066 | 0.0340249 |
| *TRANK1* | tetratricopeptide repeat and ankyrin repeat containing 1 | $9.90\times10^{-8}$ | $8.91\times10^{-5}$ |
| *TRPM6* | transient receptor potential cation channel subfamily M member 6 | $1.66\times10^{-6}$ | 0.00084436 |
| *TSPAN33* | tetraspanin 33 | 0.0003762 | 0.03698486 |
| *TSPAN9* | tetraspanin 9 | 0.00019881 | 0.02535466 |
| *TUBB2B* | tubulin beta 2B class IIb | $5.61\times10^{-12}$ | $1.31\times10^{-8}$ |
| *UNC13C* | unc-13 homolog C | 0.00018075 | 0.02535466 |
| *VPS13C* | vacuolar protein sorting 13 homolog C | $2.33\times10^{-6}$ | 0.00107991 |
| *XPNPEP2* | X-prolyl aminopeptidase 2 | $9.01\times10^{-5}$ | 0.01597091 |
| *YIPF4* | Yip1 domain family member 4 | $3.68\times10^{-5}$ | 0.00812308 |
| *ZFHX2* | zinc finger homeobox 2 | $6.90\times10^{-6}$ | 0.00259572 |
| *ZYX* | zyxin | 0.00048524 | 0.04366778 |

### Table S9 | Genes with inactivating mutations in sirenians

Please note that stop codon of *KCNK18* residue 462 in West Indian manatee and Steller's sea cow, but not dugong, presumably arose independently because the site is preceded by internal deletions in Dugongidae (Steller's sea cow and dugong) but not in the West Indian manatee.

| Gene | *Name* | *Species* | Database | Sirenian residue (AA) | Human residue (AA) | Type | NCBI SRA ID or dugong RNA-seq read |
|---|---|---|---|---|---|---|---|
| *CHIA5* | Chitinase 5 | *Trichechus manatus* | NCBI | 49 | 49 | 2bp deletion | gnl\|SRA\|SRR8616893.222332505.1:34-144 222332505 |
| | | | | | | | gnl\|SRA\|SRR8616893.29600636.1:27-137 29600636 |
| | | | | | | | gnl\|SRA\|SRR8616893.14054244.1:27-137 14054244 |
| | | | | 116 | 116 | Premature stop codon | SRA:SRR331138.54636371.1 |
| | | | | | | | SRA:SRR331138.18846010.3 |
| | | | | | | | SRA:SRR331138.48555459.3 |
| | | | | 139 | 139 | Premature stop codon | gnl\|SRA\|SRR8616893.210438724.2:46-142 210438724 |
| | | | | | | | gnl\|SRA\|SRR8616893.176672544.2:16-112 176672544 |
| | | | | | | | gnl\|SRA\|SRR8616893.159794561.1:2-98 159794561 |
| | | | | 192 | 192 | 1bp deletion | gnl\|SRA\|SRR8616893.129252417.1:60-135 129252417 |
| | | | | | | | gnl\|SRA\|SRR8616893.124944060.1:37-111 124944060 |
| | | | | | | | gnl\|SRA\|SRR8616893.49347084.2:28-102 49347084 |

| | | | | | | | |
|---|---|---|---|---|---|---|---|
| | | | | 333 | 333 | Premature stop codon | gnl\|SRA\|SRR8616893.157292849.2:30-122 157292849 |
| | | | | | | | gnl\|SRA\|SRR8616893.146353683.1:30-122 146353683 |
| | | | | | | | gnl\|SRA\|SRR8616893.128710503.1:19-111 128710503 |
| | | *Dugong dugon* | In this study | 49 | 49 | 2bp deletion | lcl\|V350022740L4C004R0631339513*909_617_240/2_19037110___1 |
| | | | | | | | lcl\|V350022740L4C002R0031242885*221_747_383/2_1419688___1 |
| | | | | | | | V350022740L4C003R0070319403*603_482_1284/2_24154153___1 |
| | | | | 116 | 116 | Premature stop codon | V350022740L4C004R0450737534#0_0_0/1 |
| | | | | | | | V350022740L4C006R0241189258#813_225_692/2 |
| | | | | | | | V350022740L4C004R0310773487#952_771_1157/2 |
| | | | | 139 | 139 | Premature stop codon | lcl\|V350022740L4C003R0660197361*323_1430_241/2_10679638___1 |
| | | | | | | | lcl\|V350022740L4C001R0390251877*779_825_275/2_6486318___1 |
| | | | | | | | lcl\|V350022740L4C003R0660197361*323_1430_241/1_10679638___1 |
| | | | | 333 | 333 | Premature stop codon | lcl\|V350022740L4C003R0640450876*0_0_0/2_0___1 |
| | | | | | | | lcl\|V350022740L4C003R0310199110*574_276_1372/2_25194423___1 |
| | | | | | | | lcl\|V350022740L4C005R0421125977*1241_1402_951/1_9032551___1 |
| *KLK8* | Kallikrein 8 | *Trichechus manatus* | NCBI | 89 | 89 | Premature stop codon | gnl\|SRA\|SRR8616893.194922441.1:50-134 194922441 |

| | | | | | |
|---|---|---|---|---|---|
| | | | | | gnl\|SRA\|SRR8616893.224186147.1:32-116 224186147 |
| | | | | | gnl\|SRA\|SRR8616893.147126882.2:4-88 147126882 |
| | | 156 | 156 | Premature stop codon | gnl\|SRA\|SRR8616893.250007951.1:54-109 250007951 |
| | | | | | gnl\|SRA\|SRR8616893.251832290.1:54-109 251832290 |
| | | | | | gnl\|SRA\|SRR8616893.245022569.1:36-91 245022569 |
| | | 258 | 258 | 2bp deletion | gnl\|SRA\|SRR8616893.240002590.2:72-108 240002590 |
| | | | | | gnl\|SRA\|SRR8616893.250852032.2:35-71 250852032 |
| | | | | | gnl\|SRA\|SRR8616893.237707383.1:70-106 237707383 |
| *Dugong dugon* | In this study | 10 | 10 | Premature stop codon | lcl\|V350022740L4C004R0380568886*134_199_1205/2_20013072___1 |
| | | | | | lcl\|V350022740L4C004R0380568885*223_1205_1143/2_27300194___1 |
| | | | | | lcl\|V350022740L4C003R0440186991*1251_150_1285/2_1473243___1 |
| | | 109 | 109 | 2bp insertion | lcl\|V350022740L4C001R0340380110*1431_108_1239/1_2638775___1 |
| | | | | | lcl\|V350022740L4C003R0430499513*71_1315_276/1_15137165___1 |
| | | | | | lcl\|V350022740L4C005R0530258529*1071_134_1450/2_6269869___1 |
| | | 157 | 157 | Premature stop codon | lcl\|V350022740L4C004R0630404385*937_96_1014/2_16071847___1 |

| | | | | | | | |
|---|---|---|---|---|---|---|---|
| | | | | | | | lcl\|V350022740L4C006R0350606850*59_1009_1081/2_7952156___1 |
| | | | | | | | lcl\|V350022740L4C006R0130902588*1096_825_1361/1_21795415___1 |
| *LYG1* | Lysozyme g1 | *Trichechus manatus* | NCBI | 156 | 148 | Premature stop codon | SRR8616893.101515585.1_101515585 |
| | | | | | | | SRR8616893.93027165.1_93027165 |
| | | | | | | | SRR8616893.89507411.2_89507411 |
| | | *Dugong dugon* | In this study | 149 | 141 | 1bp deletion | lcl\|V350022740L4C002R0720427604*732_455_1239/2_14127157___1 |
| | | | | | | | lcl\|V350022740L4C003R0390516922*732_455_1239/2_14127157___1 |
| | | | | | | | lcl\|V350022740L4C005R0230872953*778_211_96/2_9624662___1 |
| | | | | 156 | 148 | Premature stop codon | V350022740L4C005R0410405174#185_1184_175/2 |
| | | | | | | | V350022740L4C005R0230872953#778_211_96/2 |
| | | | | | | | V350022740L4C003R0390516922#732_455_1239/2 |
| *ALOX15* | Arachidonate 15-lipoxygenase | *Trichechus manatus* | NCBI | 247 | 245 | 2bp deletion | gnl\|SRA\|SRR24090881.2231088.1:3566-3665 a6190dc7-2a80-4be9-9ff5-92f8de6ca74f |
| | | | | | | | gnl\|SRA\|SRR24090881.1761432.1:1185-1276 4dde59d0-00ef-42de-85a3-1e0d623cb57e |
| | | | | | | | gnl\|SRA\|SRR24090881.1761646.1:3328-3420 861cc5b1-170b-45c7-9244-fd52761e53f6 |
| | | | | 287 | 285 | Premature stop codon | gnl\|SRA\|SRR24090881.134977.1:2213-2300 b5861bb1-b4c1-4033-bb18-c52f46e44ed2 |

| | | | | | |
|---|---|---|---|---|---|
| | | | | | gnl\|SRA\|SRR24090881.1761432.1:438-527 4dde59d0-00ef-42de-85a3-1e0d623cb57e |
| | | | | | gnl\|SRA\|SRR24090881.1761646.1:4031-4121 861cc5b1-170b-45c7-9244-fd52761e53f6 |
| | | 402 | 398 | 2bp deletion | gnl\|SRA\|SRR24090881.1354404.1:4601-4681 556b3e9a-fd7b-4f22-b768-20b275a1c6fe |
| | | | | | gnl\|SRA\|SRR24090881.1838683.1:3920-3996 b5465727-072b-4824-aa48-3d54cf5f5694 |
| | | 602 | 597 | Premature stop codon | gnl\|SRA\|SRR24090881.2128931.1:1550-1626 e844d204-b979-44ea-a8c0-e6c9399a2785 |
| | | | | | gnl\|SRA\|SRR24090881.2422364.1:3543-3614 2e54a645-7793-4ec2-8069-3ec8d67f840c |
| | | 617 | 612 | Premature stop codon | gnl\|SRA\|SRR24090881.2128931.1:1730-1810 e844d204-b979-44ea-a8c0-e6c9399a2785 |
| | | | | | gnl\|SRA\|SRR24090881.2422364.1:3358-3442 2e54a645-7793-4ec2-8069-3ec8d67f840c |
| *Dugong dugon* | In this study | 244 | 242 | 1bp deletion | lcl\|V350022740L4C006R0601276725 988 966 1190/2 14012874 1 |
| | | | | | lcl\|V350022740L4C003R0280411238 121 1127 1196/1 2305988 1 |
| | | | | | lcl\|V350022740L4C001R0200652853 616 1283 80/2 9046084 1 |
| | | 250 | 248 | 1bp deletion | lcl\|V350022740L4C006R0601276725 988 966 1190/2 14012874 1 |
| | | | | | lcl\|V350022740L4C003R0280411238 121 1127 1196/1 2305988 1 |
| | | | | | lcl\|V350022740L4C001R0200652853 616 1283 80/2 9046084 1 |
| | | 287 | 285 | Premature stop codon | lcl\|V350022740L4C004R0190671377 560 827 311/1 11917685 1 |

| | | | | | | | |
|---|---|---|---|---|---|---|---|
| | | | | | | | lcl\|V350022740L4C006R0350324301 1496 1279 1089/2 24698297 1 |
| | | | | | | | lcl\|V350022740L4C006R0350324302 1496 1279 1089/2 24698297 1 |
| | | | | 398 | 394 | Premature stop codon | lcl\|V350022740L4C006R0320304002 708 692 942/1 980953 1 |
| | | | | | | | lcl\|V350022740L4C003R0451381703 11 186 199/1 8311103 1 |
| | | | | | | | lcl\|V350022740L4C002R0600826287 0 0 0/2 0 1 |
| | | | | 401 | 397 | 2bp deletion | lcl\|V350022740L4C006R0170239380 166 1373 1397/1 23523884 1 |
| | | | | | | | lcl\|V350022740L4C003R0451381703 11 186 199/1 8311103 1 |
| | | | | | | | lcl\|V350022740L4C002R0600826287 0 0 0/2 0 1 |
| *AWAT2* | Acyl-CoA wax alcohol acyltransferase 2 | *Trichechus manatus* | NCBI | 253 | 227 | Premature stop codon | gnl\|SRA\|SRR24090881.1134039.1:423-505 3737ce92-9166-4a3f-b2a7-232a8be74554 |
| | | | | | | | gnl\|SRA\|SRR24090881.626359.1:2491-2573 f388799f-2867-43ce-83ae-141757ab0530 |
| | | | | | | | gnl\|SRA\|SRR24090881.718277.1:3960-4041 8cd9ba8c-8a3a-4afc-817d-f98535597d19 |
| | | | | 347 | 317 | Premature stop codon | gnl\|SRA\|SRR24090881.718277.1:3476-3577 8cd9ba8c-8a3a-4afc-817d-f98535597d19 |
| | | *Dugong dugon* | In this study | 347 | 317 | Premature stop codon | lcl\|V350022740L4C002R0590279540 1104 1108 1171/2 3445948 1 |
| | | | | | | | lcl\|V350022740L4C005R0521054638 0 0 0/2 0 1 |

| | | | | | | | |
|---|---|---|---|---|---|---|---|
| | | | | | | | lcl\|V350022740L4C005R0530501728 788 647 614/1 6886881 1 |
| *CYP4V2* | Cytochrome P450 family 4 subfamily V member 2 | *Trichechus manatus* | NCBI | 124 | 124 | Premature stop codon | gnl\|SRA\|SRR24090881.3233497.1:560-647 b0de4a41-ce77-49a7-8cf4-a14b64a50120 |
| | | | | | | | gnl\|SRA\|SRR24090881.1216350.1:6178-6264 7f574ebf-f042-4d6c-945a-65edbdc08653 |
| | | | | | | | gnl\|SRA\|SRR24090881.474186.1:4698-4786 7d959e0d-c31c-4205-89b8-86a62454ca22 |
| | | | | 189 | 189 | 1bp deletion | gnl\|SRA\|SRR24090881.474186.1:5665-5751 7d959e0d-c31c-4205-89b8-86a62454ca22 |
| | | | | | | | gnl\|SRA\|SRR24090877.196853.1:3097-3185 1e4a5e54-ae36-416f-9c0b-8da47c1b5041 |
| | | | | | | | gnl\|SRA\|SRR24090877.1387023.1:3268-3341 4d1fccd1-29de-4215-9955-60c31ffc2cbd |
| | | | | 255 | 255 | 1bp deletion | gnl\|SRA\|SRR24090881.2777790.1:1275-1363 2c7376d8-1fc2-4ef7-b72d-51e3847ba927 |
| | | | | | | | gnl\|SRA\|SRR24090881.1216350.1:2450-2547 7f574ebf-f042-4d6c-945a-65edbdc08653 |
| | | | | | | | gnl\|SRA\|SRR24090881.1005692.1:734-817 897d625c-ac2c-41f1-85e9-ca2cff65089e |
| | | | | 341 | 340 | Premature stop codon | gnl\|SRA\|SRR24090881.2510094.1:632-707 aa41ba86-7d55-420e-8b10-a549fa8acd79 |
| | | | | | | | gnl\|SRA\|SRR24090881.2525700.1:4622-4696 d962943b-a384-4723-a528-2c6c36f67d2a |
| | | *Dugong dugon* | In this study | 35 | 35 | Premature stop codon | lcl\|V350045638L2C002R0260375800/2 |
| | | | | | | | lcl\|V350045638L3C006R0460574430/2 |
| | | | | 63 | 63 | 11bp deletion | lcl\|V350045638L2C002R0260375800/1 |

| | | | | | | | |
|---|---|---|---|---|---|---|---|
| | | | | | | | lcl\|V350045638L3C006R0460574430/1 |
| | | | | | | | lcl\|V350022740L4C001R0170383165 989 1289 217/1 7930067 1 |
| | | | | 75 | 75 | Premature stop codon | lcl\|V350022740L4C005R0220717276 556 847 426/1 1833105 1 |
| | | | | | | | lcl\|V350022740L4C006R0551126876 127 1327 1044/1 10559855 1 |
| | | | | | | | lcl\|V350022740L4C006R0620817048 127 1327 1044/1 10559855 1 |
| | | | | 153 | 153 | 1bp deletion | lcl\|V350022740L4C004R0300942989 121 953 873/1 3033283 1 |
| | | | | | | | lcl\|V350022740L4C006R0410164571 121 953 873/1 3033283 1 |
| | | | | | | | lcl\|V350022740L4C004R0600368538 149 40 965/1 8272752 1 |
| | | | | 169 | 169 | Premature stop codon | lcl\|V350022740L4C003R0350500529 557 525 1405/2 5733591 1 |
| | | | | | | | lcl\|V350022740L4C004R0130724479 890 1158 1457/1 19626387 1 |
| | | | | | | | lcl\|V350022740L4C004R0461102441 448 399 888/1 19671972 1 |
| | | | | 238 | 238 | Premature stop codon | lcl\|V350022740L4C003R0440358009 1438 1066 789/1 6595887 1 |
| | | | | | | | lcl\|V350022740L4C004R0290139025 1268 429 922/1 12569554 1 |
| | | | | | | | lcl\|V350022740L4C002R0601000874 1162 20 697/2 9027771 1 |
| *TAS2R10* | Taste 2 receptor member 10 | *Trichechus manatus* | NCBI | 102 | 101 | 1bp deletion | gnl\|SRA\|SRR24090881.582354.1:3632-3731 3e2aa9cd-b4ef-4927-b835-f935c7eb52ff |
| | | | | | | | gnl\|SRA\|SRR24090881.476537.1:3374-3473 ced7f491-f026-42b1-ac64-6f52d9fd63e0 |

| | | | | | |
|---|---|---|---|---|---|
| | | | | | gnl\|SRA\|SRR24090881.1831221.1:1619-1717 547205de-13c6-4a1c-81de-2aa0606b4003 |
| | | 152 | 151 | 1bp deletion | gnl\|SRA\|SRR24090881.1287309.1:2453-2551 cad609ca-d5f0-4775-a1c1-fa5668d10218 |
| | | | | | gnl\|SRA\|SRR24090881.1866821.1:6276-6375 58fa172e-7c9a-44af-9e29-08674d648fb7 |
| | | | | | gnl\|SRA\|SRR24090881.1831221.1:1466-1567 547205de-13c6-4a1c-81de-2aa0606b4003 |
| | | 183 | 181 | 2bp deletion | gnl\|SRA\|SRR24090881.2861264.1:4365-4469 14a685e8-151f-488a-8ca6-cb29a58ccb93 |
| | | | | | gnl\|SRA\|SRR24090881.1831221.1:1366-1469 547205de-13c6-4a1c-81de-2aa0606b4003 |
| | | | | | gnl\|SRA\|SRR24090881.1365689.1:2751-2852 643776fa-e9ca-44cb-a40c-a391232e0042 |
| | | 189 | 186 | 1bp deletion | gnl\|SRA\|SRR24090881.2861264.1:4379-4473 14a685e8-151f-488a-8ca6-cb29a58ccb93 |
| | | | | | gnl\|SRA\|SRR24090881.1831221.1:1362-1455 547205de-13c6-4a1c-81de-2aa0606b4003 |
| | | | | | gnl\|SRA\|SRR24090881.1287309.1:2351-2441 cad609ca-d5f0-4775-a1c1-fa5668d10218 |
| | | 216 | 213 | 1bp deletion | gnl\|SRA\|SRR24090881.1365689.1:2662-2761 643776fa-e9ca-44cb-a40c-a391232e0042 |
| | | | | | gnl\|SRA\|SRR24090881.1866821.1:6084-6184 58fa172e-7c9a-44af-9e29-08674d648fb7 |
| | | | | | gnl\|SRA\|SRR24090881.476537.1:3042-3140 ced7f491-f026-42b1-ac64-6f52d9fd63e0 |
| *Dugong dugon* | In this study | 14 | 14 | 1bp deletion | lcl\|V350022740L4C006R0271286399 1338 1488 891/2 10641033 1 |
| | | | | | lcl\|V350022740L4C002R0170560630 651 1175 1179/1 20126990 1 |
| | | | | | lcl\|V350022740L4C004R0100693650 0 0 0/1 0 1 |

| | | | | | | | |
|---|---|---|---|---|---|---|---|
| | | | | 63 | 62 | 1bp deletion | lcl\|V350022740L4C002R0161244670 555 1519 342/1 2673339 1 |
| | | | | | | | lcl\|V350022740L4C001R0630933971 438 1023 794/2 3919153 1 |
| | | | | | | | lcl\|V350022740L4C004R0620921059 1192 1315 998/1 14513772 1 |
| | | | | 102 | 101 | 1bp deletion | lcl\|V350022740L4C005R0470999188 520 1300 677/2 26615170 1 |
| | | | | | | | lcl\|V350022740L4C005R0470999187 520 1300 677/2 26615170 1 |
| | | | | | | | lcl\|V350022740L4C002R0161244670 555 1519 342/2 2673339 1 |
| | | | | 152 | 151 | 1bp deletion | lcl\|V350022740L4C005R0660118472 250 156 1117/1 16651568 1 |
| | | | | | | | lcl\|V350022740L4C001R0560956785 250 156 1117/1 16651568 1 |
| | | | | | | | lcl\|V350022740L4C005R0140613080 1227 241 997/1 1182965 1 |
| | | | | 189 | 186 | 1bp deletion | lcl\|V350022740L4C001R0120645076 552 1145 1274/1 5510830 1 |
| | | | | | | | lcl\|V350022740L4C001R0120645077 552 1145 1274/1 5510830 1 |
| | | | | | | | lcl\|V350022740L4C006R0690328385 1232 455 990/2 19303307 1 |
| | | | | 216 | 213 | 1bp deletion | lcl\|V350022740L4C002R0501395451 1217 522 1411/2 183140 1 |
| | | | | | | | lcl\|V350022740L4C002R0220012088 555 1519 342/1 2673339 1 |
| | | | | | | | lcl\|V350022740L4C004R0150760224 731 634 856/1 2908851 1 |
| *ABCG8* | ATP binding cassette subfamily G member 8 | *Trichechus manatus* | NCBI | 366 | 366 | Premature stop codon | SRR24090881.492313.1:3855-3941_c4523a32-9a12-4720-b38e-a245952b256f |

<table>
<tr><td></td><td></td><td></td><td></td><td></td><td>SRR24090881.1807049.1:1273-1364_89a71e54-1313-4bcc-8355-63975ade3123</td></tr>
<tr><td></td><td></td><td>566</td><td>566</td><td>1bp deletion</td><td>SRR24090881.467291.1:2718-2836_f6271214-4863-451e-b938-1d161ff81ae1</td></tr>
<tr><td></td><td></td><td></td><td></td><td></td><td>SRR24090881.2713839.1:1488-1606_92c9a6f7-153a-4635-a280-b5e4b28de57f</td></tr>
<tr><td></td><td></td><td>591</td><td>591</td><td>1bp deletion</td><td>SRR24090881.1580549.1:123-196_808e2d60-a38b-495d-acd2-e802d08837fd</td></tr>
<tr><td></td><td></td><td></td><td></td><td></td><td>SRR24090881.467291.1:1196-1273_f6271214-4863-451e-b938-1d161ff81ae1</td></tr>
<tr><td></td><td></td><td>598</td><td>598</td><td>Premature stop codon</td><td>SRR24090881.467291.1:1176-1273_f6271214-4863-451e-b938-1d161ff81ae1</td></tr>
<tr><td></td><td></td><td></td><td></td><td></td><td>SRR24090877.2703917.1:8860-8957 d03f0767-63de-4c89-ad99-131bad005cd5</td></tr>
<tr><td></td><td></td><td></td><td></td><td></td><td>SRR24090877.1309495.1:2526-2623 4bf2e05e-5541-4a4b-81e4-59de3a03209b</td></tr>
<tr><td>Dugong dugon</td><td>In this study</td><td>24</td><td>24</td><td>1bp deletion</td><td>lcl|V350022740L4C001R0320737975791_651_731/2 12331029 1</td></tr>
<tr><td></td><td></td><td></td><td></td><td></td><td>lcl|V350022740L4C005R0640430869906_423_352/2_14649527___1</td></tr>
<tr><td></td><td></td><td></td><td></td><td></td><td>lcl|V350022740L4C003R0180205924535_25_247/1_6356554___1</td></tr>
<tr><td></td><td></td><td>50</td><td>50</td><td>Premature stop codon</td><td>lcl|V350022740L4C006R0581372540535_25_247/2_6356554___1</td></tr>
<tr><td></td><td></td><td></td><td></td><td></td><td>lcl|V350022740L4C001R0050230209349_712_1225/1_2091935___1</td></tr>
<tr><td></td><td></td><td></td><td></td><td></td><td>lcl|V350022740L4C004R05703284770_0_0/1_0___1</td></tr>
<tr><td></td><td></td><td>121</td><td>121</td><td>Premature stop codon</td><td>lcl|V350022740L4C004R0460339254757_1208_1143/2 1321034 1</td></tr>
<tr><td></td><td></td><td></td><td></td><td></td><td>lcl|V350022740L4C004R069109146555_707_731/2 7813139 1</td></tr>
</table>

| | | | | | | | |
|---|---|---|---|---|---|---|---|
| | | | | | | | lcl\|V350022740L4C003R0100258589625_260_960/2 20673995 1 |
| | | | | 251 | 251 | 1bp deletion | lcl\|V350022740L4C005R0550430435525_1473_929/2_9303675___1 |
| | | | | | | | lcl\|V350022740L4C001R05413330621259_354_700/2_9664374___1 |
| | | | | | | | lcl\|V350022740L4C006R0371072101359_1317_279/2_31525092___1 |
| | | | | 294 | 294 | 1bp deletion | lcl\|V350022740L4C005R0450041991201_365_35/2 16300674 1 |
| | | | | | | | lcl\|V350022740L4C004R0620116039483_298_564/1 22425969 1 |
| | | | | | | | lcl\|V350022740L4C002R01801678421063_400_1103/2 4226999 1 |
| | | | | 440 | 440 | 1bp deletion | lcl\|V350022740L4C005R0071075318815_1145_570/2 4788003 1 |
| | | | | | | | lcl\|V350022740L4C001R0540990014959_101_772/1 8433094 1 |
| | | | | | | | lcl\|V350022740L4C003R0711068942331_1425_713/1 26747786 1 |
| | | | | 529 | 529 | 1bp deletion | lcl\|V350022740L4C003R04905536191276_1460_147/2 15328936 1 |
| | | | | | | | lcl\|V350022740L4C006R0620090676772_536_990/2 32013489 1 |
| | | | | | | | lcl\|V350022740L4C005R0221120868136_802_743/1 3794376 1 |
| *LYG2* | Lysozyme g2 | *Trichechus manatus* | NCBI | 14 | 14 | 1bp deletion | SRR24090881.889585.1:10311-10418_82de222d-3a74-44ef-ae93-4156f5409a94 |
| | | | | 31 | 31 | 1bp deletion | SRR24090881.889585.1:10311-10426_82de222d-3a74-44ef-ae93-4156f5409a94 |
| | | | | 80 | 80 | Premature stop codon | SRR24090877.2964499.1:1510-1624_2a565935-c669-49ce-a1d6-ac7f53675921 |

| | | | | | | | |
|---|---|---|---|---|---|---|---|
| | | | | | | | SRR24090877.2048204.1:3200-3318_0dba5312-78cd-48e2-a337-fe9016eea899 |
| | | *Dugong dugon* | In this study | 14 | 14 | 1bp deletion | lcl\|V350022740L4C004R0630004754787_485_1500/2 6562755 1 |
| | | | | | | | lcl\|V350022740L4C001R0620446757787_485_1500/2 6562755 1 |
| | | | | | | | lcl\|V350022740L4C006R0340002197170_1147_228/2 14969431 1 |
| | | | | 31 | 31 | 2bp deletion | lcl\|V350022740L4C004R0630004754787_485_1500/2 6562755 1 |
| | | | | | | | lcl\|V350022740L4C001R0620446757787_485_1500/2 6562755 1 |
| | | | | | | | lcl\|V350022740L4C006R0340002197170_1147_228/2 14969431 1 |
| | | | | 80 | 80 | Premature stop codon | lcl\|V350022740L4C006R05504649051223_114_886/2 18788571 1 |
| | | | | | | | lcl\|V350022740L4C004R07110027681378_404_743/1 1541457 1 |
| | | | | | | | lcl\|V350022740L4C003R0350156974631_48_846/2 25365908 1 |
| | | | | 163 | 163 | 1bp deletion | lcl\|V350022740L4C005R0600712128140_161_625/2 4941464 1 |
| | | | | | | | lcl\|V350022740L4C003R0100087181937_658_691/2 5138768 1 |
| | | | | | | | lcl\|V350022740L4C002R0170044614202_827_1164/1 20183689 1 |
| *NOX5* | NADPH oxidase 5 | *Trichechus manatus* | NCBI | 38 | 38 | 1bp deletion | SRR24090881.222725.1:4792-4909_56cf146e-6d13-4a67-85ab-30820f91e4b7 |
| | | | | | | | SRR24090881.2670743.1:6902-7023_c6534f8a-900a-42d5-994d-50e624056d96 |
| | | | | | | | SRR24090881.666546.1:6349-6446_2fa0ca16-b5e4-45eb-859e-ba6948cdd651 |

| | | | | | |
|---|---|---|---|---|---|
| | | 43 | 43 | Premature stop codon | SRR24090881.2670743.1:6900-7004_c6534f8a-900a-42d5-994d-50e624056d96 |
| | | 239 | 239 | 3bp deletion | SRR24090881.1566703.1:8531-8649_40491111-eeb3-456e-8f7b-f377065444fe |
| | | | | | SRR24090881.1847852.1:2768-2879_8c14a655-21af-478c-a43c-5b9e73102c0c |
| | | 422 | 422 | Premature stop codon | SRR24090881.1449708.1:8307-8426_9f447d95-afb5-4894-87f1-0f4b2e2d9de0 |
| | | | | | SRR24090881.1566703.1:4873-4988_40491111-eeb3-456e-8f7b-f377065444fe |
| | | | | | SRR24090881.1394305.1:12641-12758_c5c785a8-265a-4a76-9cf6-388ff5be17ae |
| | | 582 | 580 | 5bp deletion | SRR24090881.410965.1:4362-4447_a83b0880-7b22-4db7-a265-864eb0c82b5d |
| | | | | | SRR24090881.892896.1:1428-1519_a4fede83-a3be-4f78-81af-a7a02915c873 |
| | | | | | SRR24090881.1111577.1:2125-2212_c158b246-78df-4aaf-b3dc-8d7b0d0844b6 |
| | | 644 | 641 | Premature stop codon | SRR24090881.1111577.1:3304-3387_c158b246-78df-4aaf-b3dc-8d7b0d0844b6 |
| | | | | | SRR24090881.892896.1:2609-2692_a4fede83-a3be-4f78-81af-a7a02915c873 |
| | | | | | SRR24090881.1580668.1:2918-3002_eededf18-1f2f-48dc-b97f-43650c5e2cce |
| *Dugong dugon* | In this study | 38 | 38 | 1bp deletion | lcl\|V350022740L4C005R01504638561430_574_568/2 10150782 1 |
| | | | | | lcl\|V350022740L4C002R03508801181430_574_568/1 10150782 1 |
| | | | | | lcl\|V350022740L4C002R0721352340535_483_1161/2 12622926 1 |
| | | 43 | 43 | Premature stop codon | lcl\|V350022740L4C005R01504638561430_574_568/2 10150782 1 |

| | | | | | | | | |
|---|---|---|---|---|---|---|---|---|
| | | | | | | | lcl\|V350022740L4C002R03508801181430_574_568/1 10150782 | 1 |
| | | | | | | | lcl\|V350022740L4C002R0721352340535_483_1161/2 12622926 | 1 |
| | | | | 227 | 227 | 1bp deletion | lcl\|V350022740L4C002R0370226378978_942_88/1 18353882 | 1 |
| | | | | | | | lcl\|V350022740L4C004R0230005085621_169_1000/2 1208321 | 1 |
| | | | | | | | lcl\|V350022740L4C006R00613024061407_1411_164/2 1923389 | 1 |
| | | | | 371 | 371 | Premature stop codon | lcl\|V350022740L4C006R0441091444858_382_1040/2 13213927 | 1 |
| | | | | | | | lcl\|V350022740L4C006R0680531256496_1081_801/1 23127323 | 1 |
| | | | | | | | lcl\|V350022740L4C005R0120324770858_382_1040/2 13213927 | 1 |
| | | | | 582 | 580 | 17bp deletion | lcl\|V350022740L4C003R0230577488258_787_1060/2 7761448 | 1 |
| | | | | | | | lcl\|V350022740L4C004R0050256509611_327_91/2 1511219 | 1 |
| | | | | | | | lcl\|V350022740L4C003R0011260896258_787_1060/2 7761448 | 1 |
| | | | | 644 | 641 | Premature stop codon | lcl\|V350022740L4C006R0220736468182_711_197/2 21778579 | 1 |
| | | | | | | | lcl\|V350022740L4C004R0201078004182_711_197/2 21778579 | 1 |
| | | | | | | | lcl\|V350022740L4C001R03612820631203_1101_38/2 6837605 | 1 |
| *TCHH* | Trichohyalin | *Trichechus manatus* | NCBI | NA | NA | NA | No BLAST results | |
| | | *Dugong dugon* | This study | NA | NA | NA | No BLAST results | |

| | | | | | | | |
|---|---|---|---|---|---|---|---|
| *TCHHL1* | Trichohyalin like 1 | *Trichechus manatus* | NCBI | 91 | 91 | 3bp deletion | SRR24090881.593511.1:1614-1732 f3f7b9bc-fbd6-4e75-80c0-00689f268229 |
| | | | | | | | SRR24090881.582242.1:2345-2462 58dd009f-c4d5-4598-a6e0-c6930ae7abbb |
| | | | | | | | SRR24090881.1907941.1:2711-2828 574ffe90-66dc-4b9a-b35c-d07dc1dc060a |
| | | | | 292 | 290 | 1bp deletion | SRR24090877.127710.1:1409-1530 856ee87f-9c9f-417d-8989-0ec02751b6f6 |
| | | | | | | | SRR24090877.2139795.1:715-834 688d6b92-f2df-4dc8-a67a-dca439450afd |
| | | | | 336 | 334 | Premature stop codon | SRR24090877.291500.1:259-360 5a5412db-1798-4503-b501-649cbd423b75 |
| | | | | | | | SRR24090877.2146147.1:5343-5440 d8356c3e-6746-4b1b-b63b-1ebc2edfef97 |
| | | | | | | | SRR24090877.127710.1:1301-1397 856ee87f-9c9f-417d-8989-0ec02751b6f6 |
| | | | | 419 | 408 | 4bp deletion | SRR24090877.4054054.1:4125-4239 c6a52d1e-1387-4ad9-b16a-7cb52b67d0f1 |
| | | | | | | | SRR24090877.127710.1:959-1080 856ee87f-9c9f-417d-8989-0ec02751b6f6 |
| | | | | | | | SRR24090877.2146147.1:5663-5773 d8356c3e-6746-4b1b-b63b-1ebc2edfef97 |
| | | | | 639 | 628 | Premature stop codon | SRR24090877.4054054.1:3468-3587 c6a52d1e-1387-4ad9-b16a-7cb52b67d0f1 |
| | | | | | | | SRR24090877.127710.1:306-413 856ee87f-9c9f-417d-8989-0ec02751b6f6 |
| | | | | 748 | 737 | 6bp deletion | SRR24090877.751664.1:3917-4028 9b54491c-90e5-4d6e-b8e9-728d5b4b4009 |
| | | | | | | | SRR24090877.454906.1:6735-6844 103acdf6-aced-421f-8597-9d8f8a407be3 |
| | | | | | | | SRR24090877.291500.1:1536-1645 5a5412db-1798-4503-b501-649cbd423b75 |

| | | | | | |
|---|---|---|---|---|---|
| | | 766 | 755 | 3bp deletion | SRR24090877.4054054.1:3099-3209 c6a52d1e-1387-4ad9-b16a-7cb52b67d0f1 |
| | | | | | SRR24090877.454906.1:6681-6792 103acdf6-aced-421f-8597-9d8f8a407be3 |
| | | | | | SRR24090877.751664.1:3864-3974 9b54491c-90e5-4d6e-b8e9-728d5b4b4009 |
| | | 776 | 765 | 1bp deletion | SRR24090877.4054054.1:3070-3184 c6a52d1e-1387-4ad9-b16a-7cb52b67d0f1 |
| | | | | | SRR24090877.454906.1:6651-6768 103acdf6-aced-421f-8597-9d8f8a407be3 |
| | | | | | SRR24090877.751664.1:3834-3950 9b54491c-90e5-4d6e-b8e9-728d5b4b4009 |
| *Dugong dugon* | In this study | 225 | 225 | Premature stop codon | lcl\|V350022740L4C006R0040553655*122_331_857/1_17728807___1 |
| | | | | | lcl\|V350022740L4C004R0590106634*362_328_448/1_3700768___1 |
| | | | | | lcl\|V350022740L4C006R0640428796*749_28_117/1_1819407___1 |
| | | 261 | 261 | Premature stop codon | lcl\|V350022740L4C003R0030517510*697_777_235/1_4356351___1 |
| | | | | | lcl\|V350022740L4C005R0300467636*697_777_235/2_4356351___1 |
| | | | | | lcl\|V350022740L4C005R0220697492*1451_715_20/2_8518967___1 |
| | | 286 | 284 | Premature stop codon | lcl\|V350022740L4C005R0400131047*547_904_646/2_19181861___1 |
| | | | | | lcl\|V350022740L4C004R0090237104*547_904_646/2_19181861___1 |
| | | | | | lcl\|V350022740L4C004R0090237103*547_904_646/2_19181861___1 |
| | | 292 | 290 | 1bp deletion | lcl\|V350022740L4C005R0400131047*547_904_646/2_19181861___1 |

| | | | | |
|---|---|---|---|---|
| | | | | lcl\|V350022740L4C004R0090237104*547_904_646/2_19181861___1 |
| | | | | lcl\|V350022740L4C004R0090237103*547_904_646/2_19181861___1 |
| | 336 | 334 | Premature stop codon | lcl\|V350022740L4C005R0650252984*312_865_177/1_1925661___1 |
| | | | | lcl\|V350022740L4C005R0361355761*1128_1175_1441/1_1572113___1 |
| | | | | lcl\|V350022740L4C002R0360669680*665_562_615/1_133250___1 |
| | 419 | 408 | 4bp deletion | lcl\|V350022740L4C003R0060827754*556_311_1464/2_16753221___1 |
| | | | | lcl\|V350022740L4C001R0580030656*556_311_1464/2_16753221___1 |
| | | | | lcl\|V350022740L4C001R0041268584*312_865_177/2_1925661___1 |
| | 482 | 471 | Premature stop codon | lcl\|V350022740L4C004R0301400145*1496_736_862/2_3241690___1 |
| | | | | lcl\|V350022740L4C001R0620816263*33_4_1428/2_4853777___1 |
| | | | | lcl\|V350022740L4C001R0100180359*33_4_1428/2_4853777___1 |
| | 526 | 515 | 22bp deletion | lcl\|V350022740L4C005R0500382131*36_1237_345/2_2597088___1 |
| | | | | lcl\|V350022740L4C005R0500381914*36_1237_345/2_2597088___1 |
| | | | | lcl\|V350022740L4C004R0410782889*814_1010_1219/2_10708709___1 |
| | 542 | 530 | 1bp deletion | lcl\|V350022740L4C005R0050557921*1223_476_345/2_14730919___1 |
| | | | | lcl\|V350022740L4C004R0410782889*814_1010_1219/2_10708709___1 |

| | | | |
|---|---|---|---|
| | | | lcl\|V350022740L4C004R0410782672*814_1010_1219/2_10708709___1 |
| 634 | 623 | 4bp deletion | lcl\|V350022740L4C003R0140167048362_328_448/2 3700768 1 |
| | | | lcl\|V350022740L4C004R05313163051018_626_788/2 6678237 1 |
| | | | lcl\|V350022740L4C003R0310538939905_405_1152/2 11461048 1 |
| 639 | 628 | 1bp deletion | lcl\|V350022740L4C003R0140167048362_328_448/2 3700768 1 |
| | | | lcl\|V350022740L4C004R05313163051018_626_788/2 6678237 1 |
| | | | lcl\|V350022740L4C003R0310538939905_405_1152/2 11461048 1 |
| 717 | 706 | 8bp deletion | lcl\|V350022740L4C005R01805901601128_1175_1441/1 1572113 1 |
| | | | lcl\|V350022740L4C004R0501067095984_655_415/2 608753 1 |
| | | | lcl\|V350022740L4C002R0700079116117_1531_481/1 23696059 1 |
| 748 | 737 | 6bp deletion | lcl\|V350022740L4C005R0030032965516_1297_125/2 348275 1 |
| | | | lcl\|V350022740L4C001R0190074996964_332_653/1 1566577 1 |
| | | | lcl\|V350022740L4C005R0021196539234_814_1141/1 11037368 1 |
| 766 | 755 | 3bp deletion | lcl\|V350022740L4C006R053019134127_1116_362/2 7740812 1 |
| | | | lcl\|V350022740L4C002R04404190543_582_1098/1 12395041 1 |
| | | | lcl\|V350022740L4C005R02402922261235_59_556/1 7161131 1 |

| | | | | | | | |
|---|---|---|---|---|---|---|---|
| | | | | 776 | 765 | 1bp deletion | lcl\|V350022740L4C006R053019134127_1116_362/2 7740812 1 |
| | | | | | | | lcl\|V350022740L4C002R04404190543_582_1098/1 12395041 1 |
| | | | | | | | lcl\|V350022740L4C005R02402922261235_59_556/1 7161131 1 |
| *CES3* | Carboxylesterase 3 | *Trichechus manatus* | NCBI | 50 | 44 | Premature stop codon | SRR8616893.230161756.1 230161756 |
| | | | | | | | SRR8616893.259113249.1 259113249 |
| | | | | | | | SRR8616893.259113249.1 259113249 |
| | | | | 438 | 430 | Premature stop codon | SRR24090880.2869722.1:706-766 9810fa93-f445-43b7-9c9c-16a8ec930c6a |
| | | | | | | | SRR24090880.102114.1:2536-2597 cc2e227e-da48-44f1-b1a2-d7b24fe1c779 |
| | | | | | | | SRR24090880.1708043.1:485-545 42680180-98ae-4cfa-bea9-fcd02b81aed3 |
| | | *Dugong dugon* | In this study | 300 | 294 | Premature stop codon | V350022740L4C005R0330397004*673_1354_715/2_12977957___1 |
| | | | | | | | V350022740L4C005R0330397004*673_1354_715/2_12977957___1 |
| | | | | | | | V350022740L4C002R0500799161*1354_1047_751/2_12614826___1 |
| | | | | 438 | 430 | Premature stop codon | V350022740L4C005R048137690123_1471_370/2_7002964___1 |
| | | | | | | | V350022740L4C006R05113786411432_1002_1396/2_4540938___1 |
| | | | | | | | V350022740L4C006R0511378641*1432_1002_1396/2_4540938___1 |
| | | | | 453 | 444 | Premature stop codon | V350022740L4C006R02802843161023_476_806/1 10432385 1 |

| | | | | | | | |
|---|---|---|---|---|---|---|---|
| | | | | | | | V350022740L4C006R014006050481_1356_1493/1 16283621 1 |
| | | | | | | | V350022740L4C003R0410760833*1290_1190_1440/1_8433771___1 |
| | | *Hydrodamalis gigas* | In this study | 438 | 430 | Premature stop codon | ERR5559486.9163196.2:3-64 9163196 |
| | | | | | | | ERR5559486.9163196.1:18-79 9163196 |
| | | | | | | | ERR5559486.9159397.2:3-64 9159397 |
| | | | | 453 | 444 | Premature stop codon | SRR12067494.177331381.1177331381 Biological (Biological) |
| | | | | | | | SRR12067494.123799503.2123799503 Biological (Biological) |
| | | | | | | | SRR12067494.123791255.2123791255 Biological (Biological) |
| | | | | 549 | 542 | 1bp deletion | gnl\|SRA\|SRR12067494.212087335.1:1-54 212087335 |
| | | | | | | | gnl\|SRA\|SRR12067494.225892124.2:1-54 225892124 |
| | | | | | | | gnl\|SRA\|SRR12067494.67273898.1:1-54 67273898 |
| *PON1* | Paraoxonase 1 | *Trichechus manatus* | NCBI | 150 | 123 | Premature stop codon | None |
| | | | | 226 | 199 | 1bp deletion | SRA\|SRR8616893.211392162.1 211392162 |
| | | | | | | | SRA\|SRR8616893.24449605.2 24449605 |
| | | | | | | | SRA\|SRR8616893.1354008.2 1354008 |
| | | | | 343 | 316 | Premature stop codon | gnl\|SRA\|SRR8616893.180900865.1 180900865 |

| | | | | | |
|---|---|---|---|---|---|
| | | | | | gnl\|SRA\|SRR8616893.137237637.2 137237637 |
| | | | | | gnl\|SRA\|SRR8616893.136191442.1 136191442 |
| *Dugong dugon* | In this study | 54 | 27 | Premature stop codon | lcl\|V350022740L4C004R0520897376 686 1291 137/2 9141215 1 |
| | | | | | lcl\|V350022740L4C005R0640460073 1232 280 968/1 915685 1 |
| | | 86 | 59 | 1bp deletion | lcl\|V350022740L4C003R0391360360*351_297_1453/2_7680740___1 |
| | | | | | lcl\|V350022740L4C003R0190755030*0_0_0/2_0___1 |
| | | | | | lcl\|V350022740L4C005R0400095956*1109_75_522/1_6122181___1 |
| | | 134 | 107 | 2bp deletion | lcl\|V350022740L4C003R0600961420*641_1020_883/2_11120702___1 |
| | | | | | lcl\|V350022740L4C004R0200326326*1501_294_824/2_5149806___1 |
| | | | | | lcl\|V350022740L4C004R0690408471*1192_1277_1292/2_4918045___1 |
| | | 150 | 123 | Premature stop codon | lcl\|V350022740L4C003R0521203135*1353_610_1057/2_10629209___1 |
| | | | | | lcl\|V350022740L4C004R0200326326*1501_294_824/2_5149806___1 |
| | | | | | lcl\|V350022740L4C001R0450445433*91_814_376/1_14888219___1 |
| | | 333 | 305 | 20bp deletion | lcl\|V350022740L4C006R0041211771*0_0_0/2_0___1 |
| | | | | | lcl\|V350022740L4C005R0581338690*1474_700_593/2_2140291___1 |

| | | | | | | | |
|---|---|---|---|---|---|---|---|
| | | | | | | | lcl\|V350022740L4C003R0690373184*1474_700_593/2_2140291___1 |
| *KCNK18* | Potassium two pore domain channel subfamily K member 18 | *Trichechus manatus* | NCBI | 280 | 55 | Premature stop codon | SRR24090881.22980.1:1487-1588 d1f68a3e-4baa-488b-8895-4d1afd66010f |
| | | | | | | | SRR24090881.2225469.1:4267-4369 6a36221a-2ee1-4c63-ab26-e6912975e99f |
| | | | | | | | SRR24090881.1863039.1:2610-2711 7ba8b66d-acb4-43b4-962d-d41953554864 |
| | | | | 408 | 183 | Premature stop codon | SRR24090881.1720750.1:2596-2687 c3b9a788-1896-4b70-acb2-7f9bfddd076b |
| | | | | | | | SRR24090881.1494441.1:199-292 5cc589b1-b12e-4d1d-a08f-a061a6f42d9a |
| | | | | | | | SRR24090881.2119242.1:1296-1393 e247809f-5dbe-4bdb-9e10-d128264a494e |
| | | | | 422 | 198 | 11bp deletion | gnl\|SRA\|SRR8616893.157018565.1:45-120 157018565 |
| | | | | | | | gnl\|SRA\|SRR8616893.71815200.1:45-120 71815200 |
| | | | | | | | gnl\|SRA\|SRR8616893.222332216.1:1-60 222332216 |
| | | | | 462 | 234 | Premature stop codon | SRR24090881.2119242.1:1138-1239 e247809f-5dbe-4bdb-9e10-d128264a494e |
| | | | | | | | SRR24090881.1720750.1:2442-2540 c3b9a788-1896-4b70-acb2-7f9bfddd076b |
| | | | | 502 | 274 | Premature stop codon | SRR24090881.1720750.1:2330-2430 c3b9a788-1896-4b70-acb2-7f9bfddd076b |
| | | | | | | | SRR24090881.1944218.1:841-929 22d69217-cfc4-4190-ab11-b9e6788d2456 |
| | | | | 507 | 279 | Premature stop codon | SRR24090880.908340.1:3896-3994 fe960a55-1a4a-4f75-8464-dc351ac421ce |

| | | | | | |
|---|---|---|---|---|---|
| *Dugong dugon* | In this study | 280 | 55 | Premature stop codon | V350022740L4C004R0660121241 979 1290 1344/2 9309378 1 |
| | | | | | V350022740L4C006R0520056691 1458 1250 418/1 11995360 1 |
| | | | | | V350022740L4C002R0671209433 1458 1250 418/1 11995360 1 |
| | | 422 | 198 | 11bp deletion | lcl\|V350022740L4C005R0671279219*1434_986_172/1_9699561___1 |
| | | | | | lcl\|V350022740L4C003R0190928973*1529_1414_379/2_5805673___1 |
| | | | | | lcl\|V350022740L4C002R0270696033*726_23_1302/1_15336326___1 |
| | | 526 | 301 | 1bp deletion | lcl\|V350022740L4C002R0421172253*762_545_704/1_19904563___1 |
| | | | | | lcl\|V350022740L4C003R0110160033*158_298_671/1_10709090___1 |
| | | | | | lcl\|V350022740L4C005R0020485777*946_856_676/1_13565442___1 |
| *Hydrodamalis gigas* | In this study | 280 | 55 | Premature stop codon | SRR12067500.29671061.129671061 Biological (Biological) |
| | | 422 | 198 | 11bp deletion | gnl\|SRA\|SRR12067494.130640529.1:1-51 130640529 |
| | | | | | gnl\|SRA\|SRR12067494.152965580.1:3-52 152965580 |
| | | | | | gnl\|SRA\|SRR12067494.103621824.1:3-52 103621824 |
| | | 458 | 230 | Premature stop codon | gnl\|SRA\|SRR12067494.220334659.2:1-54 220334659 |
| | | | | | gnl\|SRA\|SRR12067494.126759574.1:1-54 126759574 |
| | | 462 | 234 | Premature stop codon | ERR5559486.45902189.2:1-133 45902189 |
| | | | | | ERR5559486.108617651.1:1-133 108617651 |

| | | | | |
|---|---|---|---|---|
| | | | | ERR5559486.108617651.2:1-133 108617651 |
| | 526 | 301 | 1bp deletion | gnl\|SRA\|SRR12067494.99914576.1:1-54 99914576 |
| | | | | gnl\|SRA\|SRR12067494.58981731.2:1-54 58981731 |
| | | | | gnl\|SRA\|SRR12067494.143642195.2:1-54 143642195 |

**Table S10 | Dating sirenian gene loss events**

The table shows the $K_a/K_s$ value of the terminal branch leading to two sirenians (dugong and West Indian manatee) that have lost a gene (referred to as $K$) and the $K_a/K_s$ value for all other afrotherians (referred to as $K_s$) that have an intact gene. The lower and upper bound of the divergence time between these species and their closest sister species (referred to as $T$) was obtained from TimeTree [57]. Applying a method described by [58], a gene loss date was estimated as $T_n=T\times(K-K_s)/(1-K_s)$. The estimated divergence time of dugong and manatees from our dataset was 31.2 (27.4-37.0) Mya. A limitation of this method is that sirenian gene loss cannot be dated if a gene of interest is missing or heavily fragmented in any non-sirenian species in our afrotherian dataset.

| Gene | Function(s) | $K_a/K_s$ value entire branch ($K$) | $Ka/Ks$ value for species with a functional gene ($K_s$) | Estimated time of sirenian gene loss (Mya) |
|---|---|---|---|---|
| *ABCG8* | Diet | 0.36 | 0.15 | 7.73 (6.78-9.16) |
| *ALOX15* | Skin | 0.63 | 0.26 | 15.53 (13.64-18.41) |
| *AWAT2* | Skin | 1.17 | 0.69 | 47.78 (41.96-56.65) |
| *CES3* | Cholesterol ester metabolism<br>Xenobiotic clearance | 0.62 | 0.35 | 13.13 (11.52-15.56) |
| *CHIA5* | Diet | 0.86 | 0.20 | 25.70 (22.57-30.48) |
| *CYP4V2* | Vision | 0.92 | 0.22 | 27.81 (24.42-32.98) |
| *KCNK18* | Brain: Circadian clock<br>Skin: Cold- and xenobiotic-induced pain | 0.62 | 0.26 | 15.30 (13.43-18.15) |
| *LYG1* | Immunity (skin) | 0.68 | 0.32 | 16.40 (14.10-19.45) |
| *LYG2* | Skin | 0.50 | 0.35 | 7.44 (6.53-8.82) |
| *NOX5* | Oxidative stress | 0.37 | 0.12 | 8.77 (7.70-10.40) |
| *PON1* | Oxidative stress<br>Xenobiotic clearance | 0.79 | 0.25 | 22.32 (19.60-26.48) |
| *TAS2R10* | Taste (bitter) | 1.46 | 0.64 | 71.20 (62.52-84.43) |

**Table S11 | Sirenian-specific amino acid substitutions in thyroid hormone pathway and circadian clock genes**

Potential functional effects of substitutions were predicted using PolyPhen-2 and SIFT.

| Gene | Name | Sirenian Site | Human site | Foreground residue | Background residue | Radical change(s) | PolyPhen-2 | PolyPhen-2-score | SIFT | SIFT score | UniProt Feature site description |
|---|---|---|---|---|---|---|---|---|---|---|---|
| *CFTR* | CF transmembrane conductance regulator | 276 | 276 | R | C | charge | Probably damaging | 0.999 | Damaging | 0.01 | ABC transmembrane type-1 1 |
| | | 297 | 297 | W | R | charge | Probably damaging | 0.999 | Damaging | 0.01 | ABC transmembrane type-1 1 |
| | | 332 | 332 | T | I | polarity | Bening | 0.029 | Tolerated | 0.42 | ABC transmembrane type-1 1 |
| | | 393 | 393 | L | V | not_radical | Bening | 0.00 | Tolerated | 0.59 | |
| | | 730 | 730 | K | E | charge | Bening | 0.29 | Tolerated | 0.09 | |
| | | 781 | 779 | L | Q | polarity | Bening | 0.24 | Tolerated | 0.2 | |
| | | 974 | 971 | E | G | charge | Bening | 0.005 | Tolerated | 0.91 | ABC transmembrane type-1 2 |
| | | 1042 | 1039 | R | Q | charge | Probably damaging | 0.999 | Damaging | 0.03 | ABC transmembrane type-1 2 |
| | | 1282 | 1277 | L | I | not_radical | Bening | 0.000 | Tolerated | 0.1 | ABC transporter 2 |
| *DUOX2* | Dual oxidase 2 | 7 | 7 | V | E | charge | Bening | 0.392 | Damaging | 0.02 | |
| | | 29 | 29 | Q | L | polarity | Bening | 0.004 | Damaging | 0.02 | |
| | | 404 | 404 | T | N | polarity&volume | Bening | 0.002 | Tolerated | 0.3 | Peroxidase-like; mediates peroxidase activity |
| | | 451 | 451 | G | D | charge | Bening | 0.000 | Tolerated | 0.28 | Peroxidase-like; mediates peroxidase activity |
| | | 469 | 469 | T | E | charge | Bening | 0.035 | Damaging | 0.00 | Peroxidase-like; mediates peroxidase activity |
| | | 698 | 698 | A | S | polarity | Bening | 0.329 | Damaging | 0.00 | |
| | | 915 | 915 | E | D | not_radical | Bening | 0.012 | Tolerated | 0.62 | EF-hand 3 |
| | | 936 | 936 | C | R | charge | Probably damaging | 0.999 | Damaging | 0.01 | EF-hand 3 |
| | | 964 | 962 | R | C | charge | Bening | 0.000 | Tolerated | 0.41 | |
| | | 965 | 963 | Q | R | charge | Possibly damaging | 0.659 | Tolerated | 0.46 | |
| | | 1008 | 1006 | R | P | charge | Bening | 0.001 | Tolerated | 0.46 | Interaction with TXNDC11 |
| | | 1052 | 1050 | L | A | polarity&volume | Bening | 0.002 | Tolerated | 0.09 | Interaction with TXNDC11 |
| | | 1478 | 1476 | H | L | charge | Probably damaging | 0.965 | Tolerated | 0.32 | |

| | | | | | | | | | | | |
|---|---|---|---|---|---|---|---|---|---|---|---|
| *DUOXA2* | Dual oxidase maturation factor 2 | 146 | 146 | R | L | charge | Probably damaging | 0.999 | Damaging | 0.00 | |
| | | 279 | 279 | L | R | charge | Probably damaging | 1 | Tolerated | 0.11 | |
| *KCNE2* | Potassium voltage-gated channel subfamily E regulatory subunit 2 (LQT6) | 28 | 28 | M | R | charge | Probably damaging | 0.497 | Damaging | 0,01 | Slow voltage-gated potassium channel |
| | | 75 | 75 | R | K | not_radical | Probably damaging | 0.992 | Damaging | 0.05 | Slow voltage-gated potassium channel |
| *KCNQ1* | Potassium voltage-gated channel subfamily Q member 1 | 388 | 325 | G | D | charge | Probably damaging | 1.000 | Tolerated | 1.00 | Ion_trans |
| | | 429 | 366 | S | N | polarity&volume | Bening | 0.029 | Tolerated | 0.10 | |
| | | 660 | 583 | F | T | polarity | Bening | 0.266 | Damaging | 0.00 | |
| *LPO* | Lactoperoxidase | 42 | 42 | T | K | charge | Probably damaging | 0.998 | Tolerated | 0.13 | |
| | | 68 | 68 | G | R | charge | Bening | 0.305 | Tolerated | 0.39 | |
| | | 112 | 112 | V | L | not_radical | Bening | 0.025 | Tolerated | 0.53 | |
| | | 118 | 118 | C | S | polarity&volume | Probably damaging | 0.992 | Damaging | 0.03 | |
| | | 634 | 632 | V | L | not_radical | Bening | 0.143 | Tolerated | 0.15 | An_peroxidase |
| *SLC5A4* | Solute carrier family 5 member 4 | 413 | 412 | R | K | not_radical | Bening | 0.022 | Tolerated | 0.99 | SSF |
| *SLC5A5* | Solute carrier family 5 member 5 (NIS) | 140 | 142 | L | M | not_radical | Bening | 0.00 | Tolerated | 0.12 | SSF |
| | | 201 | 203 | G | W | polarity | Bening | 0.00 | Tolerated | 0.33 | SSF |
| | | 319 | 321 | A | P | not_radical | Bening | 0.387 | Tolerated | 0.12 | SSF |
| | | 443 | 445 | A | V | polarity&volume | Bening | 0.015 | Tolerated | 1.00 | SSF |
| | | 536 | 539 | S | T | not_radical | Probably damaging | 0.997 | Tolerated | 0.10 | |

| | | | | | | | | | | | |
|---|---|---|---|---|---|---|---|---|---|---|---|
| *SLC5A6* | Solute carrier family 5 member 6 | 28 | 26 | S | I | polarity | Bening | 0.004 | Tolerated | 0.08 | |
| | | 430 | 428 | F | I | polarity&volume | Bening | 0.003 | Tolerated | 0.21 | SSF |
| | | 522 | 520 | L | Q | polarity | Bening | 0 | Damaging | 0.04 | |
| *SLC5A8* | Solute carrier family 5 member 8 | 51 | 51 | G | R | charge | Bening | 0 | Tolerated | 0.07 | SSF |
| | | 93 | 93 | T | F | polarity | Bening | 0 | Tolerated | 0.52 | SSF |
| | | 171 | 171 | A | V | polarity&volume | Bening | 0.01 | Tolerated | 0.15 | SSF |
| | | 432 | 432 | T | I | polarity | Possibly damaging | 0.89 | Damaging | 0.02 | SSF |
| *TPO* | Thyroid peroxidase | 106 | 105 | I | V | not_radical | Possibly damaging | 0.483 | Tolerated | 0.08 | EPO_TPO |
| | | 279 | 278 | N | A | polarity | Possibly damaging | 0.65 | Tolerated | 0.12 | |
| | | 609 | 607 | V | A | polarity&volume | Possibly damaging | 0.488 | Tolerated | 0.09 | |
| *TSHR* | Thyroid stimulating hormone receptor | 18 | 18 | V | D | charge | Bening | 0 | Damaging | 0.04 | |
| | | 270 | 270 | V | L | not_radical | Bening | 0.015 | Tolerated | 1.00 | |
| | | 300 | 300 | I | M | not_radical | Bening | 0.007 | Tolerated | 0.36 | |
| | | 310 | 310 | G | R | charge | Possibly damaging | 0.734 | Tolerated | 0.41 | |
| | | 536 | 536 | K | R | not_radical | Bening | 0.259 | Tolerated | 0.37 | 7tm_1 |
| | | 736 | 736 | H | E | charge | Bening | 0 | Damaging | 0.01 | |
| | | 746 | 746 | N | H | charge | Bening | 0.201 | Tolerated | 0.05 | |
| *CLOCK* | Clock circadian regulator | 218 | 208 | A | S | polarity | Bening | 0 | Tolerated | 0.15 | |
| *CRY1* | Cryptochrome circadian regulator 1 | 167 | 167 | E | V | charge | Bening | 0.045 | Damaging | 0.03 | |
| | | 462 | 461 | S | L | polarity | Possibly damaging | 0.495 | Damaging | 0.03 | FAD_binding_7 |
| *CRY2* | Cryptochrome circadian regulator 2 | 527 | 526 | A | V | polarity&volume | Bening | 0.109 | Damaging | 0.00 | |
| *DBP* | D-box binding PAR bZIP | 59 | 59 | A | P | not_radical | Bening | 0.001 | Tolerated | 0.21 | |

| | | | | | | | | | | | |
|---|---|---|---|---|---|---|---|---|---|---|---|
| | transcription factor | | | | | | | | | | |
| | | 114 | 114 | L | V | not_radical | Possibly damaging | 0.62 | Tolerated | 1.00 | |
| | | 173 | 173 | V | A | polarity&volume | Possibly damaging | 0.844 | Tolerated | 0.06 | |
| *NALCN* | Sodium leak channel, non-selective | 393 | 393 | V | M | not_radical | Bening | 0.179 | Tolerated | 0.3 | Ion_trans |
| | | 852 | 852 | L | V | not_radical | Bening | 0 | Tolerated | 0.2 | |
| *NFIL3* | Nuclear factor, interleukin 3 regulated | 405 | 404 | I | S | polarity | Bening | 0.04 | Damaging | 0.05 | Vert_IL3-reg_TF |
| *PER1* | Period circadian regulator 1 | 179 | 179 | E | Q | charge | Possibly damaging | 0.865 | Tolerated | 1.00 | |
| | | 512 | 512 | G | V | polarity | Possibly damaging | 0.612 | Damaging | 0.00 | |
| | | 584 | 584 | Q | K | charge | Probably damaging | 0.993 | Damaging | 0.03 | |
| | | 634 | 634 | V | I | not_radical | Probably damaging | 0.734 | Tolerated | 0.46 | Required for phosphorylation by CSNK1E |
| | | 954 | 948 | L | P | polarity&volume | Probably damaging | 0.997 | Damaging | 0.04 | |
| *PER2* | Period circadian regulator 2 | 113 | 112 | R | I | charge | Probably damaging | 0.998 | Damaging | 0.00 | Nuclear export signal 1 |
| | | 182 | 181 | K | V | charge | Possibly damaging | 0.8 | Damaging | 0.00 | PAS 1 |
| | | 269 | 267 | G | E | charge | Probably damaging | 0.997 | Tolerated | 0.21 | |
| | | 457 | 455 | S | L | polarity | Possibly damaging | 0.946 | Tolerated | 0.22 | |
| | | 469 | 467 | C | H | charge | Probably damaging | 1 | Damaging | 0.00 | Nuclear export signal 2 |
| | | 478 | 476 | Y | H | charge | Probably damaging | 0.989 | Tolerated | 0.30 | |
| | | 581 | 572 | V | A | polarity&volume | Possibly damaging | 0.594 | Tolerated | 0.32 | CSNK1E binding domain |
| | | 658 | 648 | T | G | not_radical | Bening | 0.034 | Damaging | 0.03 | |
| | | 766 | 755 | S | Q | polarity&volume | Possibly damaging | 0.679 | Tolerated | 0.76 | |
| | | 770 | 759 | P | H | charge | Bening | 0.312 | Tolerated | 0.13 | |
| | | 827 | 815 | Q | G | polarity&volume | Possibly damaging | 0.822 | Damaging | 0.03 | |
| | | 862 | 850 | G | V | polarity | Probably damaging | 0.998 | Damaging | 0.01 | |
| | | 963 | 935 | I | A | polarity&volume | Possibly damaging | 0.647 | Damaging | 0.04 | Polar residues |
| | | 1014 | 970 | I | L | not_radical | Probably damaging | 0.999 | Tolerated | 0.07 | |

| | | | | | | | | | | | |
|---|---|---|---|---|---|---|---|---|---|---|---|
| | | 1071 | 1025 | R | P | charge | Possibly damaging | 0.766 | Damaging | 0.00 | |
| | | 1161 | 1107 | T | M | polarity | Bening | 0.159 | Tolerated | 0.15 | |
| | | 1164 | 1110 | H | G | charge | Possibly damaging | 0.901 | Damaging | 0.03 | |
| | | 1182 | 1128 | V | L | not_radical | Probably damaging | 0.999 | Tolerated | 0.13 | |
| | | 1197 | 1143 | T | L | polarity | Probably damaging | 0.976 | Damaging | 0.01 | |
| | | 1264 | 1210 | P | C | polarity | Possibly damaging | 0.952 | Tolerated | 0.16 | CRY binding domain |
| *PER3* | Period circadian regulator 3 | 252 | 220 | Y | C | polarity&volume | Probably damaging | 1.00 | Damaging | 0.04 | |
| | | 308 | 276 | M | I | not_radical | Probably damaging | 1.00 | Damaging | 0.01 | PAS 2 |
| | | 567 | 529 | D | N | charge | Probably damaging | 0.999 | Damaging | 0.03 | |
| | | 593 | 554 | T | L | polarity | Bening | 0.022 | Tolerated | 0.09 | |
| | | 672 | 627 | T | A | polarity | Probably damaging | 0.958 | Damaging | 0.02 | Region： CSNK1E binding domain |
| | | 725 | 680 | V | A | polarity&volume | Possibly damaging | 0.920 | Damaging | 0.02 | Region： CSNK1E binding domain |
| | | 829 | 780 | S | A | polarity | Bening | 0.090 | Tolerated | 0.65 | |
| | | 983 | 931 | I | S | polarity | Possibly damaging | 0.952 | Tolerated | 0.09 | |
| | | 1079 | 954 | T | A | polarity | Probably damaging | 1.00 | Damaging | 0.03 | |
| | | 1104 | 978 | M | Q | polarity | Possibly damaging | 0.949 | Tolerated | 0.39 | |
| | | 1118 | 992 | D | A | charge | Probably damaging | 0.986 | Damaging | 0.02 | |
| | | 1185 | 1059 | F | V | polarity&volume | Bening | 0.278 | Tolerated | 0.06 | |
| | | 1218 | 1092 | K | S | charge | Bening | 0.325 | Damaging | 0.01 | |
| *SIK2* | Salt inducible kinase 2 | 522 | 522 | L | V | not_radical | Bening | 0.005 | Damaging | 0.02 | |
| | | 543 | 543 | H | Q | charge | Bening | 0.009 | Damaging | 0.00 | |
| | | 900 | 897 | R | P | charge | Possibly damaging | 0.454 | Damaging | 0.00 | |
| *SIK3* | Salt inducible kinase 3 | 498 | 408 | Q | P | polarity | Probably damaging | 0.971 | Damaging | 0.00 | |
| | | 953 | 863 | N | S | polarity&volume | Bening | 0.004 | Damaging | 0.00 | |
| *TIMELESS* | Timeless circadian regulator | 35 | 35 | I | V | not_radical | Bening | 0.079 | Tolerated | 0.10 | Required for homodimerization and for interaction with CRY1 and CHEK1 |
| | | 58 | 58 | T | A | polarity | Possibly damaging | 0.931 | Tolerated | 0.13 | Required for homodimerization and for interaction with CRY1 and CHEK1 |

| | | | | | | | | | |
|---|---|---|---|---|---|---|---|---|---|
| 232 | 232 | A | E | charge | Probably damaging | 1 | Tolerated | 0.07 | Required for homodimerization and for interaction with CRY1 and CHEK1 |
| 258 | 258 | H | Q | charge | Probably damaging | 1 | Damaging | 0.01 | Required for homodimerization and for interaction with CRY1 and CHEK1 |
| 331 | 331 | E | Q | charge | Bening | 0.011 | Tolerated | 0.34 | |
| 450 | 450 | L | V | not_radical | Bening | 0.025 | Tolerated | 0.06 | |
| 565 | 564 | T | A | polarity | Bening | 0.265 | Damaging | 0.00 | |
| 889 | 885 | D | H | charge | Bening | 0.014 | Tolerated | 0.09 | |
| 975 | 971 | I | L | not_radical | Bening | 0.418 | Tolerated | 0.36 | |
| 977 | 972 | K | E | charge | Bening | 0.081 | Damaging | 0.00 | |
| 1099 | 1090 | Y | Q | polarity&volume | Bening | 0.387 | Damaging | 0.00 | Interaction with PARP1 |
| 1113 | 1102 | R | E | charge | Bening | 0.005 | Damaging | 0.00 | |

**Table S12 | Statistics of dugong whole-genome resequencing samples**

QLD denotes Queensland, Australia; NSW, New South Wales, Australia; WA, Western Australia, Australia. Samples SRR17870680, ERR5621402, and DRR251525 were obtained from NCBI SRA. The Rx ratio denotes the ratio of sequence reads that align to the X chromosome to reads that align to autosomal chromosomes and allows for sex determination of a sample (ca. 1.0 for females and 0.5 for males). Rx.CI denotes the Rx ratio 95% Confidence Interval upper and lower bound. Clean reads refer to the number of reads processed (removal of adaptors sequences and low-quality base calls) using *SOAPnuke* and *Flexbar*; mapping rate, percentage of reads that mapped to the dugong genome. Sequencing depth (average coverage across the genome) was calculated using *mosdepth* after downstream BAM file filtering to remove PCR duplicates and reads with a quality score below 30 (Q30). For comparative analyses, all samples were to downsampled to ~10× coverage.

| Sample ID | Sample location | Rx ratio | Rx.CI | Predicted sex | Clean reads | Mapping rate (%) | Sequencing depth (×) |
|---|---|---|---|---|---|---|---|
| SRR17870680 | Exmouth Gulf, WA | 0.94 | 0.94-0.95 | F | 649,073,882 | 98.03 | 32.01 |
| ERR5621402 | Coogee Beach, NSW | 0.55 | 0.54-0.55 | M | 719,917,332 | 97.80 | 35.50 |
| DRR251525 | Okinawa, Japan | 0.90 | 0.89-0.92 | F | 1,097,858,936 | 98.24 | 54.14 |
| AB16008 | Airlie Beach, QLD | 0.93 | 0.92-0.94 | F | 202,766,042 | 99.3% | 9.93 |
| AB16009 | Airlie Beach, QLD | 0.93 | 0.92-0.93 | F | 211,553,210 | 99.3% | 10.49 |
| AB16010 | Airlie Beach, QLD | 0.49 | 0.49-0.50 | M | 220,082,820 | 99.4% | 10.59 |
| BG15013 | Bowling Green Bay, QLD | 0.93 | 0.92-0.94 | F | 245,198,972 | 99.4% | 12.09 |
| BG15014 | Bowling Green Bay, QLD | 0.93 | 0.93-0.94 | F | 237,077,906 | 98.9% | 11.66 |
| BG15018 | Bowling Green Bay, QLD | 0.95 | 0.94-0.95 | F | 208,241,464 | 99.4% | 10.27 |
| CV16040 | Clairview, QLD | 0.94 | 0.94-0.95 | F | 252,128,106 | 97.9% | 12.31 |
| CV16059 | Clairview, QLD | 0.94 | 0.94-0.95 | F | 204,401,232 | 98.8% | 10.00 |
| CV16060 | Clairview, QLD | 0.93 | 0.93-0.94 | F | 219,502,504 | 98.9% | 10.73 |
| CV16061 | Clairview, QLD | 0.93 | 0.92-0.93 | F | 211,665,556 | 97.6% | 10.24 |
| CV16062 | Clairview, QLD | 0.93 | 0.93-0.94 | F | 254,118,810 | 96.9% | 12.19 |
| CV16063 | Clairview, QLD | 0.93 | 0.93-0.94 | F | 203,976,830 | 98.5% | 9.93 |
| CV16064 | Clairview, QLD | 0.94 | 0.93-0.94 | F | 209,503,792 | 96.8% | 9.96 |
| CV16065 | Clairview, QLD | 0.92 | 0.89-0.94 | F | 198,008,252 | 99.5% | 9.71 |
| CV16067 | Clairview, QLD | 0.93 | 0.92-0.93 | F | 210,596,544 | 99.0% | 10.51 |
| D110423 | Torres Strait, QLD | 0.50 | 0.49-0.50 | M | 230,377,224 | 99.4% | 11.02 |

| | | | | | | | |
|---|---|---|---|---|---|---|---|
| D110424 | Torres Strait, QLD | 0.48 | 0.48-0.49 | M | 232,543,368 | 99.4% | 11.20 |
| D110425A | Torres Strait, QLD | 0.48 | 0.47-0.48 | M | 212,677,440 | 99.4% | 10.20 |
| D110425B | Torres Strait, QLD | 0.48 | 0.48-0.49 | M | 216,264,460 | 99.4% | 10.41 |
| D110426 | Torres Strait, QLD | 0.49 | 0.49-0.50 | M | 179,459,920 | 99.3% | 8.45 |
| GS10410 | Great Sandy Strait, QLD | 0.96 | 0.96-0.97 | F | 186,188,228 | 98.4% | 9.35 |
| GS10436 | Great Sandy Strait, QLD | 0.96 | 0.95-0.96 | F | 265,346,328 | 99.5% | 13.56 |
| GS10437 | Great Sandy Strait, QLD | 0.50 | 0.49-0.50 | M | 257,324,238 | 99.4% | 12.77 |
| GS10438 | Great Sandy Strait, QLD | 0.96 | 0.95-0.96 | F | 267,974,152 | 87.6% | 11.38 |
| GS10440 | Great Sandy Strait, QLD | 0.51 | 0.51-0.51 | M | 184,083,944 | 94.9% | 8.61 |
| GS10463 | Great Sandy Strait, QLD | 0.97 | 0.96-0.97 | F | 210,175,768 | 98.1% | 10.56 |
| GS10464 | Great Sandy Strait, QLD | 0.51 | 0.51-0.51 | M | 265,861,832 | 96.6% | 12.77 |
| GS10465 | Great Sandy Strait, QLD | 0.96 | 0.95-0.96 | F | 230,176,318 | 99.5% | 11.73 |
| GS10466 | Great Sandy Strait, QLD | 0.50 | 0.50-0.50 | M | 251,255,394 | 94.9% | 11.69 |
| GS10467 | Great Sandy Strait, QLD | 0.96 | 0.95-0.96 | F | 227,477,698 | 95.8% | 11.04 |
| GS10468 | Great Sandy Strait, QLD | 0.97 | 0.96-0.97 | F | 255,045,206 | 99.6% | 13.06 |
| GS10469 | Great Sandy Strait, QLD | 0.96 | 0.95-0.96 | F | 228,084,294 | 98.5% | 11.46 |
| GS10470 | Great Sandy Strait, QLD | 0.96 | 0.96-0.97 | F | 225,496,236 | 99.4% | 11.50 |
| GS10487 | Great Sandy Strait, QLD | 0.50 | 0.50-0.51 | M | 226,006,358 | 99.4% | 11.22 |
| GS10489 | Great Sandy Strait, QLD | 0.95 | 0.95-0.95 | F | 225,252,460 | 98.7% | 11.29 |
| GS11471 | Great Sandy Strait, QLD | 0.94 | 0.93-0.94 | F | 261,425,544 | 98.0% | 13.07 |
| GS11472 | Great Sandy Strait, QLD | 0.96 | 0.95-0.96 | F | 226,343,068 | 96.4% | 11.08 |
| GS11474 | Great Sandy Strait, QLD | 0.49 | 0.49-0.50 | M | 209,234,486 | 98.7% | 10.31 |
| GS11475 | Great Sandy Strait, QLD | 0.94 | 0.94-0.95 | F | 209,032,864 | 96.2% | 10.22 |
| GS11476 | Great Sandy Strait, QLD | 0.95 | 0.95-0.96 | F | 207,806,604 | 95.2% | 10.02 |
| GS11477 | Great Sandy Strait, QLD | 0.96 | 0.95-0.96 | F | 195,442,134 | 99.4% | 9.95 |
| GS11478 | Great Sandy Strait, QLD | 0.92 | 0.91-0.92 | F | 227,740,852 | 93.0% | 10.74 |
| GS11480 | Great Sandy Strait, QLD | 0.95 | 0.95-0.96 | F | 212,008,656 | 96.0% | 10.31 |
| HB10001 | Hervey Bay, QLD | 0.94 | 0.93-0.94 | F | 238,126,576 | 90.1% | 10.49 |

| | | | | | | | |
|---|---|---|---|---|---|---|---|
| HB10003 | Hervey Bay, QLD | 0.95 | 0.94-0.95 | F | 228,354,142 | 99.4% | 11.42 |
| HB10004 | Hervey Bay, QLD | 0.96 | 0.95-0.96 | F | 218,644,040 | 99.4% | 11.09 |
| HB10005 | Hervey Bay, QLD | 0.51 | 0.50-0.51 | M | 241,754,670 | 99.2% | 11.96 |
| HB10006 | Hervey Bay, QLD | 0.96 | 0.95-0.96 | F | 230,969,098 | 97.9% | 11.48 |
| HB10007 | Hervey Bay, QLD | 0.96 | 0.95-0.96 | F | 220,027,734 | 91.6% | 9.94 |
| HB10008 | Hervey Bay, QLD | 0.96 | 0.95-0.96 | F | 229,066,940 | 99.4% | 11.62 |
| HB10009 | Hervey Bay, QLD | 0.94 | 0.94-0.95 | F | 231,837,962 | 97.7% | 11.46 |
| HB10010 | Hervey Bay, QLD | 0.96 | 0.96-0.96 | F | 258,289,108 | 99.4% | 13.14 |
| HB10012 | Hervey Bay, QLD | 0.91 | 0.90-0.92 | F | 199,860,132 | 97.1% | 9.82 |
| HB10013 | Hervey Bay, QLD | 0.96 | 0.96-0.97 | F | 227,344,364 | 99.3% | 11.61 |
| HB11001 | Hervey Bay, QLD | 0.93 | 0.93-0.94 | F | 222,557,932 | 99.4% | 11.14 |
| HB11002 | Hervey Bay, QLD | 0.95 | 0.94-0.95 | F | 233,270,642 | 98.7% | 11.54 |
| HB11003 | Hervey Bay, QLD | 0.50 | 0.50-0.50 | M | 231,036,470 | 99.4% | 11.32 |
| HB11004 | Hervey Bay, QLD | 0.94 | 0.94-0.94 | F | 220,232,312 | 98.5% | 10.87 |
| HB11005 | Hervey Bay, QLD | 0.50 | 0.50-0.50 | M | 252,922,980 | 99.5% | 12.32 |
| HB11006 | Hervey Bay, QLD | 0.96 | 0.95-0.97 | F | 176,039,116 | 99.3% | 8.75 |
| HB11069 | Hervey Bay, QLD | 0.95 | 0.94-0.95 | F | 219,630,820 | 98.9% | 10.77 |
| HB11070 | Hervey Bay, QLD | 0.93 | 0.92-0.93 | F | 215,045,726 | 99.0% | 10.66 |
| HB11075 | Hervey Bay, QLD | 0.94 | 0.93-0.94 | F | 269,837,680 | 99.0% | 13.35 |
| HB11079 | Hervey Bay, QLD | 0.96 | 0.95-0.96 | F | 233,622,522 | 99.5% | 11.67 |
| HB11080 | Hervey Bay, QLD | 0.50 | 0.50-0.50 | M | 207,213,160 | 99.4% | 10.10 |
| HB11081 | Hervey Bay, QLD | 0.50 | 0.50-0.50 | M | 202,021,616 | 99.1% | 9.83 |
| HB11082 | Hervey Bay, QLD | 0.50 | 0.50-0.51 | M | 104,771,802 | 99.3% | 10.65 |
| MB10044 | Moreton Bay, QLD | 0.50 | 0.49-0.50 | M | 245,596,948 | 96.2% | 11.61 |
| MB10045 | Moreton Bay, QLD | 0.48 | 0.48-0.49 | M | 238,967,806 | 99.5% | 11.87 |
| MB10046 | Moreton Bay, QLD | 0.49 | 0.49-0.49 | M | 230,176,802 | 99.5% | 11.41 |
| MB10047 | Moreton Bay, QLD | 0.49 | 0.49-0.49 | M | 212,588,546 | 98.2% | 10.34 |
| MB10048 | Moreton Bay, QLD | 0.49 | 0.49-0.49 | M | 197,221,478 | 99.5% | 9.76 |

| | | | | | | | |
|---|---|---|---|---|---|---|---|
| MB10049 | Moreton Bay, QLD | 0.50 | 0.49-0.50 | M | 207,209,880 | 99.5% | 10.27 |
| MB10051 | Moreton Bay, QLD | 0.49 | 0.49-0.50 | M | 204,820,090 | 99.2% | 10.10 |
| MB10052 | Moreton Bay, QLD | 0.93 | 0.92-0.93 | F | 199,767,594 | 98.2% | 9.96 |
| MB10056 | Moreton Bay, QLD | 0.50 | 0.50-0.50 | M | 119,236,684 | 98.9% | 12.19 |
| MB10057 | Moreton Bay, QLD | 0.94 | 0.94-0.95 | F | 245,450,126 | 98.9% | 12.33 |
| MB10058 | Moreton Bay, QLD | 0.50 | 0.50-0.50 | M | 260,396,088 | 75.0% | 8.83 |
| MB10059 | Moreton Bay, QLD | 0.95 | 0.95-0.96 | F | 241,494,606 | 98.0% | 11.94 |
| MB12352 | Moreton Bay, QLD | 0.95 | 0.95-0.96 | F | 261,069,912 | 99.1% | 13.13 |
| MB12355 | Moreton Bay, QLD | 0.95 | 0.94-0.95 | F | 267,221,292 | 99.2% | 13.47 |
| MB12357 | Moreton Bay, QLD | 0.50 | 0.50-0.50 | M | 247,009,096 | 99.4% | 12.20 |
| MB14552 | Moreton Bay, QLD | 0.95 | 0.94-0.95 | F | 240,761,136 | 99.4% | 12.11 |
| MB14553 | Moreton Bay, QLD | 0.50 | 0.50-0.50 | M | 261,111,958 | 98.7% | 12.76 |
| MB14554 | Moreton Bay, QLD | 0.93 | 0.93-0.94 | F | 251,272,698 | 99.2% | 12.59 |
| MB14555 | Moreton Bay, QLD | 0.95 | 0.95-0.96 | F | 240,531,822 | 99.4% | 12.15 |
| MB14557 | Moreton Bay, QLD | 0.51 | 0.50-0.51 | M | 237,337,772 | 97.8% | 11.62 |
| MB14560 | Moreton Bay, QLD | 0.50 | 0.49-0.50 | M | 231,925,566 | 98.5% | 11.45 |
| MB14561 | Moreton Bay, QLD | 0.51 | 0.50-0.51 | M | 229,378,714 | 99.4% | 11.50 |
| MB14562 | Moreton Bay, QLD | 0.51 | 0.50-0.51 | M | 266,777,668 | 99.0% | 13.28 |
| MB14563 | Moreton Bay, QLD | 0.50 | 0.50-0.51 | M | 251,336,516 | 95.8% | 12.00 |
| MB14564 | Moreton Bay, QLD | 0.50 | 0.50-0.50 | M | 237,737,708 | 99.5% | 11.88 |
| MB14565 | Moreton Bay, QLD | 0.94 | 0.94-0.95 | F | 713,408,396 | 99.1% | 35.83 |
| MB16789 | Moreton Bay, QLD | 0.95 | 0.94-0.95 | F | 136,559,610 | 99.4% | 6.91 |
| MB16796 | Moreton Bay, QLD | 0.95 | 0.94-0.95 | F | 219,995,112 | 99.4% | 11.08 |
| MB18030 | Moreton Bay, QLD | 0.95 | 0.94-0.96 | F | 249,004,940 | 98.3% | 12.39 |
| MB18041 | Moreton Bay, QLD | 0.51 | 0.50-0.51 | M | 252,932,710 | 99.3% | 12.57 |
| MB20195 | Moreton Bay, QLD | 0.95 | 0.94-0.95 | F | 232,288,232 | 99.4% | 11.77 |
| MB20198 | Moreton Bay, QLD | 0.95 | 0.95-0.96 | F | 236,899,694 | 99.3% | 12.04 |

**Table S13 | List of genes under selective sweep between dugong populations north and south of the Whitsundays Islands.**

Genes were identified using XP-EHH 100, XP-CLR 101, π, and $F_{st}$). Gene models with an XP prefix refers to NCBI GenBank annotations; evm.model, initial dugong gene models; homol.model, gene models obtained by adding additional homolog protein data.

| Chromosome | Start | End | Gene model | Description |
|---|---|---|---|---|
| **18** | 24,549,953 | 24,551,831 | XP_003418414 | CLPX caseinolytic mitochondrial matrix peptidase chaperone subunit X |
| **18** | 25,202,082 | 25,539,778 | XP_007456442 | putative V-set and immunoglobulin domain-containing-like protein IGHV4OR15-8-like |
| **18** | 25,333,034 | 25,333,399 | evm.model.chr19.127 | K06856 immunoglobulin heavy chain |
| **18** | 25,521,103 | 25,522,971 | XP_004409458 | CLPX; ATP-dependent Clp protease ATP-binding subunit clpX-like, mitochondrial precursor |
| **18** | 25,583,962 | 25,585,326 | homol.model_1203 | NUP42; nucleoporin NUP42 isoform X2 |
| **18** | 26,350,303 | 26,411,351 | evm.model.chr19.133 | K06856 immunoglobulin heavy chain |

**Table S14 | Runs of homozygosity (ROHs) statistics of dugong whole-genome resequencing samples**

QLD denotes Queensland, Australia; NSW, New South Wales, Australia; WA, Western Australia, Australia. Samples SRR17870680 (Exmouth Gulf, WA), ERR5621402 (Coogee Beach, NSW), and DRR251525 (Okinawa, Japan) were obtained from NCBI SRA.

| Sample | ROH>100kb | | | ROH>1Mb | | | ROH>5Mb | | | ROH>10Mb | | |
|---|---|---|---|---|---|---|---|---|---|---|---|---|
| | ROH Number | ROH Length (bp) | $F_{ROH}$ | ROH Number | ROH Length (bp) | $F_{ROH}$ | ROH Number | ROH Length (bp) | $F_{ROH}$ | ROH Number | ROH Length (bp) | $F_{ROH}$ |
| SRR17870680 | 2,723 | 1,275,993,905 | 0.440773 | 169 | 329,629,866 | 0.113866 | 9 | 60,073,762 | 0.020752 | 0 | 0 | 0.000000 |
| DRR251525 | 2,492 | 1,762,094,597 | 0.608690 | 389 | 934,384,760 | 0.322769 | 29 | 236,330,529 | 0.081637 | 4 | 73,537,394 | 0.025402 |
| ERR5621402 | 2,543 | 1,347,913,879 | 0.465617 | 277 | 478,941,095 | 0.165443 | 7 | 48,669,533 | 0.016812 | 0 | 0 | 0.000000 |
| AB16008 | 3,436 | 919,448,002 | 0.317610 | 36 | 44,755,991 | 0.015460 | 0 | 0 | 0.000000 | 0 | 0 | 0.000000 |
| AB16009 | 1,917 | 415,328,546 | 0.143469 | 5 | 5,799,069 | 0.002003 | 0 | 0 | 0.000000 | 0 | 0 | 0.000000 |
| AB16010 | 1,939 | 441,506,904 | 0.152512 | 15 | 20,487,269 | 0.007077 | 0 | 0 | 0.000000 | 0 | 0 | 0.000000 |
| BG15013 | 1,762 | 352,488,010 | 0.121762 | 1 | 1,464,967 | 0.000506 | 0 | 0 | 0.000000 | 0 | 0 | 0.000000 |
| BG15014 | 1,825 | 378,976,675 | 0.130912 | 5 | 7,542,872 | 0.002606 | 0 | 0 | 0.000000 | 0 | 0 | 0.000000 |
| BG15018 | 1,880 | 389,466,369 | 0.134535 | 4 | 6,122,416 | 0.002115 | 0 | 0 | 0.000000 | 0 | 0 | 0.000000 |
| CV16040 | 2,816 | 608,596,711 | 0.210231 | 1 | 1,065,280 | 0.000368 | 0 | 0 | 0.000000 | 0 | 0 | 0.000000 |
| CV16059 | 2,674 | 566,331,829 | 0.195631 | 5 | 5,739,898 | 0.001983 | 0 | 0 | 0.000000 | 0 | 0 | 0.000000 |
| CV16060 | 2,627 | 627,124,822 | 0.216631 | 17 | 20,491,817 | 0.007079 | 0 | 0 | 0.000000 | 0 | 0 | 0.000000 |
| CV16061 | 2,615 | 587,579,352 | 0.202971 | 7 | 7,834,250 | 0.002706 | 0 | 0 | 0.000000 | 0 | 0 | 0.000000 |
| CV16062 | 2,985 | 660,201,172 | 0.228057 | 8 | 10,057,355 | 0.003474 | 0 | 0 | 0.000000 | 0 | 0 | 0.000000 |
| CV16063 | 2,572 | 553,315,064 | 0.191135 | 3 | 3,702,150 | 0.001279 | 0 | 0 | 0.000000 | 0 | 0 | 0.000000 |
| CV16064 | 2,853 | 664,977,369 | 0.229707 | 13 | 16,754,019 | 0.005787 | 0 | 0 | 0.000000 | 0 | 0 | 0.000000 |
| CV16065 | 2,128 | 328,321,674 | 0.113414 | 0 | 0 | 0.000000 | 0 | 0 | 0.000000 | 0 | 0 | 0.000000 |
| CV16067 | 2,649 | 580,060,843 | 0.200373 | 8 | 8,739,312 | 0.003019 | 0 | 0 | 0.000000 | 0 | 0 | 0.000000 |
| D110423 | 1,689 | 333,359,816 | 0.115154 | 7 | 8,141,995 | 0.002813 | 0 | 0 | 0.000000 | 0 | 0 | 0.000000 |
| D110424 | 1,687 | 337,138,941 | 0.116460 | 2 | 2,335,035 | 0.000807 | 0 | 0 | 0.000000 | 0 | 0 | 0.000000 |
| D110425A | 1,754 | 370,313,260 | 0.127919 | 2 | 2,514,108 | 0.000868 | 0 | 0 | 0.000000 | 0 | 0 | 0.000000 |
| D110425B | 1,744 | 340,389,584 | 0.117583 | 2 | 2,479,494 | 0.000857 | 0 | 0 | 0.000000 | 0 | 0 | 0.000000 |
| D110426 | 1,648 | 352,381,696 | 0.121725 | 6 | 7,495,621 | 0.002589 | 0 | 0 | 0.000000 | 0 | 0 | 0.000000 |

| | | | | | | | | | | | | |
|---|---|---|---|---|---|---|---|---|---|---|---|---|
| GS10410 | 3,287 | 675,783,692 | 0.233440 | 2 | 2,245,470 | 0.000776 | 0 | 0 | 0.000000 | 0 | 0 | 0.000000 |
| GS10436 | 3,390 | 736,370,539 | 0.254368 | 8 | 8,847,247 | 0.003056 | 0 | 0 | 0.000000 | 0 | 0 | 0.000000 |
| GS10437 | 3,131 | 750,425,092 | 0.259223 | 18 | 21,751,183 | 0.007514 | 0 | 0 | 0.000000 | 0 | 0 | 0.000000 |
| GS10438 | 3,221 | 655,755,474 | 0.226521 | 3 | 3,344,753 | 0.001155 | 0 | 0 | 0.000000 | 0 | 0 | 0.000000 |
| GS10440 | 3,198 | 732,200,345 | 0.252928 | 6 | 7,271,252 | 0.002512 | 0 | 0 | 0.000000 | 0 | 0 | 0.000000 |
| GS10463 | 3,349 | 701,147,638 | 0.242201 | 6 | 7,574,322 | 0.002616 | 0 | 0 | 0.000000 | 0 | 0 | 0.000000 |
| GS10464 | 3,441 | 740,953,455 | 0.255951 | 4 | 4,803,374 | 0.001659 | 0 | 0 | 0.000000 | 0 | 0 | 0.000000 |
| GS10465 | 3,065 | 673,362,600 | 0.232603 | 11 | 13,385,748 | 0.004624 | 0 | 0 | 0.000000 | 0 | 0 | 0.000000 |
| GS10466 | 3,192 | 687,689,355 | 0.237552 | 1 | 1,111,612 | 0.000384 | 0 | 0 | 0.000000 | 0 | 0 | 0.000000 |
| GS10467 | 3,269 | 677,732,820 | 0.234113 | 6 | 8,423,935 | 0.002910 | 0 | 0 | 0.000000 | 0 | 0 | 0.000000 |
| GS10468 | 3,411 | 703,832,927 | 0.243129 | 7 | 7,718,826 | 0.002666 | 0 | 0 | 0.000000 | 0 | 0 | 0.000000 |
| GS10469 | 3,752 | 768,880,016 | 0.265598 | 5 | 5,331,942 | 0.001842 | 0 | 0 | 0.000000 | 0 | 0 | 0.000000 |
| GS10470 | 3,071 | 654,227,163 | 0.225993 | 7 | 7,815,861 | 0.002700 | 0 | 0 | 0.000000 | 0 | 0 | 0.000000 |
| GS10487 | 3,278 | 693,144,172 | 0.239436 | 5 | 5,778,609 | 0.001996 | 0 | 0 | 0.000000 | 0 | 0 | 0.000000 |
| GS10489 | 3,411 | 708,285,721 | 0.244667 | 5 | 5,396,755 | 0.001864 | 0 | 0 | 0.000000 | 0 | 0 | 0.000000 |
| GS11471 | 3,246 | 697,532,823 | 0.240952 | 2 | 2,385,956 | 0.000824 | 0 | 0 | 0.000000 | 0 | 0 | 0.000000 |
| GS11472 | 3,396 | 744,959,515 | 0.257335 | 7 | 10,441,777 | 0.003607 | 0 | 0 | 0.000000 | 0 | 0 | 0.000000 |
| GS11474 | 3,277 | 745,958,495 | 0.257680 | 7 | 8,274,897 | 0.002858 | 0 | 0 | 0.000000 | 0 | 0 | 0.000000 |
| GS11475 | 3,264 | 739,534,187 | 0.255461 | 9 | 10,458,527 | 0.003613 | 0 | 0 | 0.000000 | 0 | 0 | 0.000000 |
| GS11476 | 3,244 | 745,042,136 | 0.257364 | 9 | 11,342,808 | 0.003918 | 0 | 0 | 0.000000 | 0 | 0 | 0.000000 |
| GS11477 | 2,986 | 634,049,383 | 0.219023 | 7 | 7,706,058 | 0.002662 | 0 | 0 | 0.000000 | 0 | 0 | 0.000000 |
| GS11478 | 3,211 | 780,243,201 | 0.269524 | 24 | 29,016,538 | 0.010023 | 0 | 0 | 0.000000 | 0 | 0 | 0.000000 |
| GS11480 | 3,403 | 759,768,120 | 0.262451 | 9 | 11,170,571 | 0.003859 | 0 | 0 | 0.000000 | 0 | 0 | 0.000000 |
| HB10001 | 3,025 | 606,188,508 | 0.209399 | 4 | 4,673,636 | 0.001614 | 0 | 0 | 0.000000 | 0 | 0 | 0.000000 |
| HB10003 | 3,281 | 717,080,793 | 0.247705 | 6 | 7,512,993 | 0.002595 | 0 | 0 | 0.000000 | 0 | 0 | 0.000000 |
| HB10004 | 3,054 | 622,374,828 | 0.214990 | 2 | 2,211,634 | 0.000764 | 0 | 0 | 0.000000 | 0 | 0 | 0.000000 |
| HB10005 | 3,058 | 659,080,815 | 0.227670 | 8 | 9,030,624 | 0.003119 | 0 | 0 | 0.000000 | 0 | 0 | 0.000000 |
| HB10006 | 3,268 | 768,845,796 | 0.265586 | 17 | 19,551,327 | 0.006754 | 0 | 0 | 0.000000 | 0 | 0 | 0.000000 |
| HB10007 | 2,967 | 535,403,032 | 0.184947 | 0 | 0 | 0.000000 | 0 | 0 | 0.000000 | 0 | 0 | 0.000000 |

| | | | | | | | | | | | | |
|---|---|---|---|---|---|---|---|---|---|---|---|---|
| HB10008 | 3,295 | 688,090,632 | 0.237691 | 6 | 7,710,958 | 0.002664 | 0 | 0 | 0.000000 | 0 | 0 | 0.000000 |
| HB10009 | 3,184 | 747,128,629 | 0.258085 | 12 | 14,077,590 | 0.004863 | 0 | 0 | 0.000000 | 0 | 0 | 0.000000 |
| HB10010 | 3,187 | 721,676,600 | 0.249293 | 6 | 6,836,740 | 0.002362 | 0 | 0 | 0.000000 | 0 | 0 | 0.000000 |
| HB10012 | 3,382 | 747,505,861 | 0.258215 | 5 | 6,622,451 | 0.002288 | 0 | 0 | 0.000000 | 0 | 0 | 0.000000 |
| HB10013 | 3,070 | 633,106,638 | 0.218697 | 3 | 3,072,054 | 0.001061 | 0 | 0 | 0.000000 | 0 | 0 | 0.000000 |
| HB11001 | 3,121 | 679,347,437 | 0.234671 | 8 | 9,114,191 | 0.003148 | 0 | 0 | 0.000000 | 0 | 0 | 0.000000 |
| HB11002 | 3,193 | 687,452,009 | 0.237470 | 4 | 4,728,654 | 0.001633 | 0 | 0 | 0.000000 | 0 | 0 | 0.000000 |
| HB11003 | 3,130 | 631,068,415 | 0.217993 | 4 | 4,361,227 | 0.001507 | 0 | 0 | 0.000000 | 0 | 0 | 0.000000 |
| HB11004 | 3,339 | 695,211,679 | 0.240151 | 8 | 10,077,879 | 0.003481 | 0 | 0 | 0.000000 | 0 | 0 | 0.000000 |
| HB11005 | 3,131 | 660,876,141 | 0.228290 | 3 | 3,290,559 | 0.001137 | 0 | 0 | 0.000000 | 0 | 0 | 0.000000 |
| HB11006 | 3,135 | 697,686,446 | 0.241005 | 5 | 5,451,205 | 0.001883 | 0 | 0 | 0.000000 | 0 | 0 | 0.000000 |
| HB11069 | 3,379 | 758,188,208 | 0.261905 | 8 | 9,498,572 | 0.003281 | 0 | 0 | 0.000000 | 0 | 0 | 0.000000 |
| HB11070 | 3,129 | 692,655,676 | 0.239268 | 7 | 8,743,938 | 0.003020 | 0 | 0 | 0.000000 | 0 | 0 | 0.000000 |
| HB11075 | 3,128 | 646,414,519 | 0.223294 | 1 | 1,348,323 | 0.000466 | 0 | 0 | 0.000000 | 0 | 0 | 0.000000 |
| HB11079 | 2,907 | 527,719,883 | 0.182293 | 0 | 0 | 0.000000 | 0 | 0 | 0.000000 | 0 | 0 | 0.000000 |
| HB11080 | 3,104 | 635,114,324 | 0.219391 | 0 | 0 | 0.000000 | 0 | 0 | 0.000000 | 0 | 0 | 0.000000 |
| HB11081 | 3,425 | 796,444,196 | 0.275120 | 8 | 9,490,906 | 0.003278 | 0 | 0 | 0.000000 | 0 | 0 | 0.000000 |
| HB11082 | 2,146 | 338,310,888 | 0.116865 | 0 | 0 | 0.000000 | 0 | 0 | 0.000000 | 0 | 0 | 0.000000 |
| MB10044 | 3,437 | 680,555,080 | 0.235088 | 5 | 5,604,820 | 0.001936 | 0 | 0 | 0.000000 | 0 | 0 | 0.000000 |
| MB10045 | 3,349 | 739,424,772 | 0.255423 | 2 | 3,084,987 | 0.001066 | 0 | 0 | 0.000000 | 0 | 0 | 0.000000 |
| MB10046 | 3,491 | 815,942,282 | 0.281855 | 15 | 19,316,146 | 0.006672 | 0 | 0 | 0.000000 | 0 | 0 | 0.000000 |
| MB10047 | 3,456 | 749,094,488 | 0.258764 | 6 | 6,529,283 | 0.002255 | 0 | 0 | 0.000000 | 0 | 0 | 0.000000 |
| MB10048 | 3,338 | 726,664,627 | 0.251016 | 3 | 3,197,282 | 0.001104 | 0 | 0 | 0.000000 | 0 | 0 | 0.000000 |
| MB10049 | 3,278 | 672,900,527 | 0.232444 | 2 | 2,286,791 | 0.000790 | 0 | 0 | 0.000000 | 0 | 0 | 0.000000 |
| MB10051 | 3,743 | 837,487,213 | 0.289298 | 5 | 6,318,270 | 0.002183 | 0 | 0 | 0.000000 | 0 | 0 | 0.000000 |
| MB10052 | 3,533 | 734,620,591 | 0.253764 | 3 | 3,548,875 | 0.001226 | 0 | 0 | 0.000000 | 0 | 0 | 0.000000 |
| MB10056 | 2,641 | 426,288,169 | 0.147255 | 0 | 0 | 0.000000 | 0 | 0 | 0.000000 | 0 | 0 | 0.000000 |
| MB10057 | 3,539 | 782,089,437 | 0.270161 | 8 | 9,941,440 | 0.003434 | 0 | 0 | 0.000000 | 0 | 0 | 0.000000 |
| MB10058 | 3,354 | 641,693,070 | 0.221663 | 1 | 1,073,652 | 0.000371 | 0 | 0 | 0.000000 | 0 | 0 | 0.000000 |

| | | | | | | | | | | | | |
|---|---|---|---|---|---|---|---|---|---|---|---|---|
| MB10059 | 3,203 | 681,847,124 | 0.235534 | 3 | 3,375,813 | 0.001166 | 0 | 0 | 0.000000 | 0 | 0 | 0.000000 |
| MB12352 | 3,398 | 739,044,872 | 0.255292 | 5 | 5,550,668 | 0.001917 | 0 | 0 | 0.000000 | 0 | 0 | 0.000000 |
| MB12355 | 3,054 | 711,227,181 | 0.245683 | 13 | 15,248,768 | 0.005267 | 0 | 0 | 0.000000 | 0 | 0 | 0.000000 |
| MB12357 | 3,250 | 768,660,658 | 0.265522 | 11 | 13,485,478 | 0.004658 | 0 | 0 | 0.000000 | 0 | 0 | 0.000000 |
| MB14552 | 3,798 | 757,825,049 | 0.261779 | 1 | 1,033,872 | 0.000357 | 0 | 0 | 0.000000 | 0 | 0 | 0.000000 |
| MB14553 | 3,389 | 748,111,668 | 0.258424 | 11 | 12,926,565 | 0.004465 | 0 | 0 | 0.000000 | 0 | 0 | 0.000000 |
| MB14554 | 3,282 | 651,267,648 | 0.224971 | 2 | 2,036,106 | 0.000703 | 0 | 0 | 0.000000 | 0 | 0 | 0.000000 |
| MB14555 | 3,599 | 752,534,905 | 0.259952 | 8 | 9,064,741 | 0.003131 | 0 | 0 | 0.000000 | 0 | 0 | 0.000000 |
| MB14557 | 3,185 | 595,532,849 | 0.205718 | 3 | 3,304,922 | 0.001142 | 0 | 0 | 0.000000 | 0 | 0 | 0.000000 |
| MB14560 | 3,152 | 704,975,392 | 0.243523 | 14 | 16,552,927 | 0.005718 | 0 | 0 | 0.000000 | 0 | 0 | 0.000000 |
| MB14561 | 3,268 | 690,865,800 | 0.238649 | 4 | 4,752,417 | 0.001642 | 0 | 0 | 0.000000 | 0 | 0 | 0.000000 |
| MB14562 | 3,481 | 679,616,056 | 0.234763 | 0 | 0 | 0.000000 | 0 | 0 | 0.000000 | 0 | 0 | 0.000000 |
| MB14563 | 3,173 | 644,299,662 | 0.222564 | 5 | 6,515,528 | 0.002251 | 0 | 0 | 0.000000 | 0 | 0 | 0.000000 |
| MB14564 | 3,540 | 768,097,312 | 0.265328 | 9 | 10,144,503 | 0.003504 | 0 | 0 | 0.000000 | 0 | 0 | 0.000000 |
| MB14565 | 3,334 | 693,580,817 | 0.239587 | 4 | 4,496,736 | 0.001553 | 0 | 0 | 0.000000 | 0 | 0 | 0.000000 |
| MB16789 | 3,315 | 678,511,505 | 0.234382 | 4 | 4,326,422 | 0.001494 | 0 | 0 | 0.000000 | 0 | 0 | 0.000000 |
| MB16796 | 604 | 99,815,497 | 0.034480 | 0 | 0 | 0.000000 | 0 | 0 | 0.000000 | 0 | 0 | 0.000000 |
| MB18030 | 3,024 | 693,336,047 | 0.239503 | 11 | 13,249,597 | 0.004577 | 0 | 0 | 0.000000 | 0 | 0 | 0.000000 |
| MB18041 | 3,247 | 725,442,049 | 0.250593 | 8 | 9,413,241 | 0.003252 | 0 | 0 | 0.000000 | 0 | 0 | 0.000000 |
| MB20195 | 3,349 | 783,224,879 | 0.270553 | 12 | 15,988,453 | 0.005523 | 0 | 0 | 0.000000 | 0 | 0 | 0.000000 |
| MB20198 | 3,457 | 741,359,239 | 0.256092 | 7 | 8,320,133 | 0.002874 | 0 | 0 | 0.000000 | 0 | 0 | 0.000000 |
